# Microbiomes attached to fresh perennial ryegrass- are temporally resilient and adapt to changing ecological niches

**DOI:** 10.1101/2020.11.17.386292

**Authors:** Sharon A. Huws, Joan. E. Edwards, Wanchang Lin, Francesco Rubino, Mark Alston, David Swarbreck, Shabhonam Caim, Pauline Rees Stevens, Justin Pachebat, Mi-Young Won, Linda B. Oyama, Christopher J. Creevey, Alison H. Kingston-Smith

## Abstract

Gut microbiomes, such as the rumen, greatly influence host nutrition due to their feed energy-harvesting capacity. We investigated temporal ecological interactions facilitating energy-harvesting at the fresh perennial ryegrass (PRG)-biofilm interface in the rumen using an *in sacco* approach and prokaryotic metatranscriptomic profiling. Network analysis identified two distinct sub-microbiomes primarily representing primary (≤4h) and secondary (≥4h) colonisation phases and the most transcriptionally active bacterial families (i.e *Fibrobacteriaceae, Selemondaceae* and *Methanobacteriaceae*) did not interact with either sub-microbiome, indicating non-cooperative behaviour. Conversely, *Prevotellaceae* had most transcriptional activity within the primary sub-microbiome (focussed on protein metabolism) and *Lachnospiraceae* within the secondary sub-microbiome (focussed on carbohydrate degradation). Putative keystone taxa, with low transcriptional activity, were identified within both sub-microbiomes, highlighting the important synergistic role of minor bacterial families, however, we hypothesise that they may be ‘cheating’ in order to capitalise on the energy-harvesting capacity of other microbes. In terms of chemical cues underlying transition from primary to secondary colonisation phases, we suggest that AI-2 based quorum sensing plays a role, based on LuxS gene expression data, coupled with changes in PRG chemistry. In summary, this study provides the first major insight into the complex temporal ecological interactions occurring at the plant-biofilm interface within the rumen.

## Introduction

Vertebrates play host to a complex gut microbiome, generally dominated by a few well-studied groups, but with a large ensemble of minor microbial species whose contribution are only recently starting to be revealed [1-4]. While it is generally accepted that distinct microbiomes exist in distal locations in the gut, it is not clear whether a single location could simultaneously play host to multiple sub-microbiomes with distinct temporal and/or functional roles. Neither do we fully understand the importance of ecological interactions in these environments. Elucidating these temporal niche-specialised interactions could drive the generation of new strategies for targeted manipulation of vertebrate gut microbiomes for therapeutic, productivity or environmental benefits.

The rumen is a case in point and is the largest compartment of the ruminant forestomach, housing a complex microbiome that has a major impact on host nutrition and health [5]. This anoxic microbial ecosystem has evolved to harvest energy from largely recalcitrant, complex plant carbohydrates [5-9]. Rumen microbes commonly exist as biofilms on feed particles consumed by the host [10-13] and microbes within biofilms have been shown to interact intimately and influence each other’s evolutionary fitness in many ecological niches [9, 14-16]. Indeed, whilst the rumen microbes generally function symbiotically, they also compete against each other for evolutionary advantage [9, 17].

Using *rrn* Operon-based approaches and microscopy, we have previously demonstrated that rumen bacteria and anaerobic fungi attach rapidly to PRG [6, 18, 19], with biofilms being evident within 5 mins [11]. Colonisation by bacteria thereafter is biphasic, with primary (≤4h) and secondary (≥4h) colonisation phases previously described based on metataxonomy [10, 20]. We hypothesise that these distinct temporal bacterial colonisation phases represent the interaction of multiple, coherent and yet temporally distinct sub-microbiomes. While these interactions are frequently categorised as positive (e.g. mutualism), neutral (i.e. resulting in no effect) or negative (e.g. competition) [21, 22], it is becoming clear that ecological interactions in highly complex environments can also result in combinations of positive, neutral and negative outcomes. For example, amensalism in which the actor experiences no benefit or detriment and the recipient experiences a negative outcome or commensalism when the converse is true [21]. Added to this, keystone taxa are likely to have an important role mediating mutually beneficial interactions which may be disproportionate to their population size and the loss of these taxa would have a profound impact upon the ecosystem [1]. Finally, cheating behaviour is also likely to exist in such complex microbiomes, in which little cooperation is displayed by the ‘cheater’ but they gain benefit from the mutualistic cooperation displayed by other microbes [23]. Interestingly, cheating behaviour may not only benefit the cheater, but can also help maintain biodiversity [24], therefore may play an important role in the stability or resilience of complex ecosystems like the gut of vertebrates.

In order to understand these temporal ecological interactions at the plant-microbiome interface in the rumen, we used prokaryotic metatranscriptomics and gene network analysis of the attached microbial community on fresh perennial ryegrass (PRG) incubated *in sacco* in the rumen over an 8 h period. For the first time, we provide in-depth, gene-expression based understanding of the ecological interactions governing temporal niche specialisation, cooperation and competition within the PRG-attached rumen microbiome. This knowledge is essential for understanding microbial drivers of energy harvesting capacity, which influence ruminant feed efficiency and emissions, as well as to facilitate breeding of improved PRG cultivars for ruminant livestock.

## Methods

### Growth and preparation of plant material for *in sacco* incubations

PRG (*Lolium perenne* cv. AberDart) was grown from seed in plastic seed trays (length 38cm x width 24cm x depth 5cm) filled with soil/compost (Levington’s General Purpose). Plants were kept in a greenhouse under natural light with additional illumination provided (minimum 8h photoperiod). Temperature was controlled (22/19°C day/night) and plants were watered twice a week. Plants were harvested after 6 weeks by cutting them (approx. 3cm above soil level) directly before use, and then further processing them into 1cm sections using scissors. Triplicate samples of harvested plant material were also snap frozen in dry ice and stored at - 80°C for profiling of plant epiphytic prokaryotes (0h samples).

### Ruminal *in sacco* incubations

Three mature, rumen-cannulated, non-lactating Holstein x Friesian cows were used for this experiment. The experiment was conducted with the authority of Licenses under the United Kingdom Animal Scientific Procedures Act, 1986 and managed according to the protocols approved by the Aberystwyth University Animal Welfare and Ethics Review. For 2 weeks prior to the experiment, cows were fed a diet of straw and grass silage *ad libitum* (∼6.5 kg dry matter day^-1^) and were permitted field grazing on a permanent ryegrass pasture for at least 4h/day. For the duration of the experiment animals were fed silage daily in two equal meals at 07:00 and 16:00 and had constant access to straw when not field grazing.

The nylon bag technique was used as described previously [25, 26]. Stitched nylon bags (10cm × 20cm) of 100µm^2^ pore size were filled with 15g (fresh weight) of the processed 1cm length PRG and sealed at all perimeters by heating (Impulse sealer, American Int, Nl Electric, USA). For each cow, 10 bags were then connected to a 55cm plastic coated flexible cable with lacing cords, and then placed in the rumen before being attached to the cap of the fistula. Bags were placed simultaneously in the rumen of each cow shortly after animals were offered their first silage meal of the day, and two bags were removed after 1, 2, 4, 6 and 8h of incubation then processed by washing with distilled water (500mL added to plant material within bags and bags gently squeezed thereafter) to remove loosely attached microbes followed by immediate freezing in dry ice and storage at -80°C until RNA extraction.

### RNA extraction

Frozen samples were ground to a fine powder under liquid nitrogen and then RNA was extracted using a hot phenol method [27]. Essentially, aquaphenol (10mL) was added to the ground sample and then the sample was incubated at 65°C for 1h. Tubes were inverted before chloroform was added (5mL). Tubes were centrifuged (5,000 x *g* for 30 mins at 20°C) and then the upper phase was removed, then the procedure was repeated by the addition of more chloroform (5mL) and centrifugation. Lithium chloride (2M final concentration) was then added, to remove any contaminating DNA, and samples stored overnight at 4°C. Samples were subsequently centrifuged (13,000 x *g* for 30 mins at 4°C) and the supernatant discarded, then the procedure was repeated to ensure all DNA was removed. Once the supernatant was discarded the pellet was resuspended in ice-cold 80% ethanol and centrifuged (13,000 x *g* for 15 mins at 4°C), this was repeated twice before the pellet was air dried and resuspended in molecular grade water. Absence of DNA in all samples was confirmed using 16S rDNA PCR using non-barcoded primers and subsequent agarose gel electrophoresis as described in Huws *et al*., (2016). Quality and quantity of retrieved RNA was assessed using the Experion automated electrophoresis system and a RNA StdSens Analysis kit (Bio-Rad Ltd., UK).

### rRNA removal and metatranscriptome sequencing

Prokaryotic mRNA was enriched in all samples by firstly removing the polyA fraction of the mRNA pool using a MicroPoly(A)Purist kit (Ambion) according to the manufacturer’s protocol. Eukaryotic 18S rRNA was then minimised using both the RiboMinus plant and eukaryote kits (Invitrogen, Carlsbad, USA) according to the manufacturer’s protocols. Finally, 16S rRNA was minimized using the Ribo-Zero rRNA removal kit for bacteria (Epicentre, Madison, USA) according to the manufacturer’s protocol. The resulting enriched prokaryotic mRNA was prepared for sequencing using the TruSeq stranded mRNA library prep kit (illumina, California, USA) following the manufacturer’s guidelines. Subsequently, library sequencing was completed using the illumina HiSeq 2500 (illumina, California, USA) and a 100bp paired end sequencing approach.

### Metatranscriptome assembly

A flow diagram showing the steps taken to prepare and analyse the sequences obtained is shown in Supplementary Figure 1. Essentially, the assembly was performed through a series of steps in order to reduce the complexity of the process. First, the quality of raw reads was evaluated with FastQC [28] version 0.10.1, and subsequently trimmed using Trimmomatic [29] by 9 base pairs at the 5’ end and 3 base pairs at the 3’ end, including remaining adapters. Following this step, the removal of any remaining rRNA contamination was performed using MGKit (script rRNA-protozoa.py; [30]) and a custom database of rRNA sequences built by retrieving (i) the RDP databases for archaea, bacteria and fungi [31], release 11_2 and (ii) rRNA genes from protozoan species from the EBI-ENA, This custom database was used with bowtie2 (2.1.0) [32] using the ‘--no-mixed --local --sensitive-local -N 1’ options, saving only the unaligned and hence non-rRNA reads (using the ‘--un-conc-gz’ option) to files. The retained reads were then aligned to the draft *Lolium perenne* genome [33] using bowtie and the same settings, retaining only those reads that did not align to the genome. Similarly, reads that aligned to a collection of 246 publicly available genomes, in particular from the Hungate1000 collection [34], were excluded from the remaining *de novo* assembling steps, using the aforementioned options (list of genomes available in Supplementary Excel 1). Genomes with reads aligned were used for downstream analysis, and were combined with the final assembly of reads that did not align to the genome collection or the rRNA database.

The first step of the meta-transcriptome assembly of non-aligned reads involved digital normalisation to reduce the bias from more abundant transcripts in the samples. The khmer package [35] was used in a two-step procedure. First the reads were normalised on a per-sample basis using the normalize-by-median.py script (with the options -p, -k 20, -N 4, -x 16e9, -C 20). Afterwards, a collective normalisation was carried out pooling all per-sample normalised reads. The same parameters as the per-sample normalisation were used, except for -x (an option to define the hash table size), which was set to 32e9 to account for the larger size of the input data. Following digital normalisation, an overall assembly was carried out using velvet version 1.2.10 [36] with a kmer size of 31 and automatic expected coverage (-exp_cov auto). Finally, alignments of all the reads used in the assembly to the final resulting assembly were created using bowtie2 (2.1.0) [32] using the ‘--no-mixed --local --sensitive-local -N 1’. The same procedure was carried out in parallel for the reads that aligned to the available prokaryotic genomes and the results of both were combined for each sample.

### CDS prediction and quantification

Putative coding sequences (CDS) were predicted in the de novo assembled transcriptome using the default options of the ‘TransDecoder.LongOrfs’ script from Transdecoder: https://github.com/jls943/TransDecoder [37, 38] which identifies ORFs that are at least 100 amino acids long. These were combined with the predicted CDS from the genomes that had aligned mRNA reads. These annotations formed the set of CDS which were used in the subsequent functional and taxonomic analyses. Abundance (counts) of reads mapping to each CDS annotation was calculated with htseq-count from the HTSeq package [39] (options: -s no; - t CDS; -q). The predicted amino acid (AA) sequences of these CDS were used in the subsequent annotation steps.

### Taxonomic Annotation

The blastp command of the BLAST package (10.1186/1471-2105-10-421) was used with the - outfmt 6 option against the NCBI nr database. The output was then passed to the add-gff-info script of the MGKit package, which uses a last common ancestor algorithm in the taxonomy command to resolve the taxon identifiers of the annotations. The options used with add-gff-info taxonomy were: -s 40 in order to use only BLAST results that have a bit score of at least 40, -l to use the last common ancestor algorithm, and -a 10 to use only the results that are within 10 bits from the maximum bit score for each annotation. For the CDS from the genomes, the resulting predicted taxonomic assignments were cross-checked against the species the genomes represented in order to provide validity for the *de novo* predictions for the CDS from the assembly.

### Gene family identification

A series of hierarchical clustering approaches were used to identify gene family clusters within and across species. Firstly, an all-against-all search of all the AA sequences was carried out using Diamond [40], where the maximum number of target sequences was set to 1,000,000 with a minimum bit score of 60 and all other options set to the default. Next, EGN was used to identify an initial round of gene similarity clusters using the Diamond all-against-all search results as input with the “gene network” option and the following settings: E-value threshold = 1e-05, hit identity threshold = 20%, identities must correspond at least to 20% of the smallest homolog, no best reciprocal condition, and no hit coverage condition enforced [41]. Secondly, all the AA sequences were used as input to the EGGNOG mapper (v1) [42] to generate (where possible) KEGG ortholog (KO) IDs for the sequences [43-46]. The KOs were then used to identify where different EGN gene clusters were recognised as having the same function, allowing them to be combined into higher-level functional clusters using igraph in R [47]. This was followed by further manual refinement of the functional groups in MS Excel using the information from the EGGNOG annotations of the AA sequences. Finally, the taxonomic assignment, functional cluster membership and expression (count) information across each of the samples for every predicted CDS was combined into an overall table for subsequent analyses.

### Network Analysis

To perform the co-expression network analysis at the family taxonomic level, the count data needed to be pre-processed. The taxonomic families with less than 10 genes expressed were removed from the dataset and the sum of the expression for all genes in the remaining taxonomic families was calculated. Next, to account for differences in sequencing depths in each sample, the summed count data were scaled by dividing each value by the sum of the total counts from the sample it belonged and then multiplying them by the median of all sample sums. Then the scaled data were further filtered using variable filtering based on Inter-quartile range (IQR; [48, 49]), where taxonomic families with expression lower than the 1st quartile (25th percentile) were removed. The final stage of data pre-processing was data normalisation, which was completed using regularised log transformation [50].

Co-occurrence networks were subsequently constructed based on the results of correlation analysis. The correlation analysis was conducted with the Spearman’s rho rank correlation and the results were filtered based on both correlation coefficients and their corrected p-values. Only the interactions with a correlation coefficient larger than 0.7 and (Benjamini-Hochberg) adjusted p-values less than 0.1 were used for the network analysis. The constructed network was then explored and visualised using the open-source software Gephi [51]. Modularity metrics were calculated in Gephi to detect the clusters in the constructed network.

Putative keystone taxa identification was performed within each cluster of the network [52]. We hypothesised that keystone taxa would have a large impact on the community network, and any absences of them should lead to major disruption to the network [53]. To describe the importance of keystone taxa, we used a set of network level measures: transitivity, density, modularity, average path length and centralisation of eigenvector. Transitivity measures the probability of the adjacent vertices. The graph’s density calculates how many edges are compared to the maximum possible number of edges between vertices. Modularity is designed to measure the strength of division of a network into modules. The average path length is calculated as the shortest paths between all pairs of vertices. Eigenvector centralisations measures from the centrality scores according to the eigenvector centrality of vertices. In order to determine the keystone taxa, we calculated these measurements with the full network and then again after removing a node. This was calculated for every node in the network. Thus, the differences in the measurements between each removal network and the complete network could be used to reveal those taxa whose removal had the largest detrimental effect on the community network structure and, therefore, could be identified as putative keystone taxa. Based on the mechanism of these measures, if one node has a large impact on the cluster, the differential values of transitivity and density should be reduced while modularity, average path length and centralisation of eigenvector should be increased. For each measure we ranked each node based on the difference calculated for these values following the nodes removal and further produced an overall ranking for each node using the Borda count method (sum of ranks), which is widely used in the voting method for decision making of multiple ranks. The overall top ranked families from the Borda count were identified as putative keystone taxa. To fulfil this task, we used R package igraph [47], the scripts and data used to carry out the analyses described above are available on the Open Science framework at https://osf.io/rx9h2/.

### Temporal gene expression analysis

In order to carry out the statistical analysis of gene expression, the expression information was summarised by taxonomic family and functional cluster (generated as earlier described). This resulted in raw count data for each timepoint from each functional cluster expressed in every taxonomic family [54]. This data was used as input to the Bioconductor package DESEQ2 in R [50]. In this analysis, the design was “cows+time”. Initial testing of the effect of cows and time was carried out using the likelihood ratio test “LRT” in DESEQ2, where first the effect of “cows+time” was tested against a reduced model of “time”, testing for any interactions with the cows the samples were taken from. Secondly the model “cow+time” was tested against a reduced model of “cows” to test the effect of time. Only those genes identified to have a significant interaction with time from the LRT (and excluding the 2 genes that had an interaction with cows) were retained for subsequent pairwise differential expression analysis. All 10 possible pairwise comparisons between the 5 sampled timepoints (excluding timepoint zero) was carried out to identify the pattern of differential expression (DE) for those genes with a significant interaction with time using a Benjamini-Hochberg p-adjusted cut-off of 0.1 to identify significance.

Finally, “significance groups” across time (analogous to the output from a Tukey’s HSD test) were identified for each gene in each taxonomic family to allow identification of patterns of DE over time and to allow clustering of genes by their shared DE pattern. This was carried out in R, using igraph, to identify for each gene in each taxonomic family the maximal cliques in the network of timepoints that were not significantly different to each other. These maximal cliques were then converted into labels for identifying the significance groups in subsequent visualisations. The R script and data used to carry out the temporal gene expression as described above is available in the Open Science framework at https://osf.io/rx9h2/.

### Detailed annotation and temporal abundance of glycosyl hydrolase, peptidase and quorum sensing genes

The functional clustering described earlier grouped all glycosyl hydrolases (GH) and peptidase genes into single clusters containing all variants of these enzymes. In order to facilitate a further in-depth analysis of the expression of these important enzymes, more detailed functional analysis of these two enzyme groups was carried out. Separation of GH families was carried out in reference to the annotations available in the CAZy database (http://www.cazy.org/; [55]) using DBCan (version 2.0.0) [56], Separation of the peptidases families was carried out in reference to the MEROPS [57] database using Clustal Omega (version 1.2.4) [58] to identify groups of peptidase families in the database which were then assembled into profiles using HMMER (version 3.3) [59]. These profiles were then used to assign peptidases to families using HMMER. To facilitate the statistical analysis of this subset of the dataset, the sum of the expression of all genes assigned to each GH and peptidase family was calculated for each taxonomic family.

For the quorum sensing (QS) genes, the functional clustering identified a cluster of S-ribosylhomocysteine lyase (LuxS) genes, involved in Auto-inducer-2 (AI-2) based QS, from 5 different taxonomic families. However, no clusters of N-acyl homoserine lactone (AHL) were identified, so no analysis could be undertaken for this gene. Next, the expression of each of the GH, peptidase and QS families was normalised by calculating the Transcripts Per Million (TPM) for every gene in the entire transcriptome in MS Excel, and then extracting the calculated values for the members of the GH, peptidase and QS families. This commonly used normalisation method for RNAseq data utilises the knowledge of the length of each gene and the total sequencing effort and provides an estimate of the number of RNA molecules from this gene for every million RNA molecules in the sample. The GH and peptidase families were then separately analysed as follows. First, the TPM for all genes identified as belonging to the same enzyme family were summed for statistical analysis. For the GH and peptidase families any that fell outside the 95% most highly expressed across all timepoints for that type (i.e. GH or peptidase) were excluded. Then for each GH and peptidase family an analysis of variance (ANOVA) was performed on the summed TPM values in order to identify any significant interactions with time, including cow as a fixed effect in the design (with the ‘aov’ command in R). For the QS genes the TPM values from each taxonomic family were summed and an ANOVA was performed to identify any significant interactions of the expression of the QS genes in each taxonomic family with time. All the ANOVA p-values were then corrected for multiple testing using the Benjamini-Hochberg method with value of <0.1 used to determine enzyme families with a significant interaction with time. For each of these with a corrected p-value < 0.1, a Tukey’s HSD test was performed (using the ‘TukeyHSD’ command in R) to identify where the significant differences were occurring and to assign significance groupings across timepoints. The contribution of each taxonomic family to the expression of each GH and peptidase family at each timepoint was also determined. All data was visualised using ‘ggplot2’ in R. Detailed isoform expression analysis of GH families 3, 5, 9, 10 and 48 was carried out by extracting the AA sequences for each separately and aligning with default options in Muscle (v3.8.1551; [60]) followed by phylogenetic tree reconstruction using RAxM-ng (v1.1.0; [61]) using a JTT+G model. Finally, the expression profiles of each isoform at each timepoint was mapped to the resulting phylogenetic tree using iTOL [62]. The R scripts used to carry out this analysis is available in the Open Science framework at https://osf.io/rx9h2/.

## Results

### Overview of sequencing data

We generated prokaryotic metatranscriptome data for PRG attached prokaryotes incubated over time (1, 2, 4, 6, and 8h) within the rumen of three cannulated Holstein x Friesian cows (two replicates/animal/timepoint with replicates within animals being pooled pre-sequencing) and the PRG attached bacteria at 0h (three 0h pre-incubation PRG RNA extractions being pooled pre-sequencing). Therefore, a total of 16 samples (i.e. 3 cows x 5 timepoints + 0h sample) were sequenced to assess the gene expression of PRG attached prokaryotes over time. Information on the number of reads generated, pre and post filtering, alignment to other datasets and the number of genes expressed, for each sample can be found in Supplementary Table 1. Average read abundance/sample was 14,825,273. However, 0 h sample reads were low at 2,060,217 and did not align to any of the Hungate genomes due to the fact that bacteria colonising PRG pre-incubation were epiphytic in nature (Supplementary Excel 1). Taxonomic families associated with this timepoint further decreased post-rumen incubation, therefore, these were not included in the subsequent metatranscriptome analysis (Supplementary Excel 1).

### Network analysis

Family-based gene network correlations, showing both positive and negative gene correlations are shown in Fig.1. The layout of the network is ForceAtlas2 [63] where connections or edges are presented that have correlation coefficients (Spearmans’s rho) larger than 0.7 and adjusted p-values less than 0.1. The sizes of the nodes are proportional to the expression of genes from the corresponding family across all timepoints. The size of the nodes indicates that (in descending order) *Lachnospiraceae, Prevotellaceae, Ruminococcaceae, Selemonadaceae, Spirochaetaceae, Methanobacteriaceae, Fibrobacteraceae* and *Eubacteriacaeae* dominate, irrespective of incubation time. The family *Lachnospiraceae* was dominated by two genera (*Butyrivibrio* and *Pseudobutyrivibrio)*, while the families *Prevotellaceae, Ruminococcaceae, Selemonadaceae, Spirochaetaceae, Methanobacteriaceae, Fibrobacteraceae* and *Eubacteriacaeae* were each comprised of a single genus only, namely: *Prevotella, Ruminococcus, Selenomonas, Treponema, Methanobrevibacter, Fibrobacter*, and *Eubacterium* respectively (Supplementary Excel 1). In the interest of not repeating family and genera in every instance, we refer mainly to families in the results presented below. Further information for genera within these and other families can be found in Supplementary File 1.

**Figure 1.**
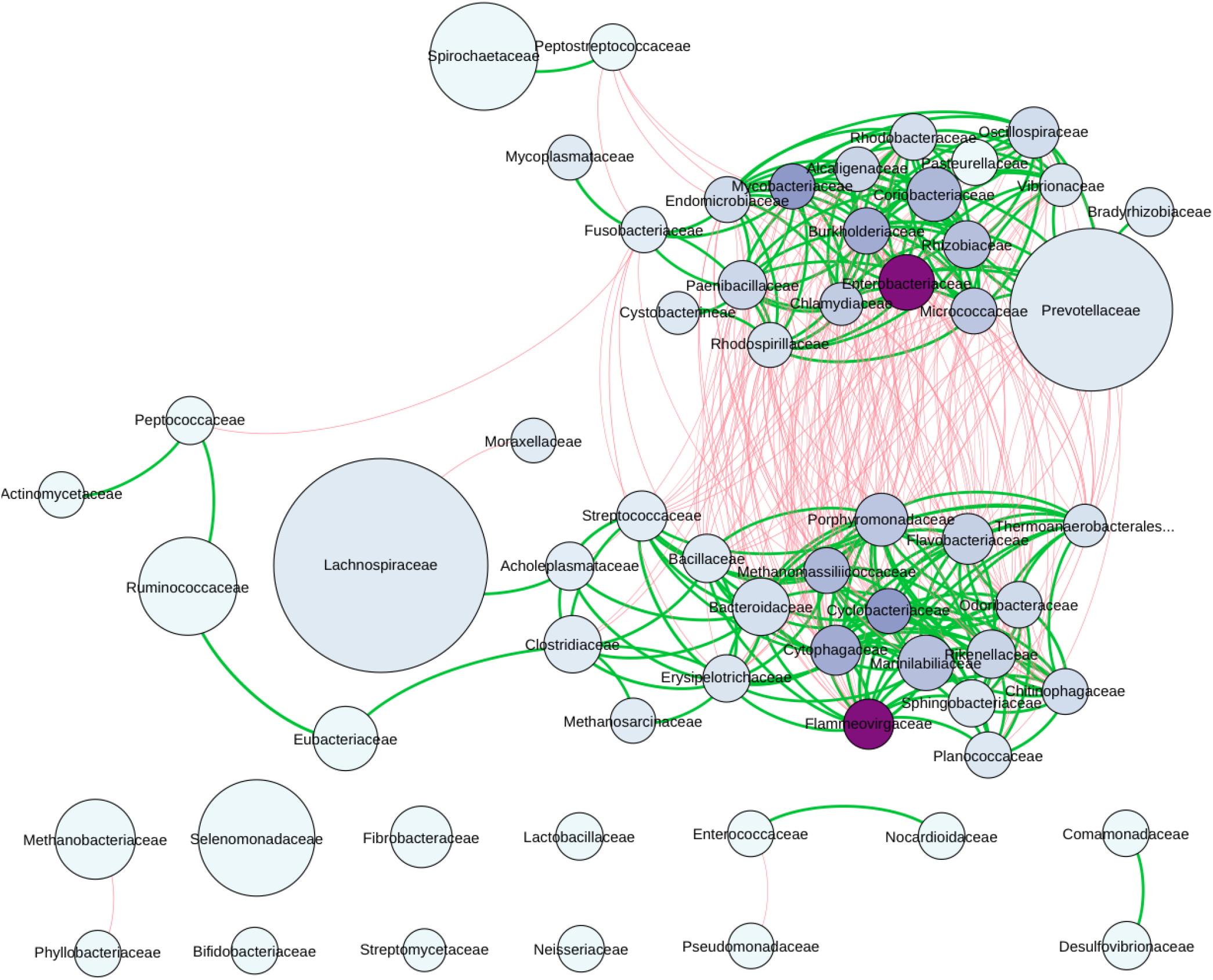
Co-occurrence gene network map showing positive (green lines) and negative gene correlations (red lines) for genes assigned to families. The size of the node denotes family relative abundance. Nodes in purple indicate putative keystone families.

The network analysis also shows that there are two large clusters in the network with strong positive correlations, and these clusters are separated by red lines which indicate negative gene expression-based correlations between the two large clusters (Fig. 1). These two clusters predominantly represent primary (≤4h post rumen incubation) and secondary (≥4h post rumen incubation) colonisation phases and can be considered as two sub-microbiomes (Fig. 1). The primary sub-microbiome is largely dominated by *Prevotellaceae*, whilst the secondary sub-microbiome is dominated by *Lachnospiraceae*, and to a lesser extent *Ruminococcaceae* and *Eubacteriaceae*. A further group of bacterial families are independent of the primary and secondary sub-microbiomes, namely families *Selemonadaceae, Spirochaetaceae, Methanobacteriaceae, Bifidobacteriaceae, Fibrobacteraceae, Lactobacilliaceae, Enterococcaceae, Pseudomonadaceae, Phyllobacteraceae, Streptomycaceae, Neisseriaceae, Nocardiodiaceae, Commomonadaceae, Desulfovibrionaceae* and *Pasturellaceae*, (in descending order of relative abundance). Also, of particular note, is the finding that the dominant families had few or even no gene correlation, suggesting selfish non-cooperative behaviour, although *Prevotellaceae* was an exception (Fig.1).

To identify keystone taxa, a set of node measurements were calculated. Among these measurements five centrality metrics (transitivity, density, modularity, average path length and centralisation of eigenvector) were also used to identify putative keystone taxa as previously done by others [53, 64-66]. These measurements included five centrality metrics: transitivity, density, modularity, average path length and centralisation of eigenvector. For the primary sub-microbiome the highest-ranking putative keystone families were *Burkholderiaceae* and *Enterobacteriaceae*, and for secondary sub-microbiome *Cyclobacteriaeceae* and *Flammeovirigaceae* (Fig. 1; Supplementary Tables 2 & 3). This suggests these families have an important role in the cohesion of these submicrobiomes although they have a much lower activity than the dominant families and are within the lower 10% of those reported in Supplementary Fig. 2A & B, and hence are not directly shown within the graph.

### Temporal niche specialisation

A total of 1,513 genes were differentially expressed (DE) by the prokaryotes colonising PRG in the rumen over time (these values exclude genes lower than 10% of total expression levels) (Fig. 2A). These genes were representative of all general functions, including cellular processing, information storage and processing, metabolism, and others which were poorly characterised or unknown (Fig. 2B). Twenty-eight distinct DE patterns across all five timepoints were observed in these genes, with as few as three genes and up to as many as 170 genes showing any particular DE pattern (Fig. 2). The DE pattern with the highest total expression across all timepoints consisted of 69 genes, and was expressed in higher abundance during secondary colonisation (≥ 4 h rumen incubation) compared to primary colonisation (≤4 h rumen incubation) (Fig. 2C and D). These 69 genes were mainly expressed by *Lachnospiraceae*, with lower expression also evident within the *Prevotellaceae* (Fig. 2E). Interestingly, the DE pattern with the second-highest expression (consisting of 96 genes) showed the opposite pattern (i.e. higher abundance during primary colonisation) and was dominated by *Prevotellaceae*, consistent with the network analysis. The remaining DE patterns identify further timepoint and taxa-specific expression profiles worthy of further in-depth analysis. Summarising these DE patterns by function identified many bacterial families with time-point specific functional roles (Figs 3-8; Supplementary Figs 3-32). As there are too many temporal functional changes evident within this complex microbiome to outline all in detail here, we have provided the raw data and graphical representation of the taxonomy by temporal expression patterns for all Enzyme Commission (EC) categories in supplementary material (Supplementary Figs 3-32) for those with an interest in specific microbial functions. In this study we concentrated on genes involved in methane metabolism, protein, carbohydrate breakdown and quorum sensing.

**Figure 2.**
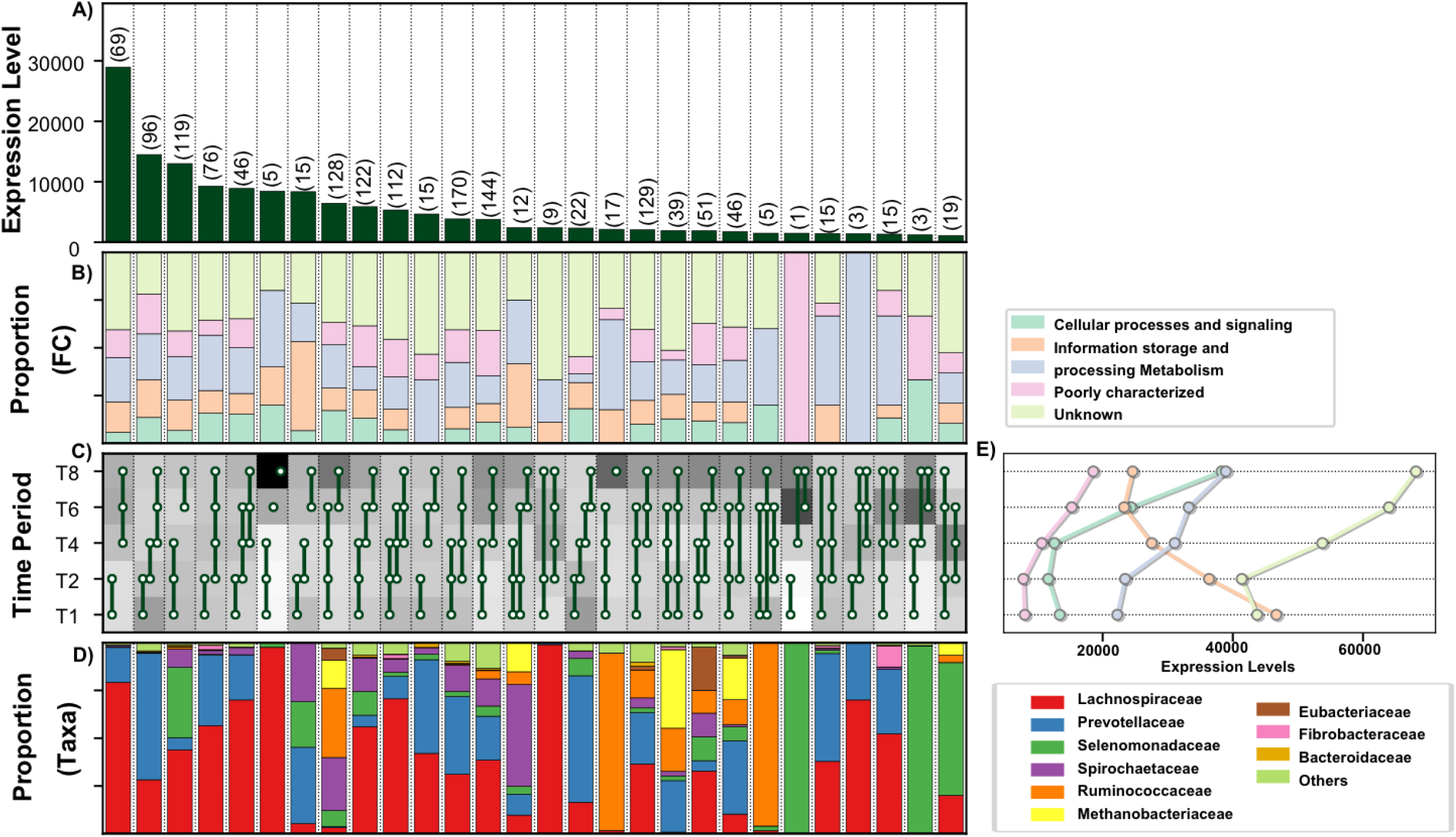
Temporal functional and taxonomic overview of the top 90% most highly expressed genes with a significant interaction with time expressed by prokaryotes attached to fresh perennial ryegrass incubated *in situ* within the rumen. Each column represents a set of genes that showed the same differential expression (DE) pattern (denoted as expression pattern on the x axis). A) Summed expression level of all the genes with the same DE pattern, and in brackets is the corresponding number of genes within the same DE pattern. B) Proportion of each major functional category (FC) represented in the set of genes with the same DE pattern. C) Visual representation of the DE patterns for each set of genes across the timepoints sampled (i.e. T1 is the 1h timepoint) where: (i) the background heatmap represents the level of expression for each timepoint (Low = White, High = Black) and (ii) the lines and dots represent the specific DE pattern shared by all genes in this set where the timepoint dots connected by a line and do not significantly different from each other. D) The proportion of the taxonomic families contributing to the expression level for each DE pattern. E) The level of expression of the major functional categories across each timepoint.

**Figure 3.**
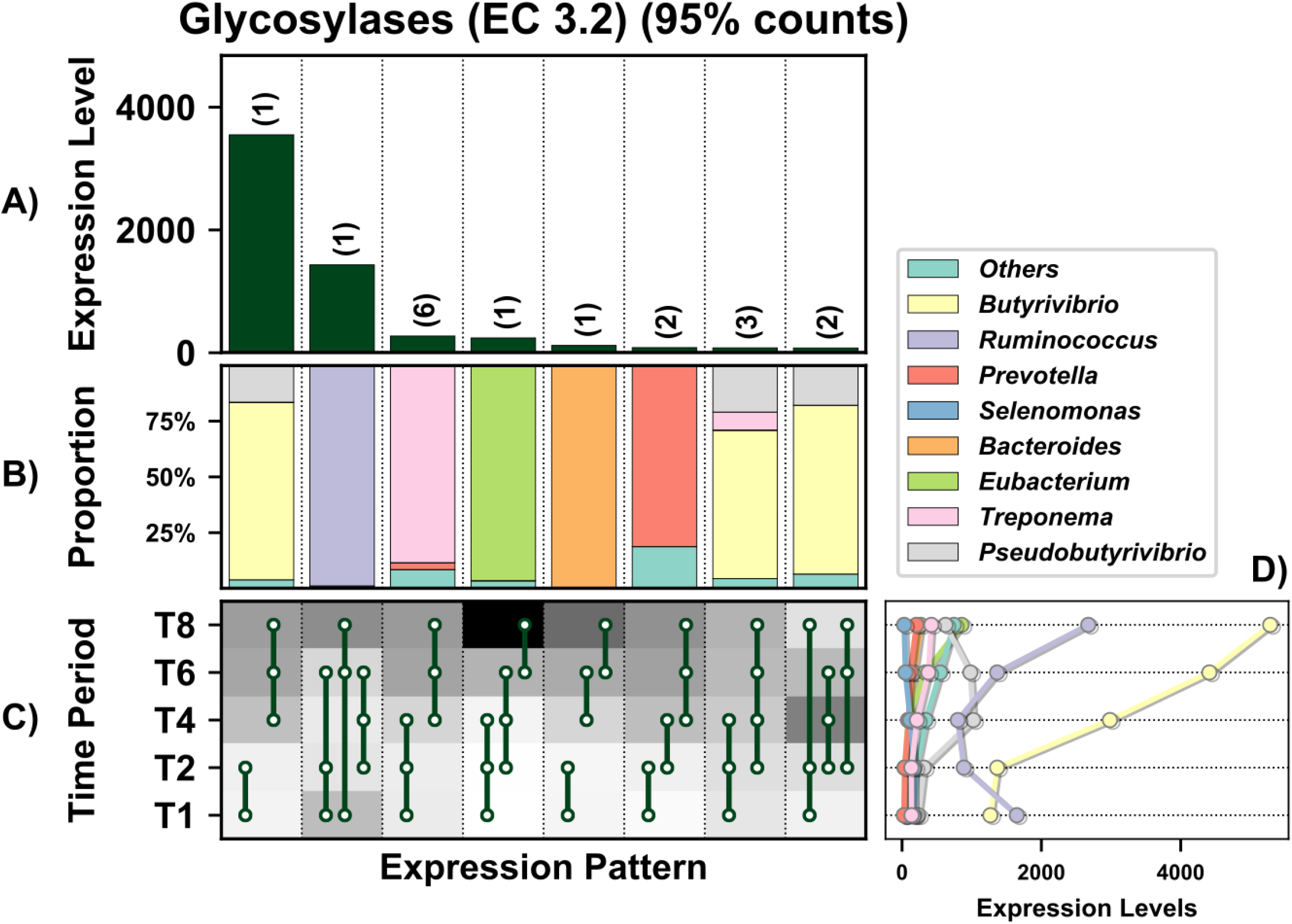
Temporal expression of the top 95% most highly expressed genes with a significant interaction with time involved in the KEGG methane metabolism pathway within the top 95% of expressed by prokaryotes attached to fresh perennial ryegrass incubated *in situ* within the rumen. Each column represents a set of genes that showed the same differential expression (DE) pattern (denoted as expression pattern on the x axis). A) Summed expression of all methane metabolism genes with the same DE pattern, in brackets the number of genes with the same DE pattern. B) The proportion of taxonomic genera contributing to the expression level for each DE pattern. C) Visual representation of the DE patterns for each set of genes across the timepoints sampled; The heatmap represents the level of expression for each timepoint (Low = White, High = Black); The lines and dots represent the specific DE pattern shared by all genes in this set where the timepoints connected by line and dots were not significantly different from each other. D) The level of expression of the genes from each taxonomic family across each timepoint.

**Figure 4.**
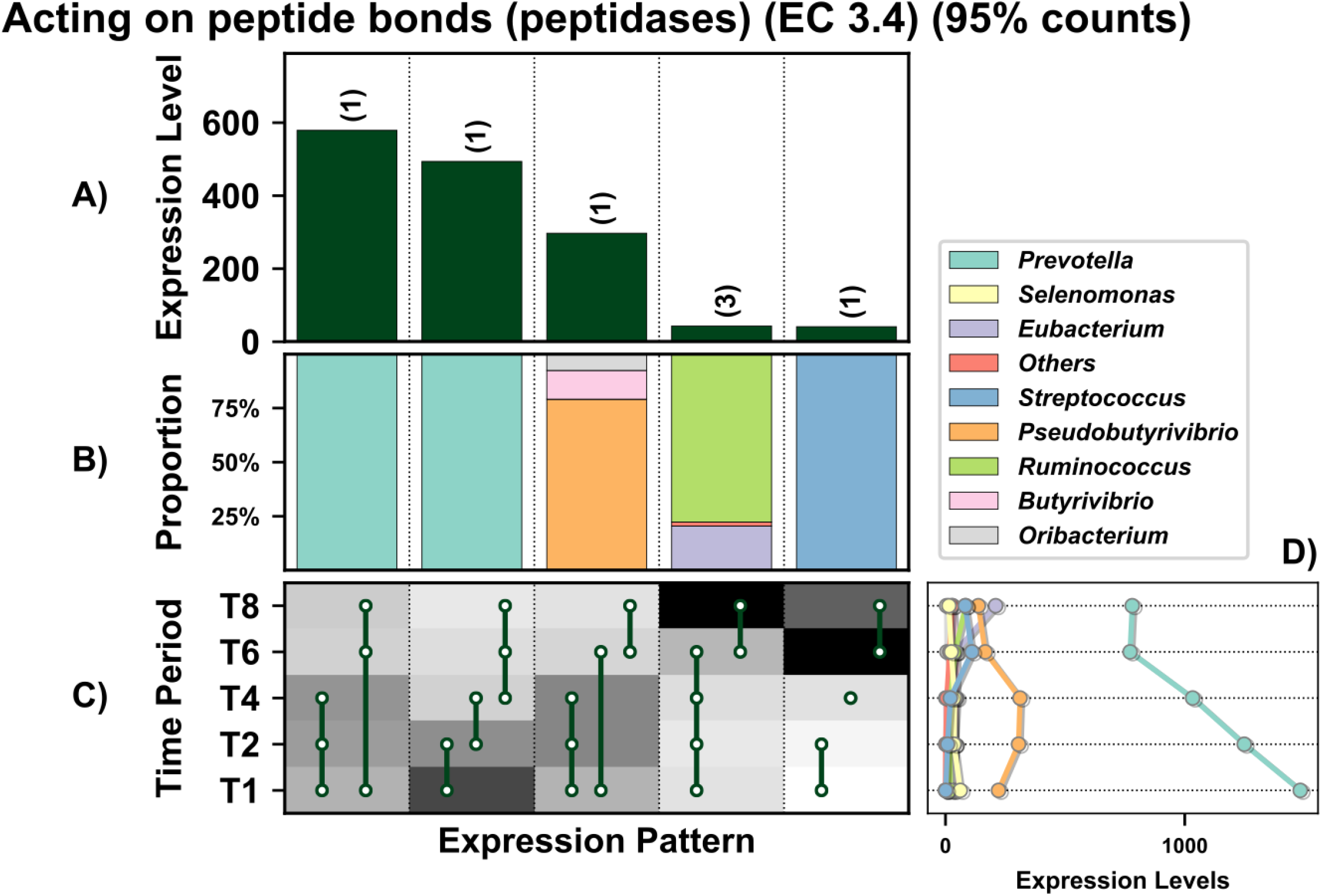
Overview of the temporal expression of the top 95% most highly expressed peptidase genes (EC 3.4) with a significant interaction with time expressed by prokaryotes attached to fresh perennial ryegrass incubated *in situ* within the rumen. Each column represents a set of genes that showed the same differential expression (DE) pattern (denoted as expression pattern on the x axis). A) Summed expression of all genes with the same DE pattern, in brackets the number of genes with the same DE pattern. B) The proportion of taxonomic genera contributing to the expression level for each DE pattern. C) Visual representation of the DE patterns for each set of genes across the timepoints sampled; The heatmap represents the level of expression for each timepoint (Low = White, High = Black); The lines and dots represent the specific DE pattern shared by all genes in this set where the timepoints connected by line and dots were not significantly different from each other. D) The level of expression of genes across each timepoint.

**Figure 5.**
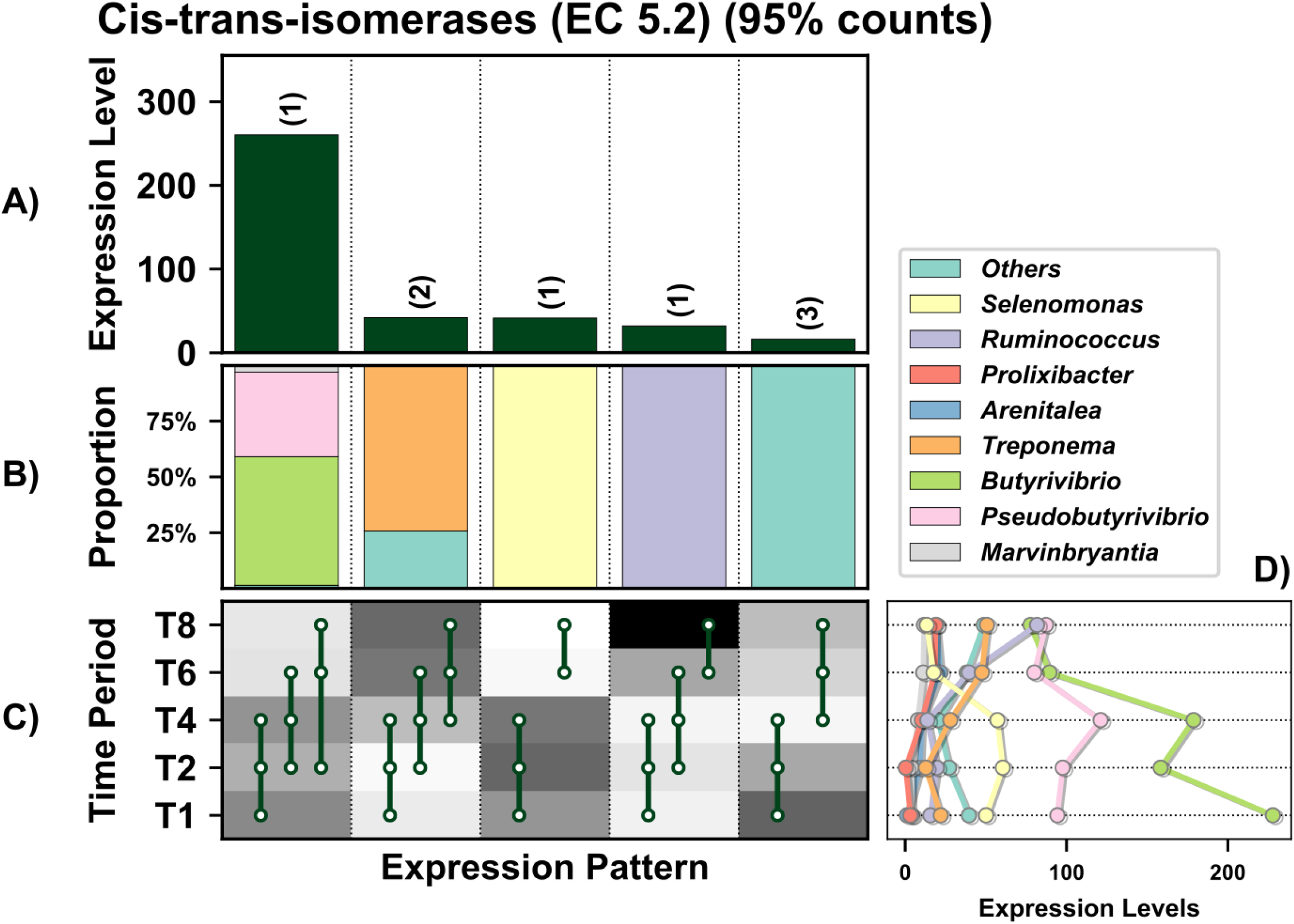
Overview of the temporal expression of the top 95% most highly expressed glycosyl hydrolase genes (EC 3.2) with a significant interaction with time expressed by prokaryotes attached to fresh perennial ryegrass incubated *in situ* within the rumen. Each column represents a set of genes that showed the same differential expression (DE) pattern (denoted as expression pattern on the x axis). A) Summed expression of all genes with the same DE pattern, in brackets the number of genes with the same DE pattern. B) The proportion of taxonomic genera contributing to the expression level for each DE pattern. C) Visual representation of the DE patterns for each set of genes across the timepoints sampled; The heatmap represents the level of expression for each timepoint (Low = White, High = Black); The lines and dots represent the specific DE pattern shared by all genes in this set where the timepoints connected by line and dots were not significantly different from each other. D) The level of expression of genes across each timepoint.

**Figure 6.**
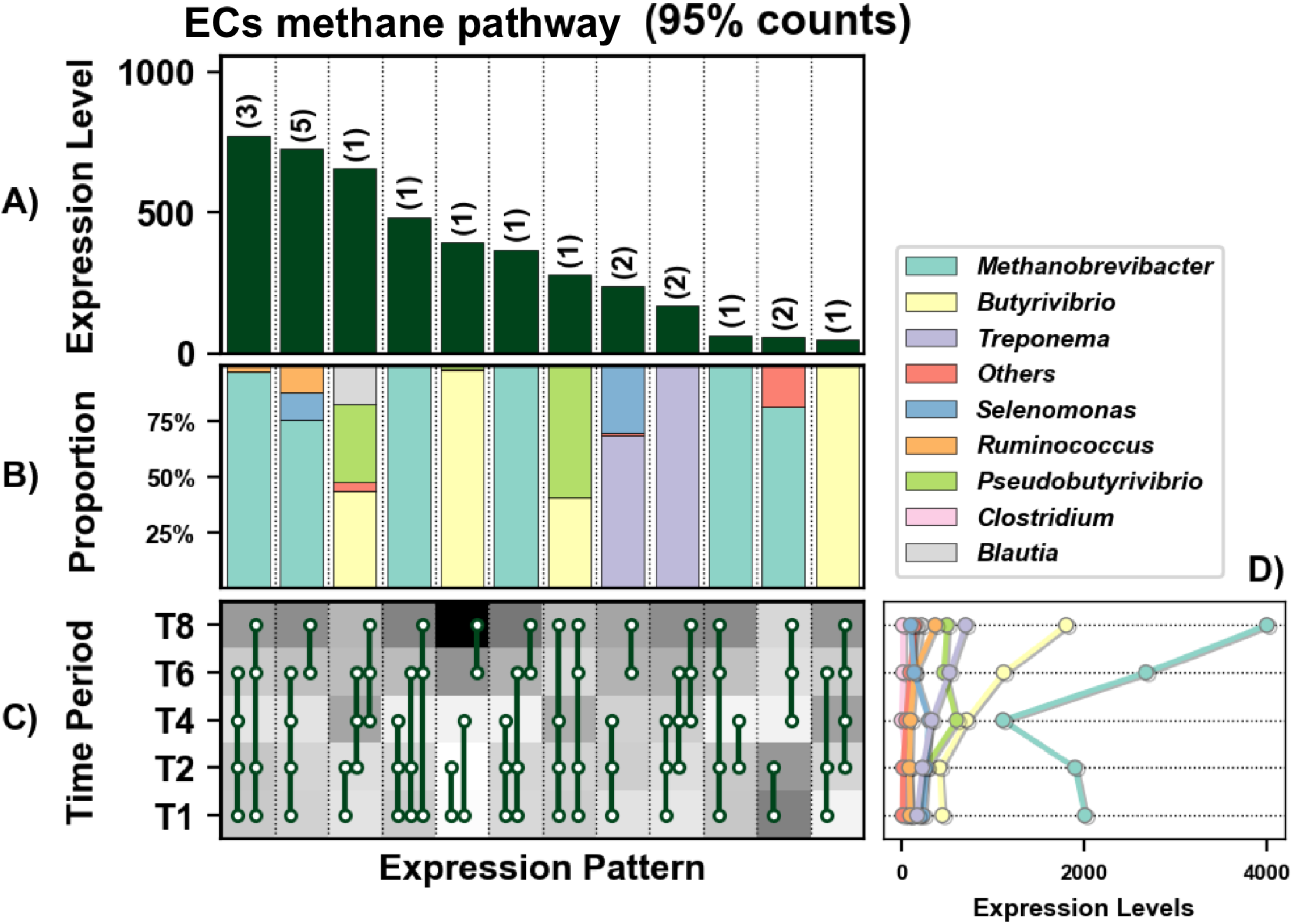
In depth analysis of the temporal expression of differentially expressed carbohydrate active enzyme (CAZymes, also known as glycosyl hydrolases (GH)) expressed genes by prokaryotes attached to fresh perennial ryegrass incubated within the rumen that differed significantly in their expression profile over rumen incubation time (line plots) and their respective taxonomic origins (bar chart below the corresponding line plot). Incubation time is indicated on the axis of the plots, i.e. T1 indicates an incubation time of 1 hour. Brown bars: family Eubacteriaceae (genus *Eubacterium*); Pink bars: family Fibrobacteriaceae (genus *Fibrobacter*); Red bars: family Lachnospiraceae (genera *Butyrivibrio* and *Pseudobutyrivibro*); Blue bars: family Prevotellaceae (genus *Prevotella*); Orange bars: Ruminococcaceae (genus *Ruminococcus*); Purple bars: Spirochaetaceae (genus *Treponema*). The significance of rumen incubation time on gene expression is indicated on each plot, with timepoint that significantly differ denoted by a different letter in the line plot

**Figure 7.**
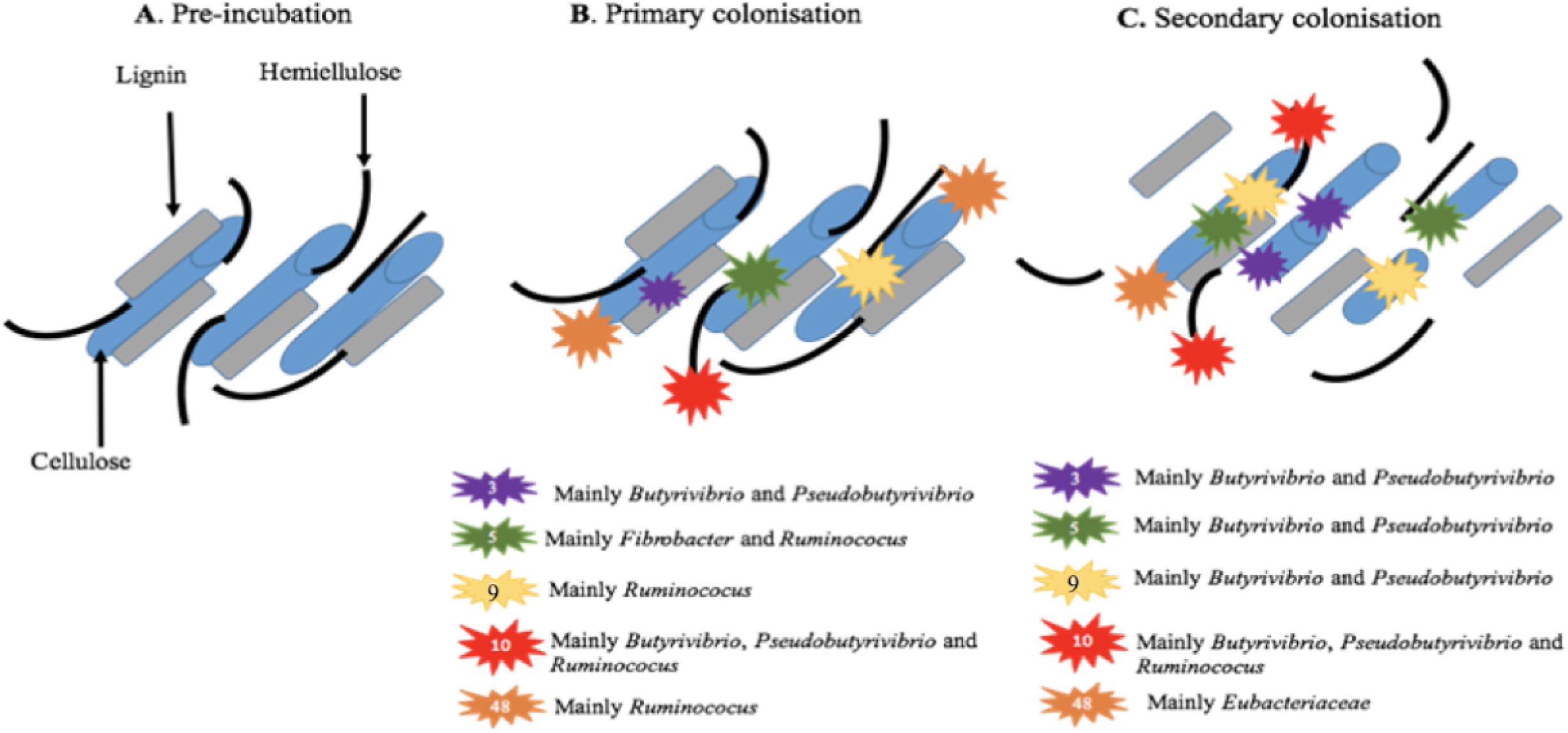
Diagrammatic representation of fresh perennial ryegrass carbohydrate breakdown over incubation time within the rumen. Diagram illustrates only the dominant carbohydrate active enzymes (CAZymes, also known as glycosyl hydrolases (GH)).GH family numbers are shown by the numbers in the symbol key. Taxonomic origins of the expressed CAZymes are also shown in the key next to the corresponding symbol. Number of GH symbols between primary (<4h) and secondary (>4h) colonisation sub-microbiomes are representative of whether expression has increased or remained constant.

**Figure 8.**
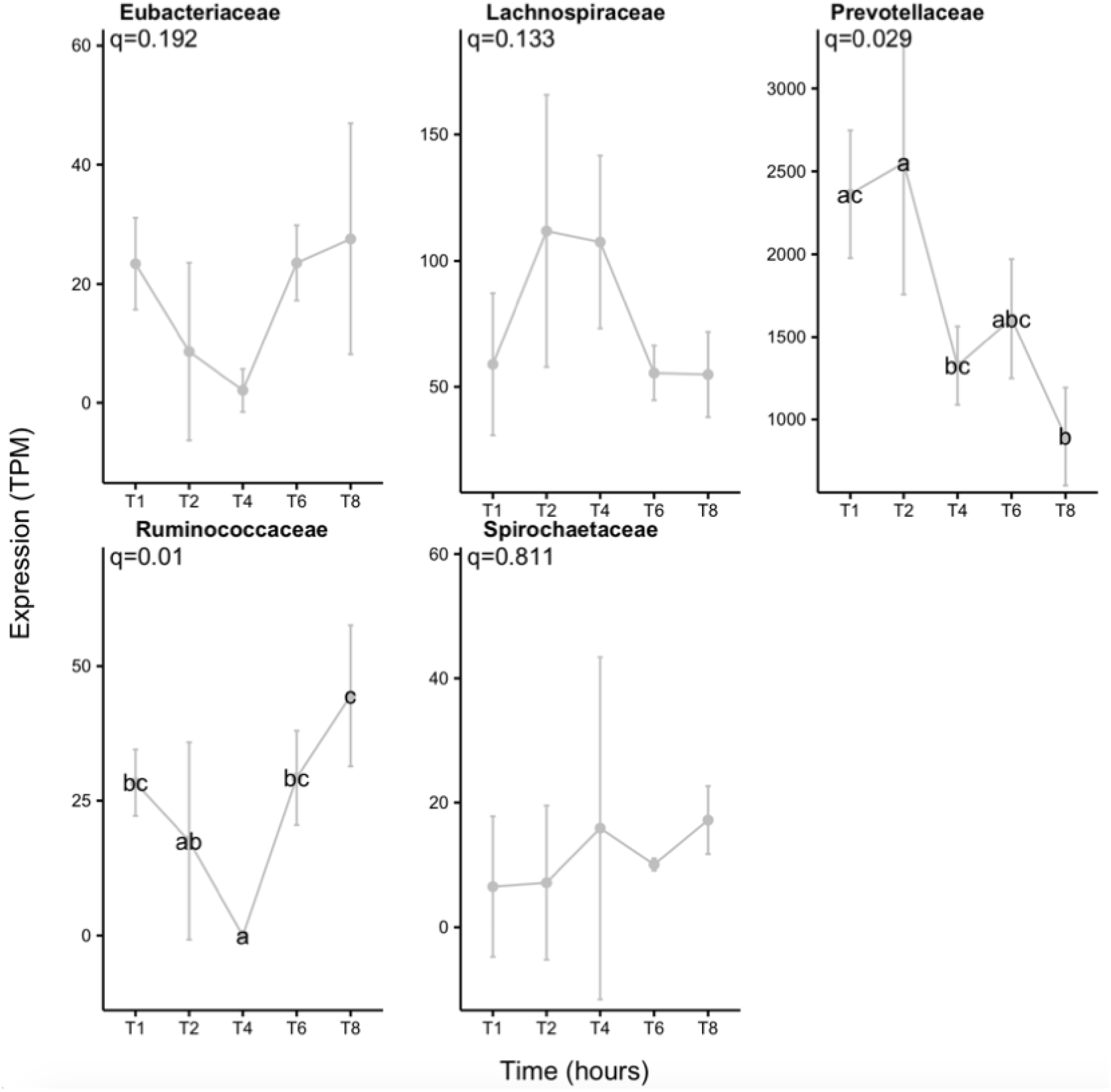
Number of expressed LuxS genes (Transcripts per million,TPM) for each prokaryotic taxonomic family colonising fresh perennial ryegrass incubated in the rumen over time. Incubation time is indicated on the axis, i.e. T1 indicates an incubation time of 1 hour. The significance of incubation time is indicated on each plot, and where significance occurs then differences between timepoints are denoted by a different letter.

Genes involved in methane metabolism were mainly expressed by *Lachnospiraceae* (interestingly mainly by genus *Butyrivibrio* and not *Pseudobutyrivibrio*) and *Methanobacteriaceae*, with expression increasing during secondary colonisation (≥4 h rumen incubation; Fig. 3; Supplementary Excel 1). Given that *Butyrivibrio* dominated the expression profile (Supplementary Excel 1) and they produce butyrate and hydrogen from fermentation of complex and simple carbohydrates, it is likely that the *Methanobrevibacter*, which utilise hydrogen to produce methane, increase in response to the higher hydrogen levels allowing them to increase methane formation and release as supported by KEGG information (Supplementary Figure 33).

Peptidase genes are important to the host in terms of breaking down protein. In total 7 genes encoding peptidases (EC 3.4) were detected and these were generally more highly expressed during primary colonisation compared with secondary colonisation, in particular within the family *Prevotellaceae*, which specialise in this activity (Figure 4; Supplementary Excel 1). When MEROPS was used for a more detailed analysis of peptidase family expression (Supplementary Excel 1; Figure 4) this indicated that 38 peptidase families were within the top 95% of all expression of this type of gene (Supplementary excel 1; Supplementary Figure 34). These genes families were most predominantly expressed by: M50, *Ruminococcaceae*; M24b, *Lachnospiraceae*; s01c, *Prevotellaceae* and *Ruminococaceae*; M16b, *Prevotellaceae* (Supplementary Excel 1; Supplementary Figure 34). In terms of temporal changes in peptidase expression, significant changes over time were noted only for M24b and M15a (expressed mainly by the bacterial families *Selemonadaceae* and *Ruminococcaceae*), with expression increasing during secondary colonisation (Supplementary Excel 1; Supplementary Figure 34). This is the converse to what was seen with overall peptidase (EC 3.4) expression data and a consequence of obtaining more granular data using MEROPS. Further elaboration of their precise role was not possible as the exact peptidase function for most of the peptides could not be identified.

Genes encoding glycosyl hydrolases (often also called carbohydrate active enzymes; CAZymes) (EC 3.2; 17 in total) were mainly up-regulated during secondary colonisation (≥ 4h rumen incubation), especially within the families *Lachnospiraceae* and *Ruminococcaceae*, which specialise in carbohydrate breakdown (Fig. 6; Supplementary Excel 1). When DBcan was then used to analyse the CAZymes, further resolution was obtained (Supplementary Excel 1; Figure 35). DBcan showed that a total of 18 CAZymes represented 95% of all expression of this type of gene in the PRG-colonising prokaryotes (Supplementary Figure 35) with enzyme families GH3, 5, 10, 9 and 48 dominating the expression profiles, in descending order (Supplementary Excel 1; Figure 35). Significant changes over time were noted for GH1 (β-glucosidases and β-galactosidases; mainly expressed by *Selemonadaceae*), 3 (endo-β-1,4-xylanases; mainly expressed by Lachnospiraceae, 5 (endo-β-1,4-xylanases; mainly expressed by *Fibrobacteraceae* and *Ruminococcaceae*), 9 (endo-β-1,4-xylanases; mainly expressed by *Ruminococcaceae*), 10 (endo-endo-beta-1,4-xylanases; mainly expressed by *Ruminococcaceae*), 11 (endo-β-1,4-xylanases; *Ruminococcaceae*), 13 (act on substrates containing α-glucoside linkages; *Spirochaetacea*e and *Selemonadaceae*), 26 (endo-β-1,4-mannanases; *Eubacteriaceae, Fibrobacteraceae, Ruminococcaceae* and *Selemonadaceae*) and 57 (includes α-amylases, α-galactosidases, amylopullulanases and 4-α-glucanotransferases; *Fibrobacteraceae* and Prevotellaceae). All of these apart from GH1 increased during secondary colonisation compared to primary colonisation (Fig 6; Supplementary Excel 1 and Figure 35). In order to contextualise this temporal fresh PRG carbohydrate degradation on a genus basis, a schematic figure was constructed which illustrates the changes in niche specialisation by the rumen bacteria, e.g. GH5 is mainly expressed by *Fibrobacter* and *Ruminococcus* during primary colonisation and by *Butyrivibrio* and *Pseudobutyrivibrio* during secondary colonisation (Fig. 7). Intriguingly, when these data are examined at an even more detailed level, it is apparent that different isoforms of each of the dominant GH families (GH3, 5, 9, 10 and 48) have time-point specific activity (Supplementary Figures 36-40).

To further understand potential drivers of switches from primary to secondary colonisation events, the temporal expression of cell signalling molecules was also investigated. AI-2-based LuxS quorum sensing temporal expression data showed that these genes were mainly expressed by *Prevotellacae*. It was also evident that *Prevotellaceae* increased expression of LuxS genes during primary colonisation and thereafter declined, which may partially drive the transition to a secondary sub-microbiome (Fig. 8).

## Discussion

Microbial interactions are complex and often explained as being either positive, neutral or negative [21, 22], with pairs of effects often found, such as in mutualism and competition. Combinations of positive, neutral and negative outcomes can also be found, for example under amensalism and commensalism [21]. This study investigated the breadth of temporal ecological interactions and niche-specialisation of fresh PRG-attached rumen prokaryotes. Using metatranscriptomics and network analysis we show, on a gene expression network basis, that temporal colonisation of fresh PRG involves an array of ecological interactions, with cooperation, mutualism as well as competitive behaviours evident. Primary (≤4h rumen incubation) and secondary (≥4h rumen incubation) PRG-attached sub-microbiomes were evident over time, as evidenced by antagonistic negative interactions between the two large clusters. Network analysis also enabled definition of interactions within these sub-microbiomes, including identification of putative keystone families. In terms of key functional processes for the host, while protein breakdown is a continual process with limited differences seen over time and between sub-microbiomes, carbohydrate degradation is higher for many CAZymes during secondary colonisation which are mainly related to breakdown of complex carbohydrates. Dominant bacteria also commonly change their function over incubation time, presumably because of the need to be ecologically plastic in a changing environment.

In terms of prokaryotic expression, the top 5 families were *Prevotellaceae, Lachnospiraceae, Selemondaceae, Ruminococcaeae* and *Fibrobacteraceae*. The family *Prevotellaceae* showed most transcriptional activity during primary colonisation, whilst the families *Lachnospiraceae* and *Ruminococcaceae* tended to be more transcriptionally active during secondary colonisation. The families *Fibrobacteriaceae* and *Ruminococcaceae* showed reasonably equal activity across primary and secondary colonisation events. These temporal shifts of families are in line with the 16S rRNA (based on cDNA) data already published for the same samples showing that RNA-based metataxonomy and metatranscriptome data findings were comparable [10]. Gene network analysis also showed that these dominant bacterial families were largely non-cooperative, as on a gene expression basis they did not interact substantially with the two sub-microbiomes, with the exception being *Prevotellacea*e. This is perhaps a consequence of the fact that their adaptation to this environment reduces the need to behave cooperatively and, therefore, could be characterised as ‘selfish’ [9]. Conversely, the less abundant bacterial families *Burkholderiaceae, Enterobacteriaceae, Cyclobacteriaeceae* and *Flammeovirigaceae* displayed highly cooperative behaviour in either the primary or secondary temporally distinct sub-microbiomes. This suggests that they may have an important role as keystone families for effective attachment and degradation of PRG.

Keystone bacteria are normally defined as species that if removed would have a negative effect on the structure and function of the community [53]. These four families are not high in abundance in the rumen [67] and were low in activity in this study. However, it seems they may play an integral role in their respective sub-microbiomes and possibly aid connectivity within the biofilms. Keystone taxa are often described as being cooperative and mutualistic, but in the context of plant colonisation in the rumen, the low levels of activity of these organisms suggest that they may be displaying commensalism by ‘cheating’ [68], perhaps capitalising on the activity of others in the sub-microbiomes. This strategy would ensure the community stays together, rewarding synergistic behaviours in the sub-microbiome, allowing harvesting of energy from the activities of other groups with relatively little input. These results also raise questions about the minimum rumen prokaryotic diversity required for effective energy harvesting, the importance of minor keystone species to this activity, and whether the presence of the dominant genera alone would be more energetically efficient. Recent data showed that ruminants with less diverse rumen microbiomes, based on bacteria only, are more feed efficient [69], suggesting that degradation of fresh PRG may be equally as efficient in the presence of the dominant prokaryotic families *Lachnospiraceae* and *Prevotellaceae* alone and comparative to the situation when they are among a complex microbiome.

The underlying mechanisms driving competitive antagonism between primary and secondary associated sub-microbiomes likely include nutrient availability and production of chemicals by the microbes themselves [70]. During early rumen incubation (<6h), fresh PRG is known to undergo self-mediated proteolysis and lipolysis, which will alter the form of the nutrients available to the rumen microbiome [71-73]. In parallel, it is known that PRG will be changing in terms of chemical content due to the rumen microbial breakdown of complex carbohydrates, lipids and proteins over time, thus providing a continually evolving niche for the microbes to inhabit [6, 10, 74]. In terms of plant nutrient degradation, we found that peptidase expression is high amongst the attached rumen prokaryotes, with limited DE seen between primary and secondary colonisation events. Most of the peptidase expression was by *Prevotella, w*ith species of this genera well known for their dominant proteolytic activity within the rumen microbiome [75, 76]. Carbohydrate breakdown was also a dominant activity of the fresh PRG attached bacteria, as denoted by the high expression of CAZymes/GHs. The preponderance of CAZyme activity within the rumen bacteria is well known and underpins their key function of energy harvesting by breaking down and fermenting of complex carbohydrates to volatile fatty acids [77, 78]. Nonetheless, the temporal expression and contribution of each CAZyme family to fresh PRG has, to our knowledge, not been demonstrated before. We show that GH3, 5, 9, 10 and 48 dominate in terms of expression by the PRG-attached rumen bacteria, all of which are involved in the degradation of recalcitrant plant cell wall cellulose and hemicellulose, with all (all apart from GH48, which remains constant) slightly increasing over incubation time. Our data also highlight the niche specialisation and plasticity amongst the attached rumen bacteria as GH5 and 9 are expressed predominantly by *Fibrobacter* and/or *Ruminococcus* during primary colonisation but during secondary colonisation GH5 and 9 are predominantly expressed by *Butyrivibrio* and *Pseudobutyrivibrio*. This clearly demonstrates the redundancy, resilience and niche plasticity that occurs within the PRG-attached rumen microbiome. We also demonstrate that GH family gene isoforms exist as has been shown previously [9] and that these within family isoforms are differentially expressed over attachment time, further illustrating the redundancy that underpins the resilience of the rumen microbiome. Increases in methane metabolism was also apparent during secondary colonisation and this was linked to *Butyrivibrio, Pseudobutyrivibrio* and *Methanobrevibacter* activity. This is proposed to be due to increased hydrogen release from carbohydrate breakdown by the two bacteria genera being utilised by *Methanobrevibacter* to produce methane.

We also postulated that alongside known chemical changes in the fresh PRG [20], bacterial cell signalling and chemical warfare may also drive temporal niche specialisation. For example, in this study, we found that family *Prevotellaceae* and genus *Prevotella*, in particular, expressed AI-2 LuxS increasingly during primary colonisation and thereafter their expression during secondary colonisation declines. These data supports the previous report that *Prevotella* were the most active in terms of expressing AI-2 LuxS genes in the rumen [17]. We also hypothesized that bacterial cell signalling may influence colonisation events and niche specialisation as it has previously been shown that furanosyl borate diester molecules encoded by the LuxS gene cause biofilm dispersal in many pure culture models [79]. Interestingly, no AHL genes were detected, which is consistent with other data, which suggests that AHL based quorum sensing is low in the rumen [17]. Genes encoding the antimicrobial peptide (AMP) Lynronne 1 (likely produced by *Prevotella ruminocola*) were also up-regulated during the primary colonisation phase of PRG and decreases in expression during the secondary phase [80]. In contrast, we showed that expression of non-ribosomally synthesised peptides and polyketide increased during secondary colonisation within fresh PRG attached bacteria [81]. The DE of these ribosomally and non-ribosomally synthesised AMPs and polyketides during the transition from primary to secondary colonisation may well contribute towards the antagonism between the bacteria characteristic of the two phases of PRG colonisation. This suggests that alongside mutualism, competition due to chemical warfare may be rife in the rumen and contribute to differences in niche occupancy over time. Interestingly, within these studies, expression of LuxS and ribosomally and non-ribosomally synthesised AMPs and polyketides was most pronounced within the genus *Prevotella*. Network analysis also showed that family *Prevotellaceae*, of which we only identified the genus *Prevotella*, had more negative than positive interactions. However, it is likely that plant chemical changes, both self-induced and microbially-mediated, alongside cell signalling and chemical warfare all collectively contribute towards the ecological plasticity seen within biofilms occurring at the plant interface.

In summary, our findings substantially expands our ecological understanding of microbial energy harvesting and niche development within the rumen. Furthermore, this study provides a major step change in our understanding of the microbial diversity and function required within a utopian microbiome for optimal host phenotype, whilst providing insight into the desired plant characteristics needed for maximum energy harvesting.

## Data accessibility

Reads of the sequence data have been deposited in the NCBI Short Read Archive under bioproject accession no. PRJNA419191. All scripts and intermediate data are deposited on the Open Science framework at https://osf.io/rx9h2/.

## Competing interests

The authors declare no competing interests.

## Author contributions

SAH, JEE, JAP and AKS conceived the project. PS and SAH completed the laboratory work. MA, DS and SC completed the library preparation and sequencing and CC, FR, WL completed downstream computational analysis with contribution from MW and LBO. SAH, CC, JEE and FR interpreted the results with contribution from MW and LBO. All authors contributed to the preparation of the manuscript and approved its final version.

## Funding

This work was supported by the Biotechnology and Biological Sciences Research Council Institute Strategic Programme Grant, Rumen Systems Biology (grant number BBS/E/W/10964) and The Genome Analysis Centre (now the Earlham Institute) Capacity and Capability Challenge Programme.

## Notes

### Competing Interest Statement

The authors have declared no competing interest.

