## Supplementary material for "Microbiomes attached to fresh perennial ryegrass- are temporally resilient and adapt to changing ecological niches"

**Supporting information**

**Supplementary Excel 1.** Taxonomy & annotations, peptidase and glycosyl hydrolase data.

**Supplementary Table 1.** Sequencing data and alignment information.

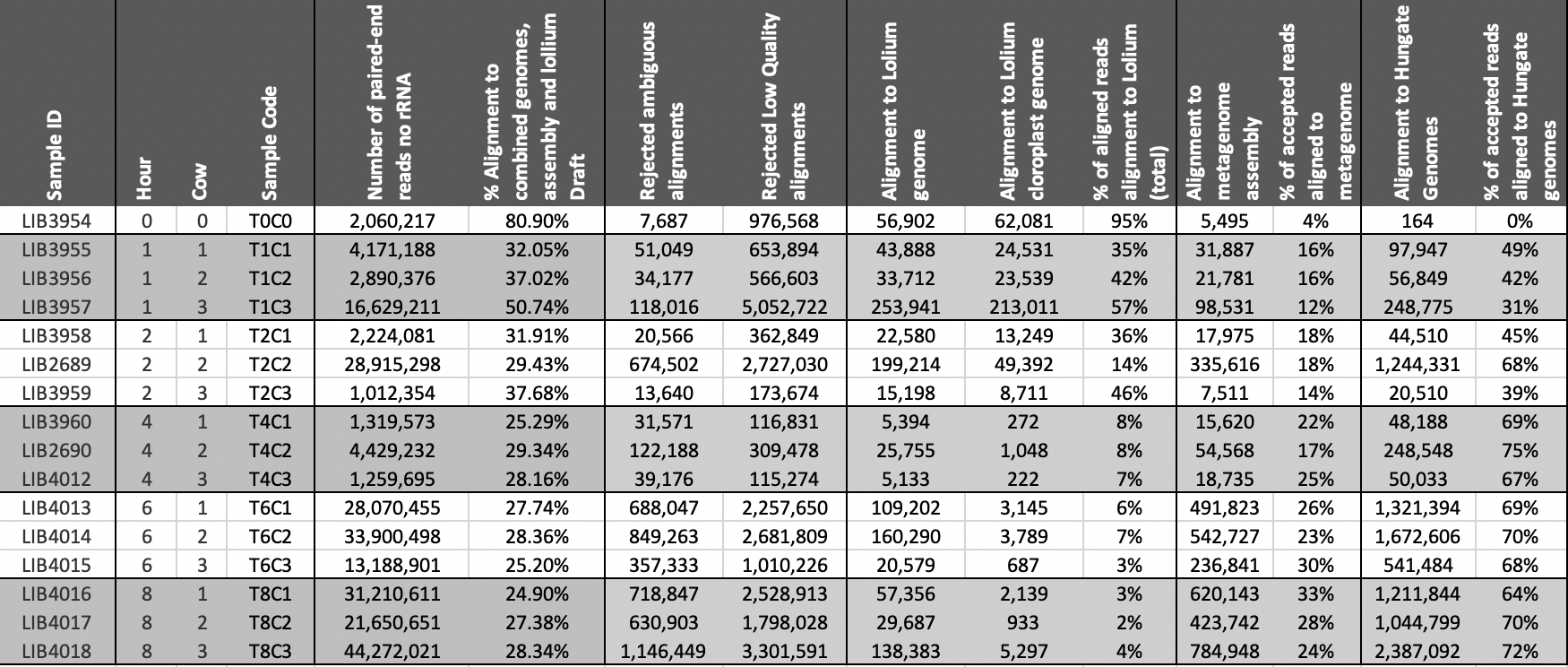

**Supplementary Table 1.** Parameters used for identification of putative keystone taxonomic families within the primary colonising sub-microbiome

| **Taxonomic Family** | **Transitivity diff** | **Density**  **Diff** | **Modularity**  **diff** | **AvePathLen**  **diff** | **CentraEigen**  **diff** | **Transitivity rank** | **Density rank** | **Modularity**  **rank** | **AvePathLen**  **Rank** | **CentraEige**  **rank** | **Sum of Ranks** | **Overall**  **rank** |
| --- | --- | --- | --- | --- | --- | --- | --- | --- | --- | --- | --- | --- |
| **Burkholderiaceae** | **-0.024494** | **-0.020256** | **0.015984** | **0.032821** | **0.021189** | **1** | **4** | **2** | **5** | **3** | **15** | **1** |
| **Enterobacteriaceae** | **-0.024494** | **-0.020256** | **0.015984** | **0.032821** | **0.021189** | **1** | **4** | **2** | **5** | **3** | **15** | **1** |
| **Rhizobiaceae** | **-0.024379** | **-0.02359** | **0.01706** | **0.036154** | **0.009229** | **3** | **1** | **1** | **3** | **10** | **18** | **3** |
| **Coriobacteriaceae** | **-0.015592** | **-0.02359** | **0.006752** | **0.059487** | **0.022978** | **7** | **1** | **8** | **1** | **2** | **19** | **4** |
| **Mycobacteriaceae** | **-0.014053** | **-0.02359** | **0.007012** | **0.052821** | **0.02674** | **8** | **1** | **7** | **2** | **1** | **19** | **4** |
| **Micrococcaceae** | **-0.016009** | **-0.016923** | **0.007491** | **0.029487** | **0.013453** | **6** | **6** | **6** | **7** | **6** | **31** | **6** |
| **Chlamydiaceae** | **-0.019243** | **-0.01359** | **0.012857** | **0.026154** | **0.013337** | **4** | **8** | **4** | **9** | **7** | **32** | **7** |
| **Tolypothrichaceae** | **-0.018521** | **-0.016923** | **0.00409** | **0.029487** | **0.018468** | **5** | **6** | **9** | **7** | **5** | **32** | **7** |
| **Alcaligenaceae** | **-0.010044** | **-0.006923** | **0.009796** | **0.016154** | **0.005689** | **9** | **12** | **5** | **13** | **13** | **52** | **9** |
| **Paenibacillaceae** | **0.001343** | **-0.010256** | **-0.004116** | **0.036154** | **0.008043** | **12** | **11** | **15** | **3** | **12** | **53** | **10** |
| **Oscillospiraceae** | **-0.003048** | **-0.01359** | **-0.005123** | **0.026154** | **0.009432** | **11** | **8** | **19** | **9** | **9** | **56** | **11** |
| **Endomicrobiaceae** | **0.002282** | **-0.01359** | **-0.004751** | **0.026154** | **0.011805** | **14** | **8** | **18** | **9** | **8** | **57** | **12** |
| **Rhodobacteraceae** | **-0.005755** | **-0.006923** | **0.003426** | **0.016154** | **0.003935** | **10** | **12** | **11** | **13** | **14** | **60** | **13** |
| **Acidaminococcaceae** | **0.002327** | **-0.000256** | **0.000301** | **0.009487** | **-0.003262** | **15** | **15** | **12** | **16** | **16** | **74** | **14** |
| **Rhodospirillaceae** | **0.012461** | **-0.00359** | **-0.008242** | **0.009487** | **0.009013** | **19** | **14** | **20** | **16** | **11** | **80** | **15** |
| **Eggerthellaceae** | **0.025436** | **0.00641** | **0.003864** | **0.019487** | **-0.010651** | **23** | **18** | **10** | **12** | **19** | **82** | **16** |
| **Vibrionaceae** | **0.013247** | **-0.000256** | **-0.010974** | **0.012821** | **-0.007602** | **20** | **15** | **21** | **15** | **18** | **89** | **17** |
| **Cystobacterineae** | **0.008595** | **0.029744** | **-0.002998** | **-0.053846** | **-0.020321** | **18** | **20** | **14** | **18** | **20** | **90** | **18** |
| **Mycoplasmataceae** | **0.002432** | **0.033077** | **-0.001472** | **-0.090513** | **-0.024176** | **16** | **21** | **13** | **20** | **22** | **92** | **19** |
| **Bradyrhizobiaceae** | **0.004063** | **0.033077** | **-0.004188** | **-0.080513** | **-0.023399** | **17** | **21** | **16** | **19** | **21** | **94** | **20** |
| **Moraxellaceae** | **0.00162** | **0.033077** | **-0.004188** | **-0.097179** | **-0.024603** | **13** | **21** | **16** | **22** | **23** | **95** | **21** |
| **Prevotellaceae** | **0.013657** | **-0.000256** | **-0.019919** | **-0.090948** | **-0.000608** | **21** | **15** | **23** | **21** | **15** | **95** | **21** |
| **Fusobacteriaceae** | **0.019971** | **0.013077** | **-0.0158** | **-0.112687** | **-0.004507** | **22** | **19** | **22** | **23** | **17** | **103** | **23** |

**Supplementary Table 2.** Parameters used for identification of putative keystone taxonomic families within the secondary colonising sub-microbiome

| **Taxonomic Family** | **Transitivity**  **diff** | **Density_diff** | **Modularity diff** | **AvePathLen diff** | **CentraEigen diff** | **Transitivity**  **rank** | **Density**  **rank** | **Modularity rank** | **AvePathLe**  **rank** | **CentraEigen**  **rank** | **Sum of Ranks** | **Overall**  **rank** |
| --- | --- | --- | --- | --- | --- | --- | --- | --- | --- | --- | --- | --- |
| **Flammeovirgaceae** | **-0.014005** | **-0.018696** | **0.014439** | **0.028116** | **0.01926** | **3** | **2** | **4** | **4** | **1** | **14** | **1** |
| **Cyclobacteriaceae** | **-0.014005** | **-0.018696** | **0.014439** | **0.028116** | **0.01926** | **3** | **2** | **4** | **4** | **1** | **14** | **1** |
| **Cytophagaceae** | **-0.011396** | **-0.022319** | **0.017428** | **0.035362** | **0.009304** | **6** | **1** | **1** | **3** | **10** | **21** | **3** |
| **Methanomassiliicoccaceae** | **-0.009152** | **-0.018696** | **0.017285** | **0.028116** | **0.018052** | **11** | **2** | **2** | **4** | **3** | **22** | **4** |
| **Marinilabiliaceae** | **-0.014153** | **-0.015072** | **0.011473** | **0.024493** | **0.016052** | **1** | **6** | **7** | **8** | **5** | **27** | **6** |
| **Porphyromonadaceae** | **-0.009152** | **-0.018696** | **0.017285** | **0.028116** | **0.018052** | **11** | **2** | **2** | **4** | **3** | **22** | **4** |
| **Prolixibacteraceae** | **-0.014153** | **-0.015072** | **0.011473** | **0.024493** | **0.016052** | **1** | **6** | **7** | **8** | **5** | **27** | **6** |
| **Flavobacteriaceae** | **-0.011175** | **-0.011449** | **0.011056** | **0.02087** | **0.012697** | **7** | **8** | **9** | **10** | **7** | **41** | **8** |
| **Rikenellaceae** | **-0.013054** | **-0.011449** | **0.008529** | **0.017246** | **0.011517** | **5** | **8** | **10** | **12** | **8** | **43** | **9** |
| **Leptospiraceae** | **-0.00186** | **-0.011449** | **0.013789** | **0.02087** | **0.009772** | **13** | **8** | **6** | **10** | **9** | **46** | **10** |
| **Odoribacteraceae** | **-0.010633** | **-0.007826** | **0.005608** | **0.002754** | **0.0069** | **8** | **11** | **12** | **14** | **11** | **56** | **11** |
| **Chitinophagaceae** | **-0.010633** | **-0.007826** | **0.005608** | **0.002754** | **0.0069** | **8** | **11** | **12** | **14** | **11** | **56** | **11** |
| **Hymenobacteraceae** | **-0.010633** | **-0.007826** | **0.005608** | **0.002754** | **0.0069** | **8** | **11** | **12** | **14** | **11** | **56** | **11** |
| **Bacteroidaceae** | **0.019921** | **-0.007826** | **-0.011817** | **0.042609** | **0.001475** | **23** | **11** | **20** | **2** | **14** | **70** | **14** |
| **Thermoanaerobacterales_Family_III._Incertae_Sedis** | **0.002956** | **-0.00058** | **0.007369** | **0.006377** | **-0.000265** | **17** | **16** | **11** | **13** | **16** | **73** | **15** |
| **Crocinitomicaceae** | **-0.001744** | **-0.00058** | **0.004897** | **-0.004493** | **-0.000448** | **14** | **16** | **15** | **17** | **17** | **79** | **17** |
| **Erysipelotrichaceae** | **0.026539** | **-0.004203** | **-0.024449** | **0.071594** | **-0.004232** | **25** | **15** | **22** | **1** | **18** | **81** | **18** |
| **Sphingobacteriaceae** | **-0.000644** | **-0.00058** | **0.004897** | **-0.004493** | **0.000134** | **15** | **16** | **15** | **17** | **15** | **78** | **16** |
| **Planococcaceae** | **0.017221** | **0.013913** | **0.003455** | **-0.018986** | **-0.015404** | **21** | **19** | **17** | **21** | **19** | **97** | **19** |
| **Bacillaceae** | **0.022093** | **0.013913** | **-0.016117** | **-0.004493** | **-0.018293** | **24** | **19** | **21** | **17** | **20** | **101** | **20** |
| **Lachnospiraceae** | **0.00129** | **0.046522** | **-0.005391** | **-0.124058** | **-0.030446** | **16** | **25** | **18** | **24** | **25** | **108** | **21** |
| **Methanosarcinaceae** | **0.005323** | **0.042899** | **-0.010627** | **-0.062464** | **-0.028312** | **18** | **24** | **19** | **23** | **24** | **108** | **21** |
| **Clostridiaceae** | **0.009588** | **0.028406** | **-0.03207** | **-0.037101** | **-0.022385** | **19** | **22** | **25** | **22** | **21** | **109** | **23** |
| **Streptococcaceae** | **0.018462** | **0.021159** | **-0.030901** | **-0.008116** | **-0.024021** | **22** | **21** | **24** | **20** | **22** | **109** | **23** |
| **Acholeplasmataceae** | **0.009662** | **0.032029** | **-0.027285** | **-0.182029** | **-0.024282** | **20** | **23** | **23** | **25** | **23** | **114** | **25** |

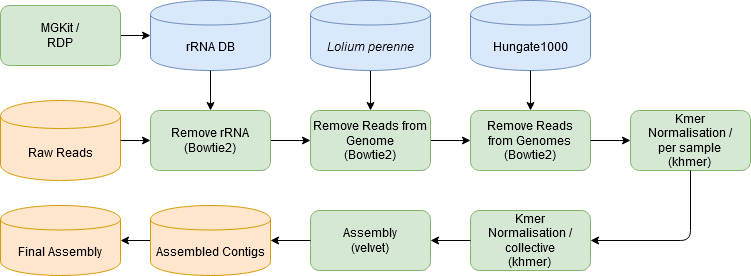

**Supplementary Figure 1.** Flow chart detailing the methodology used to assemble the raw reads into contigs for further analysis. In green the software used, in blue the public data used and in orange the study data.

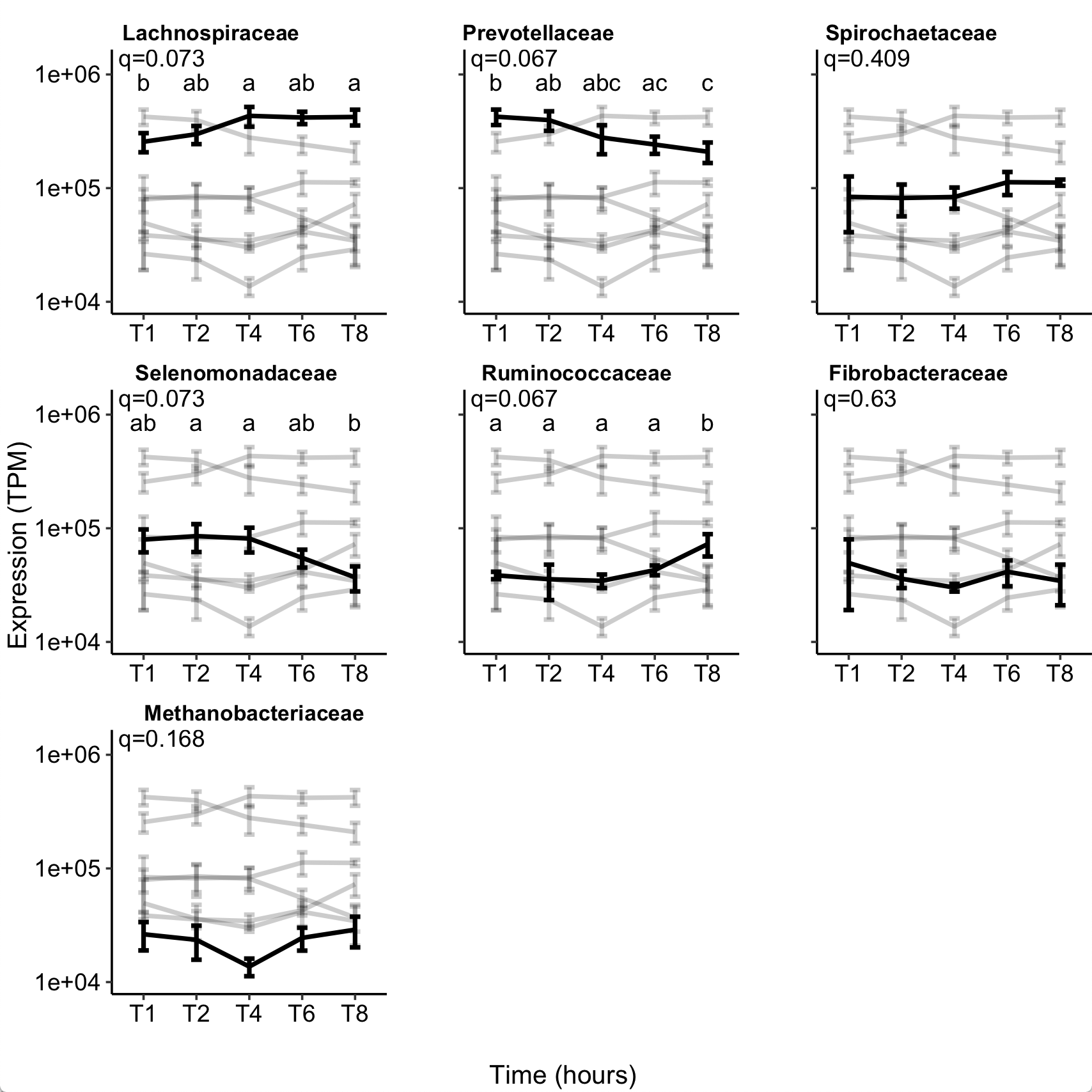

**Supplementary Figure 2.** Number of expressed sequencing reads (Transcripts per million, TPM), shown using a log scale, for each prokaryotic taxonomic family colonising perennial ryegrass incubated in the rumen over time. The seven taxonomic families representing 95% of the gene expression data from all timepoints are shown (i.e. T1 represents 1 hour of rumen incubation). The black lines shows the data for the representative family named above the plot, and fainter greyscale lines represent the data for the other families in order to show overall proportion of the highlighted family compared with all other dominant families.

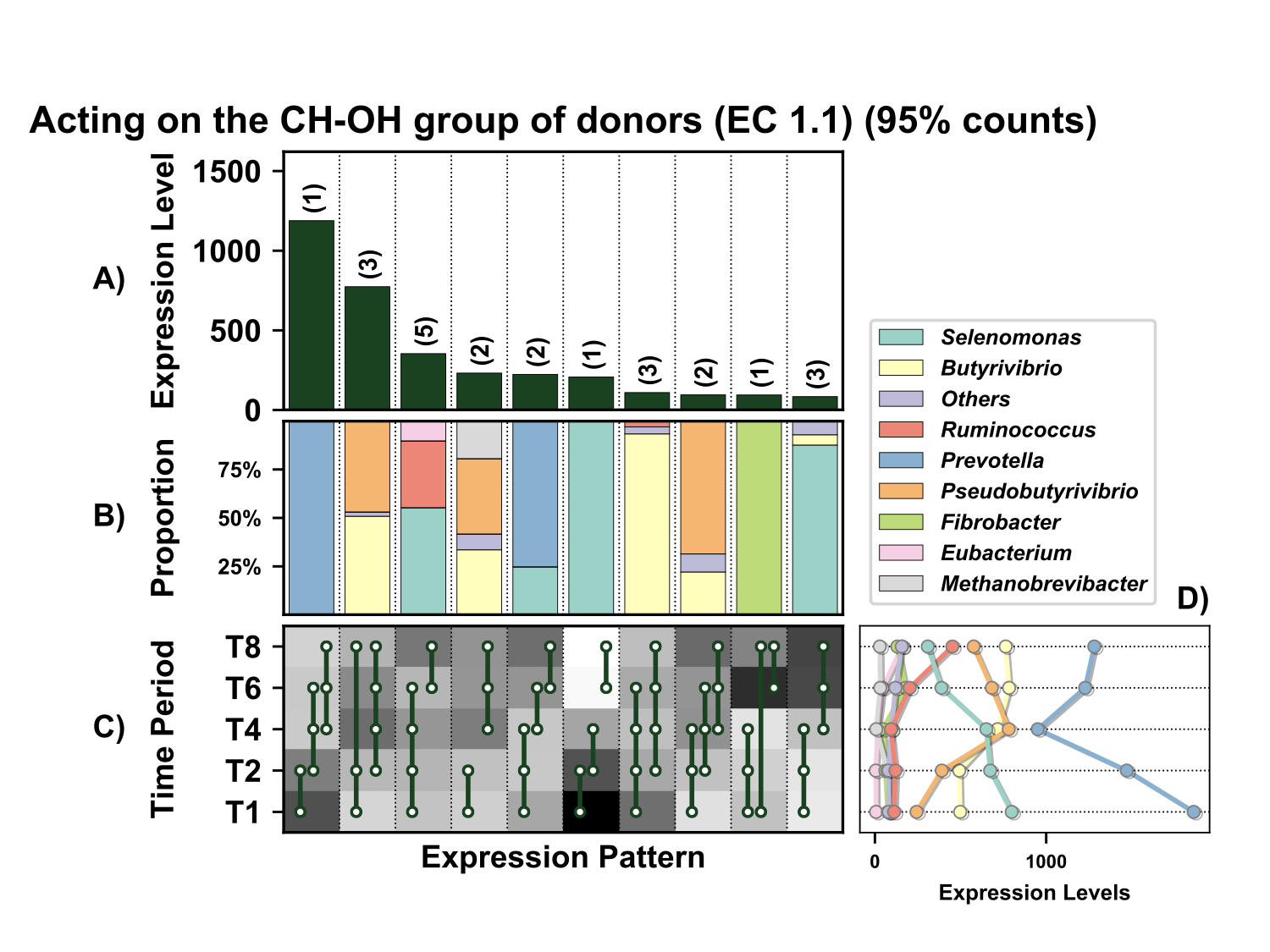

**Supplementary Figure 3.** Temporal expression of the top 95% most highly expressed genes acting on the CH-OH group of donors (EC 1.1) with a significant interaction with time expressed by prokaryotes attached to fresh perennial ryegrass incubated *in situ* within the rumen. Each column represents a set of genes that showed the same differential expression (DE) pattern (denoted as expression pattern on x axis). A) Summed expression of all genes with the same DE pattern, in brackets the number of genes with the same DE pattern. B) The proportion of taxonomic genera contributing to the expression level for each DE pattern. C) Visual representation of the DE patterns for each set of genes across the timepoints sampled; The heatmap represents the level of expression for each timepoint (Low = White, High = Black); The lines and dots represent the specific DE pattern shared by all genes in this set where the timepoints connected by line and dots were not significantly different from each other. D) The level of expression of genes across each timepoint.

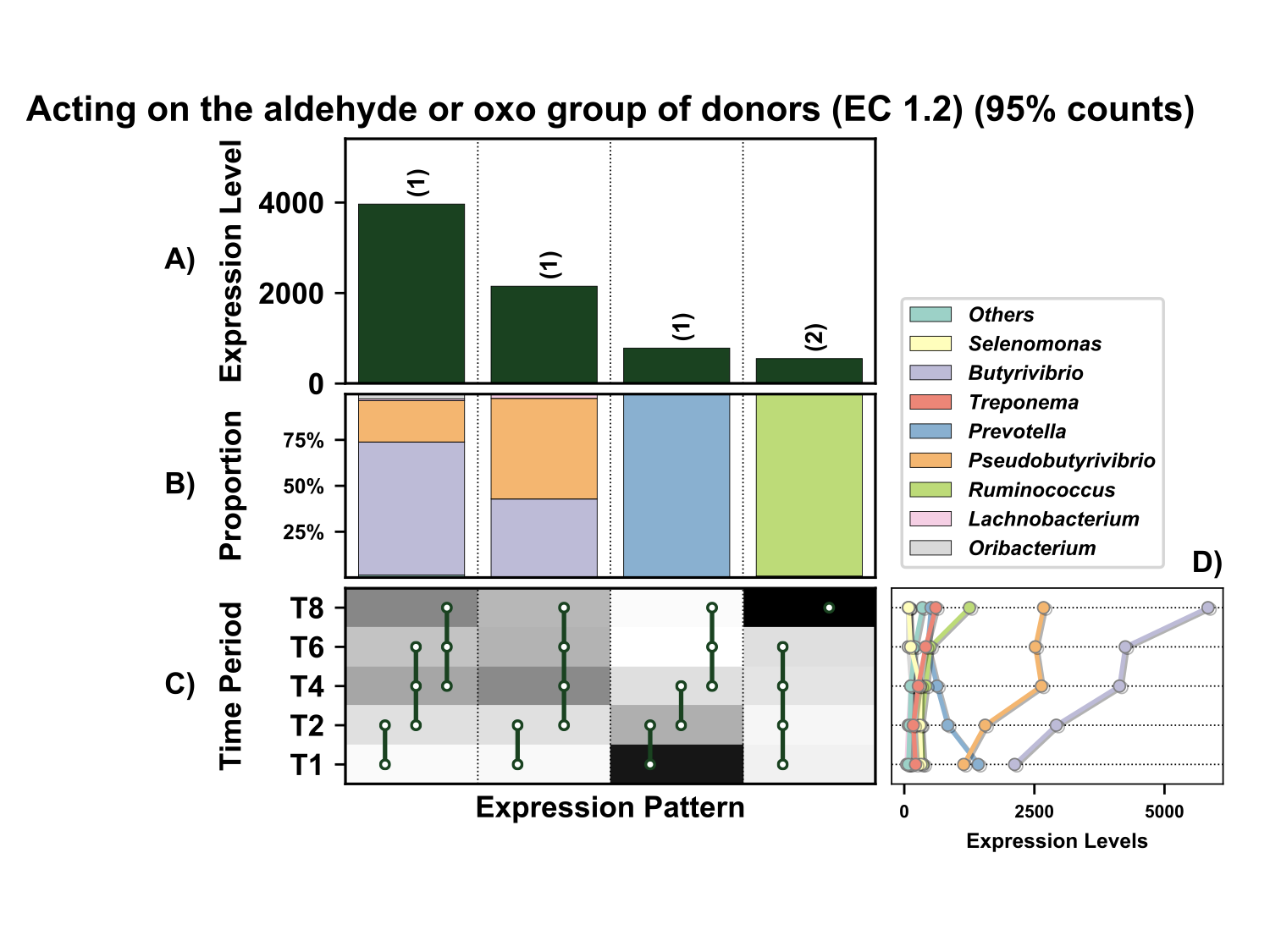

**Supplementary Figure 4.** Temporal expression of the top 95% most highly expressed genes acting on the aldehyde or oxo group of donors (EC 1.2) with a significant interaction with time expressed by prokaryotes attached to fresh perennial ryegrass incubated *in situ* within the rumen. Each column represents a set of genes that showed the same differential expression (DE) pattern (denoted as expression pattern on x axis). A) Summed expression of all genes with the same DE pattern, in brackets the number of genes with the same DE pattern. B) The proportion of taxonomic genera contributing to the expression level for each DE pattern. C) Visual representation of the DE patterns for each set of genes across the timepoints sampled; The heatmap represents the level of expression for each timepoint (Low = White, High = Black); The lines and dots represent the specific DE pattern shared by all genes in this set where the timepoints connected by line and dots were not significantly different from each other. D) The level of expression of genes across each timepoint.

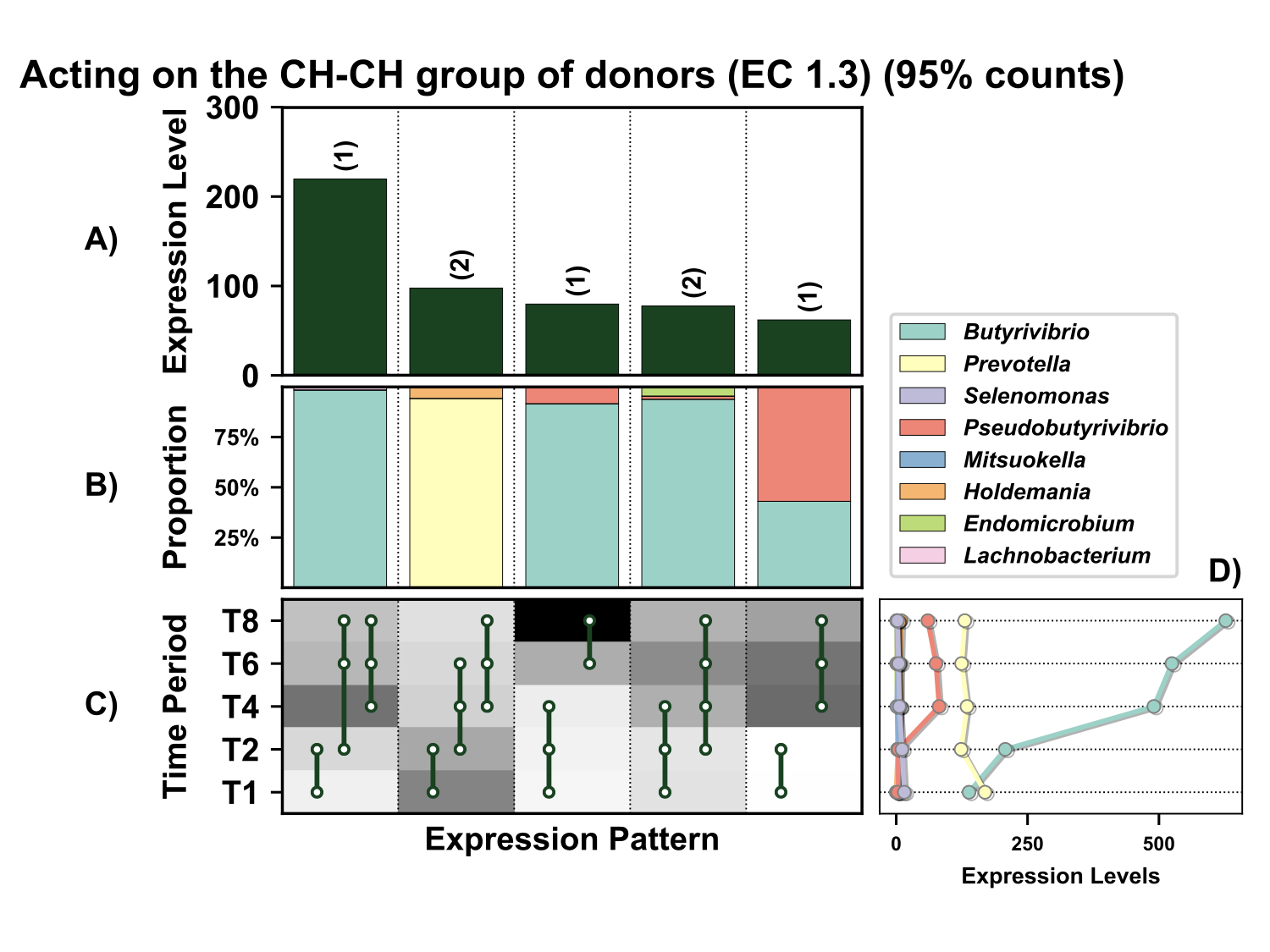

**Supplementary Figure 5.** Temporal expression of the top 95% most highly expressed genes acting on the CH-CH group of donors (EC 1.3) with a significant interaction with time expressed by prokaryotes attached to fresh perennial ryegrass incubated *in situ* within the rumen. Each column represents a set of genes that showed the same differential expression (DE) pattern (denoted as expression pattern on x axis). A) Summed expression of all genes with the same DE pattern, in brackets the number of genes with the same DE pattern. B) The proportion of taxonomic genera contributing to the expression level for each DE pattern. C) Visual representation of the DE patterns for each set of genes across the timepoints sampled; The heatmap represents the level of expression for each timepoint (Low = White, High = Black); The lines and dots represent the specific DE pattern shared by all genes in this set where the timepoints connected by line and dots were not significantly different from each other. D) The level of expression of genes across each timepoint.

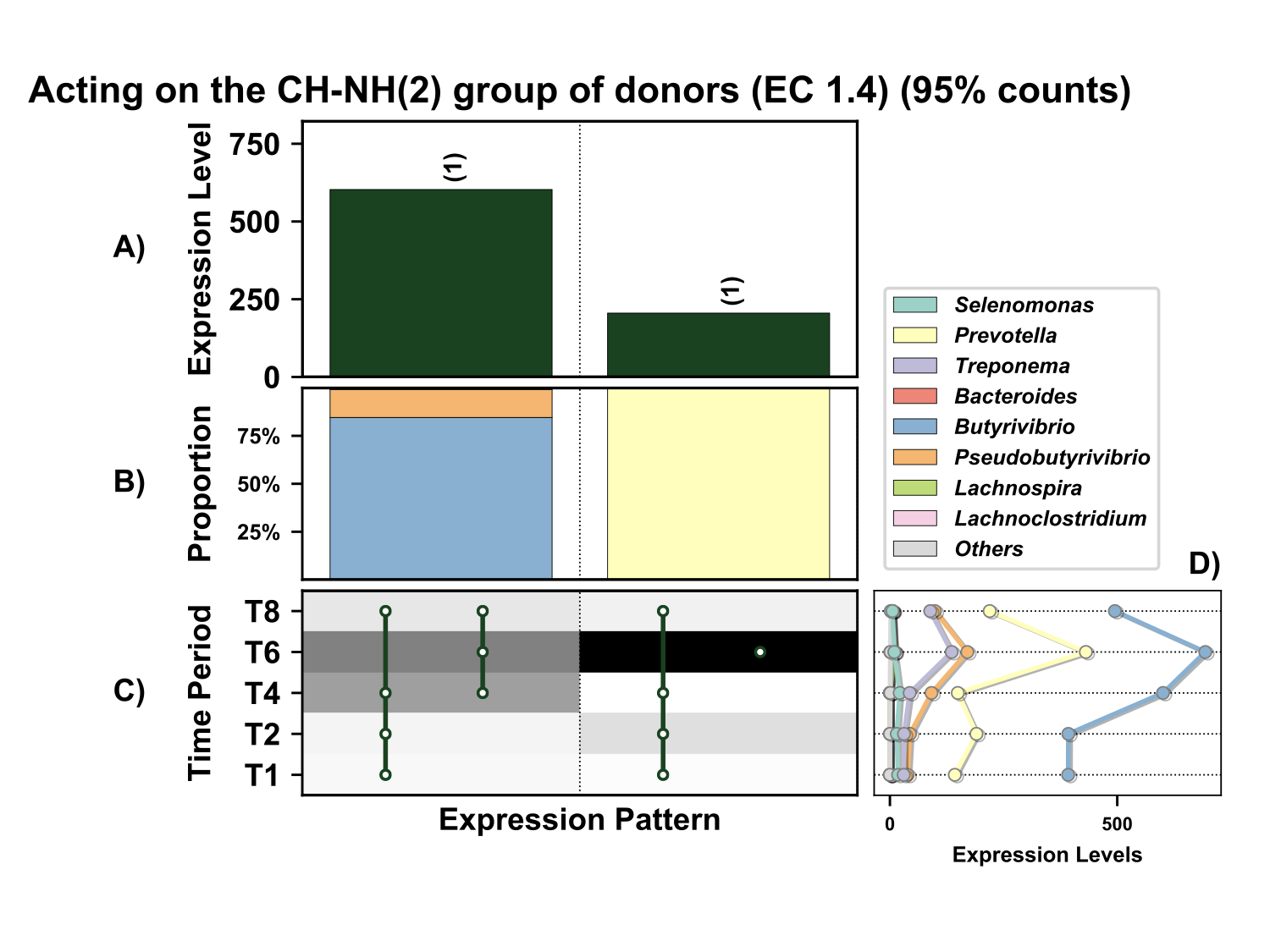
**Figure 6.** Temporal expression the top 95% most highly expressed genes acting on the CH-NH_2_ group of donors (EC 1.4) with a significant interaction with time expressed by prokaryotes attached to fresh perennial ryegrass incubated *in situ* within the rumen. Each column represents a set of genes that showed the same differential expression (DE) pattern (denoted as expression pattern on x axis). A) Summed expression of all genes with the same DE pattern, in brackets the number of genes with the same DE pattern. B) The proportion of taxonomic genera contributing to the expression level for each DE pattern. C) Visual representation of the DE patterns for each set of genes across the timepoints sampled; The heatmap represents the level of expression for each timepoint (Low = White, High = Black); The lines and dots represent the specific DE pattern shared by all genes in this set where the timepoints connected by line and dots were not significantly different from each other. D) The level of expression of genes across each timepoint.

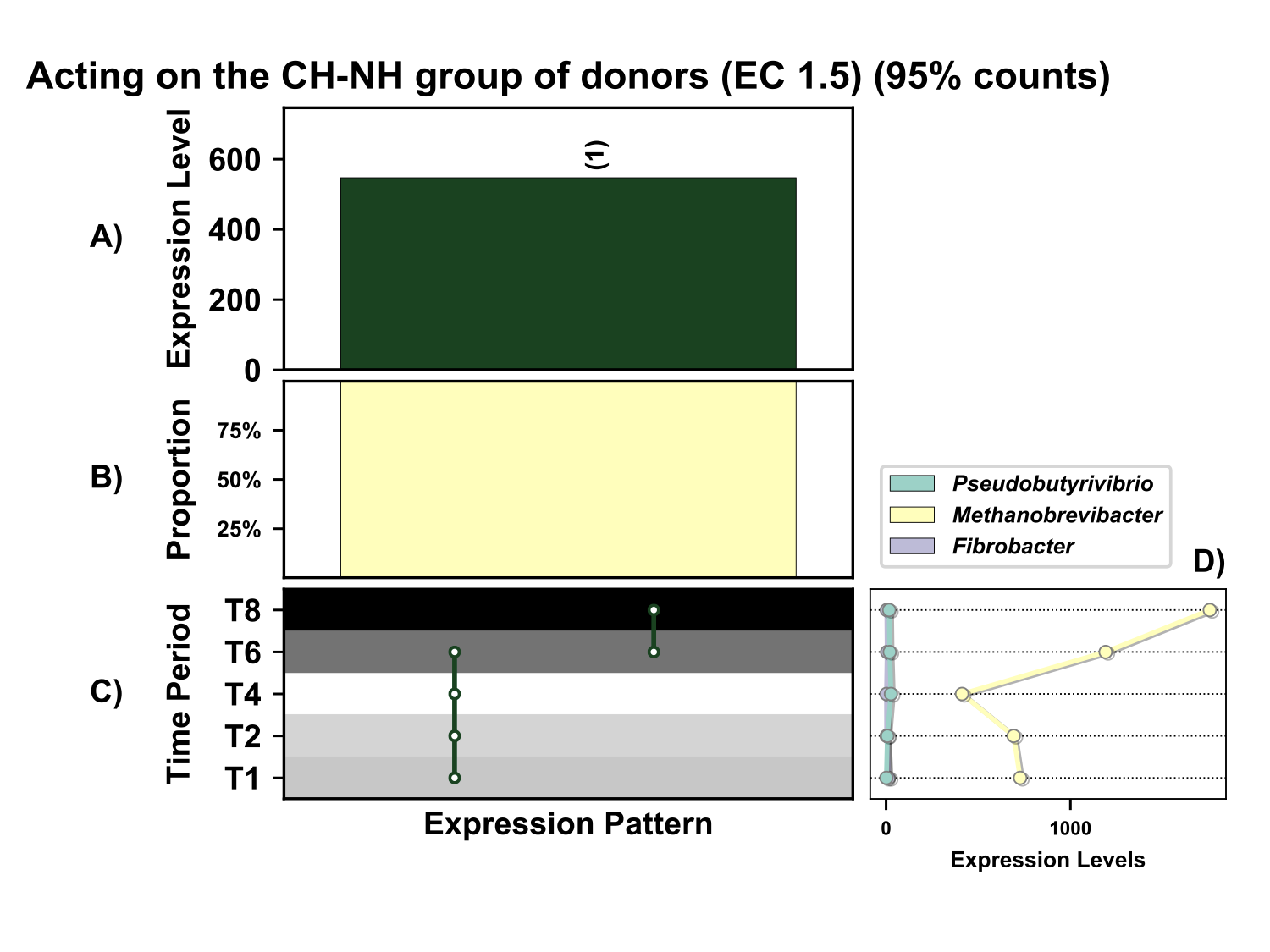
**Supplementary Figure 7.** Temporal expression of genes the top 95% most highly expressed genes acting on the CH-NH group of donors (EC 1.5) with a significant interaction with time expressed by prokaryotes attached to fresh perennial ryegrass incubated *in situ* within the rumen. Each column represents a set of genes that showed the same differential expression (DE) pattern (denoted as expression pattern on x axis). A) Summed expression of all glycosylases with the same DE pattern, in brackets the number of genes with the same DE pattern. B) The proportion of taxonomic genera contributing to the expression level for each DE pattern. C) Visual representation of the DE patterns for each set of genes across the timepoints sampled; The heatmap represents the level of expression for each timepoint (Low = White, High = Black); The lines and dots represent the specific DE pattern shared by all genes in this set where the timepoints connected by line and dots were not significantly different from each other. D) The level of expression of genes across each timepoint.

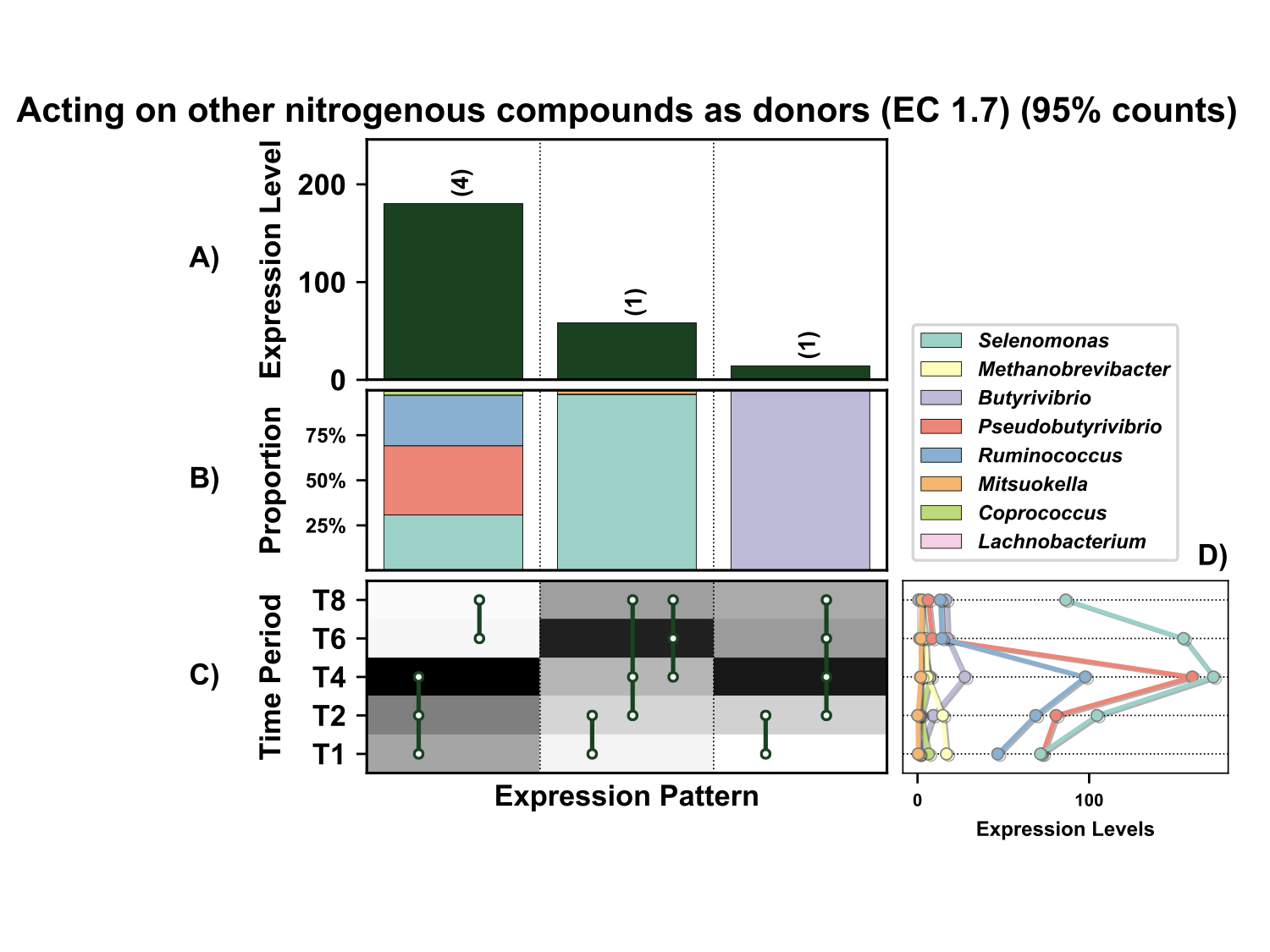
**Supplementary Figure 8.** Temporal expression of the top 95% most highly expressed genes acting on other nitrogenous compounds as donors (EC 1.7) with a significant interaction with time expressed by prokaryotes attached to fresh perennial ryegrass incubated *in situ* within the rumen. Each column represents a set of genes that showed the same differential expression (DE) pattern (denoted as expression pattern on x axis). A) Summed expression of all genes with the same DE pattern, in brackets the number of genes with the same DE pattern. B) The proportion of taxonomic genera contributing to the expression level for each DE pattern. C) Visual representation of the DE patterns for each set of genes across the timepoints sampled; The heatmap represents the level of expression for each timepoint (Low = White, High = Black); The lines and dots represent the specific DE pattern shared by all genes in this set where the timepoints connected by line and dots were not significantly different from each other. D) The level of expression of genes across each timepoint.

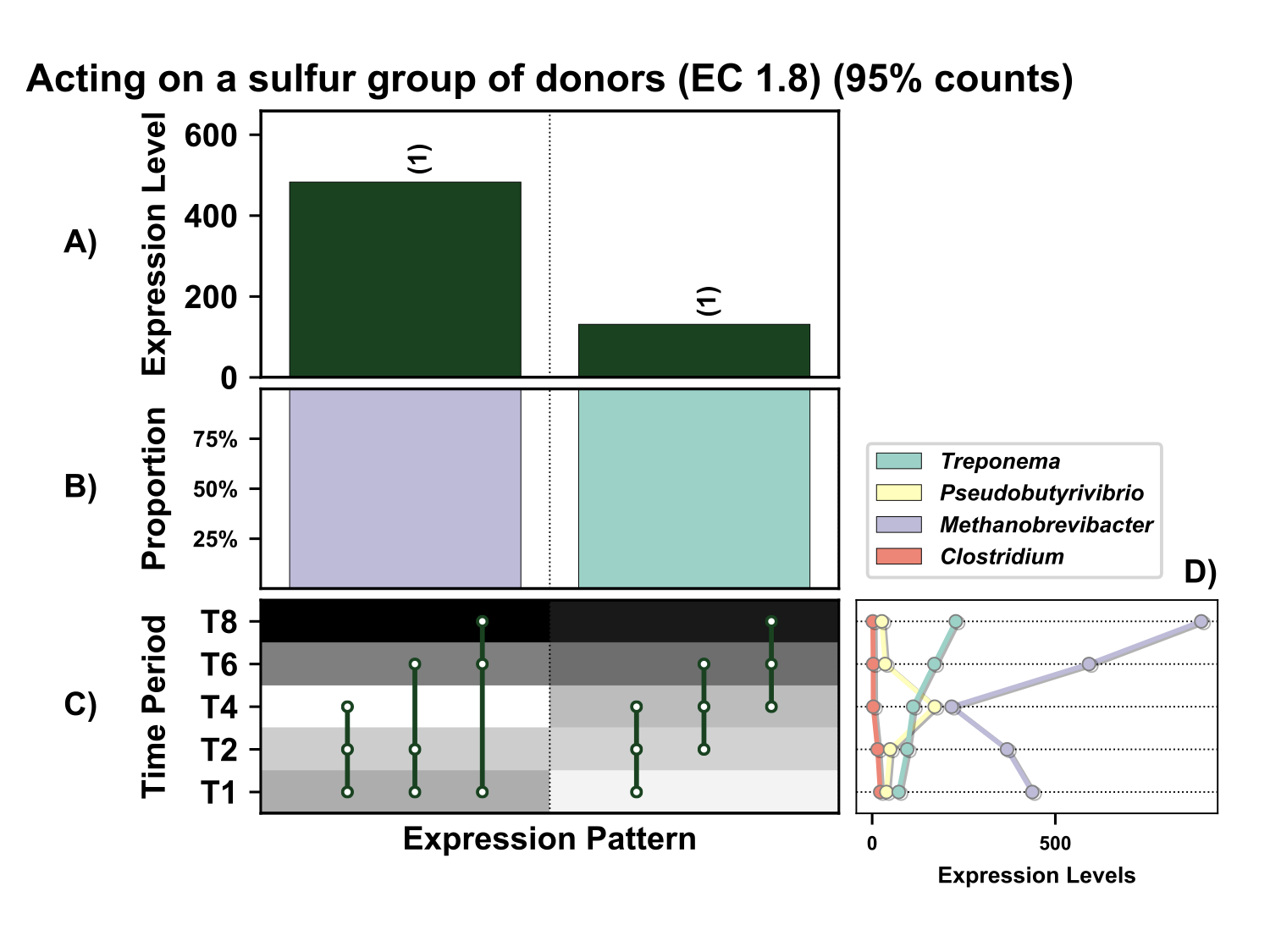
**Supplementary Figure 9.** Temporal expression of the top 95% most highly expressed genes acting on a sulfur group of donors (EC 1.8) with a significant interaction with time expressed by prokaryotes attached to fresh perennial ryegrass incubated *in situ* within the rumen. Each column represents a set of genes that showed the same differential expression (DE) pattern. A) Summed expression of all genes with the same DE pattern, in brackets the number of genes with the same DE pattern. B) The proportion of taxonomic genera contributing to the expression level for each DE pattern. C) Visual representation of the DE patterns for each set of genes across the timepoints sampled; The heatmap represents the level of expression for each timepoint (Low = White, High = Black); The lines and dots represent the specific DE pattern shared by all genes in this set where the timepoints connected by line and dots were not significantly different from each other. D) The level of expression of genes across each timepoint.

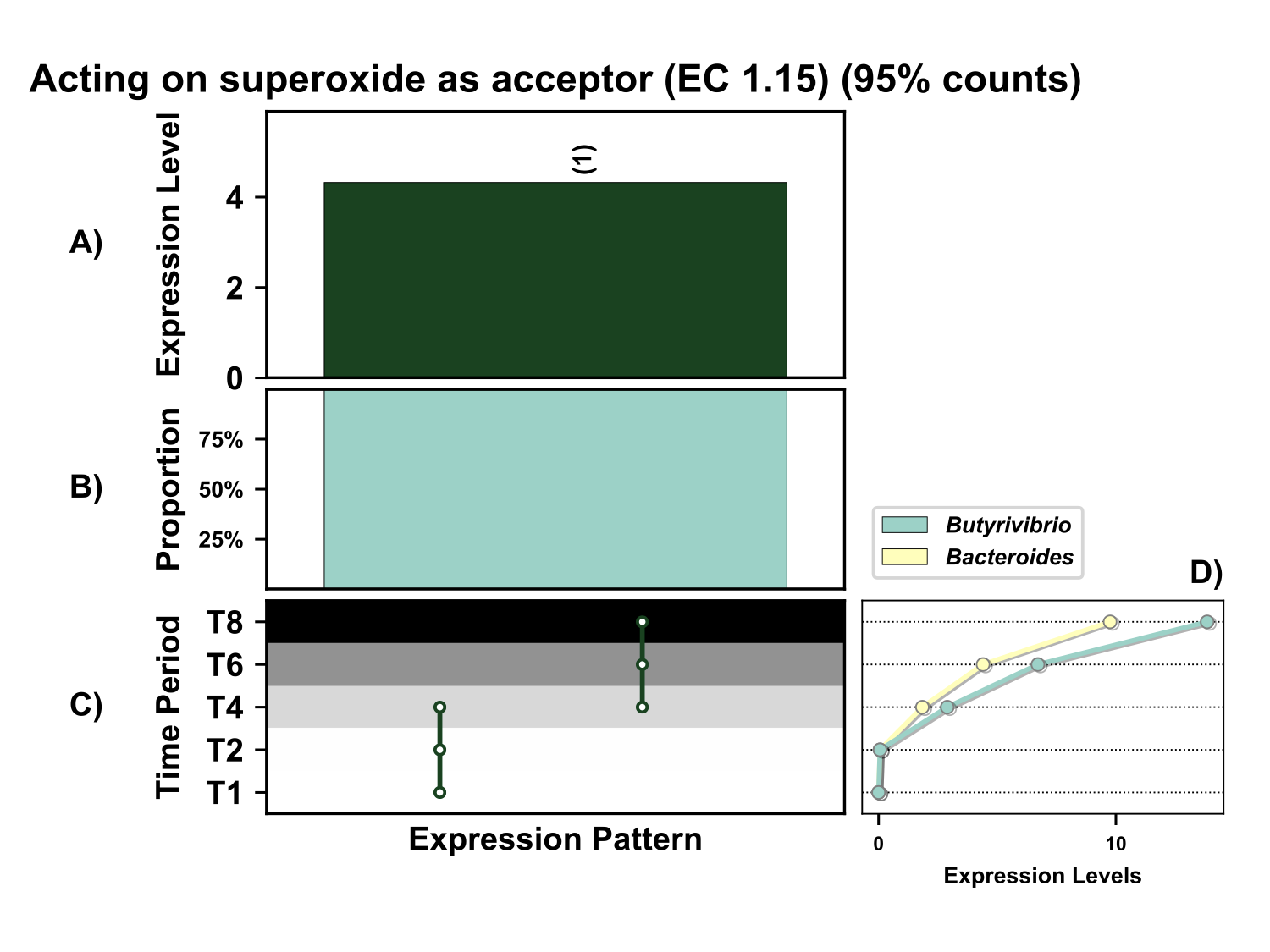

**Supplementary Figure 10.** Temporal expression of the top 95% most highly expressed genes acting on superoxide radicals as acceptors (EC 1.15) with a significant interaction with time expressed by prokaryotes attached to fresh perennial ryegrass incubated *in situ* within the rumen. Each column represents a set of genes that showed the same differential expression (DE) pattern (denoted as expression pattern on x axis). A) Summed expression of all genes with the same DE pattern, in brackets the number of genes with the same DE pattern. B) The proportion of taxonomic genera contributing to the expression level for each DE pattern. C) Visual representation of the DE patterns for each set of genes across the timepoints sampled; The heatmap represents the level of expression for each timepoint (Low = White, High = Black); The lines and dots represent the specific DE pattern shared by all genes in this set where the timepoints connected by line and dots were not significantly different from each other. D) The level of expression of genes across each timepoint.

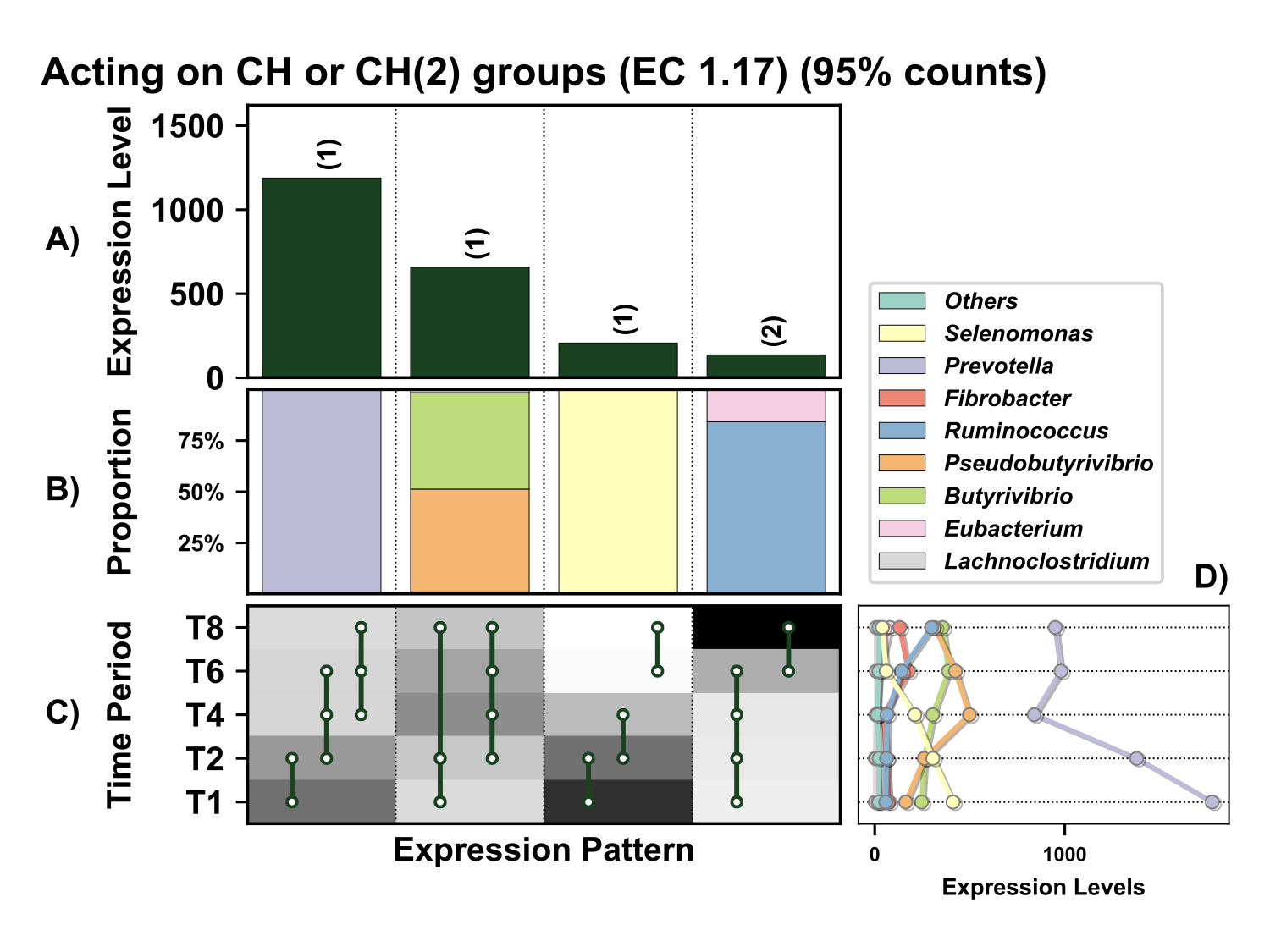

**Supplementary Figure 11.** Temporal expression of the top 95% most highly expressed genes acting on the CH or CH_2_ groups (EC 1.17) with a significant interaction with time expressed by prokaryotes attached to fresh perennial ryegrass incubated *in situ* within the rumen. Each column represents a set of genes that showed the same differential expression (DE) pattern (denoted as expression pattern on x axis). A) Summed expression of all genes with the same DE pattern, in brackets the number of genes with the same DE pattern. B) The proportion of taxonomic genera contributing to the expression level for each DE pattern. C) Visual representation of the DE patterns for each set of genes across the timepoints sampled; The heatmap represents the level of expression for each timepoint (Low = White, High = Black); The lines and dots represent the specific DE pattern shared by all genes in this set where the timepoints connected by line and dots were not significantly different from each other. D) The level of expression of genes across each timepoint.

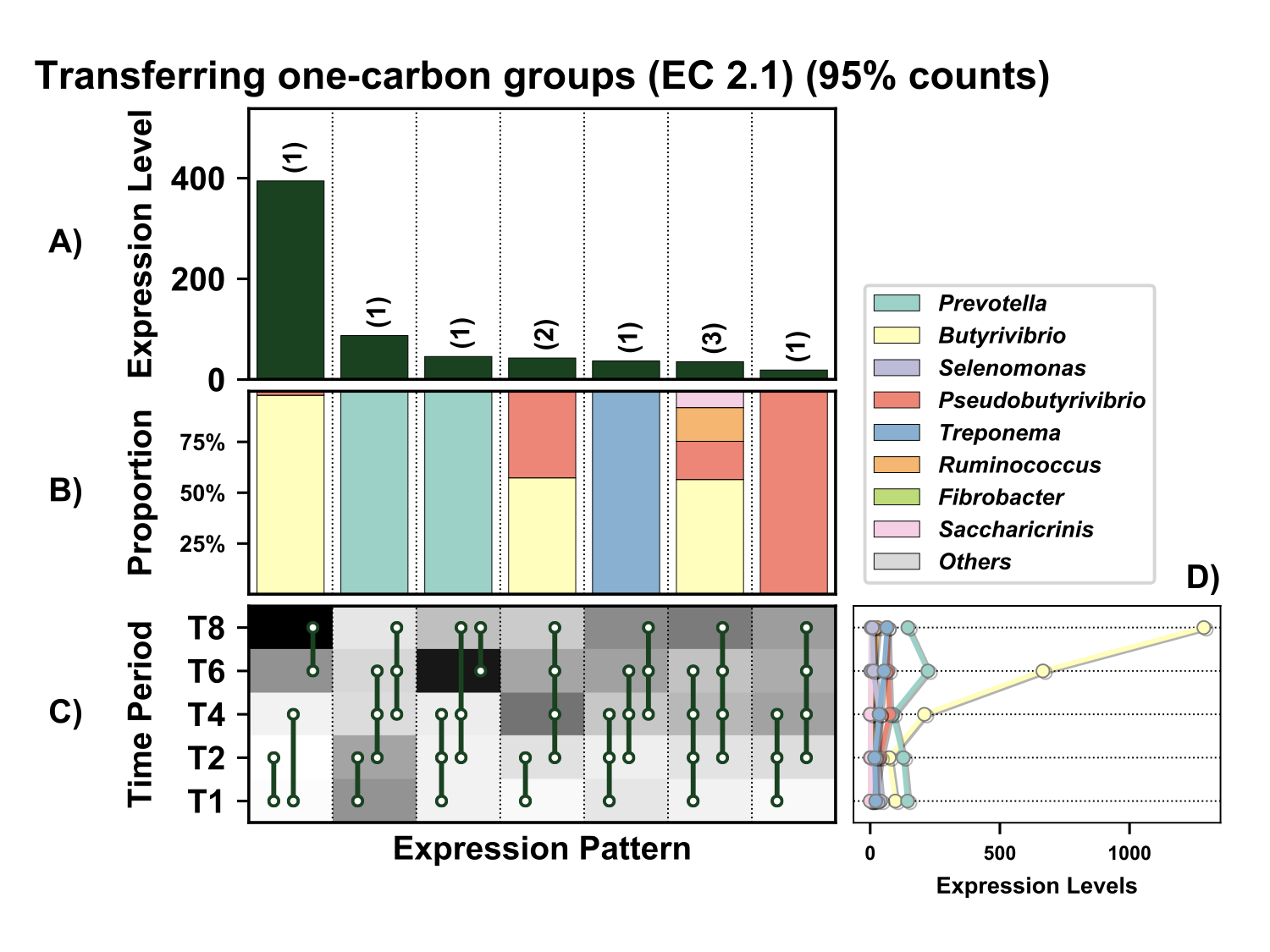

**Supplementary Figure 12.** Temporal expression of the top 95% most highly expressed genes involved in transfer one-carbon groups, Methylase (EC 2.1) with a significant interaction with time expressed by prokaryotes attached to fresh perennial ryegrass incubated *in situ* within the rumen. Each column represents a set of genes that showed the same differential expression (DE) pattern (denoted as expression pattern on x axis). A) Summed expression of all genes with the same DE pattern, in brackets the number of genes with the same DE pattern. B) The proportion of taxonomic genera contributing to the expression level for each DE pattern. C) Visual representation of the DE patterns for each set of genes across the timepoints sampled; The heatmap represents the level of expression for each timepoint (Low = White, High = Black); The lines and dots represent the specific DE pattern shared by all genes in this set where the timepoints connected by line and dots were not significantly different from each other. D) The level of expression of genes across each timepoint.

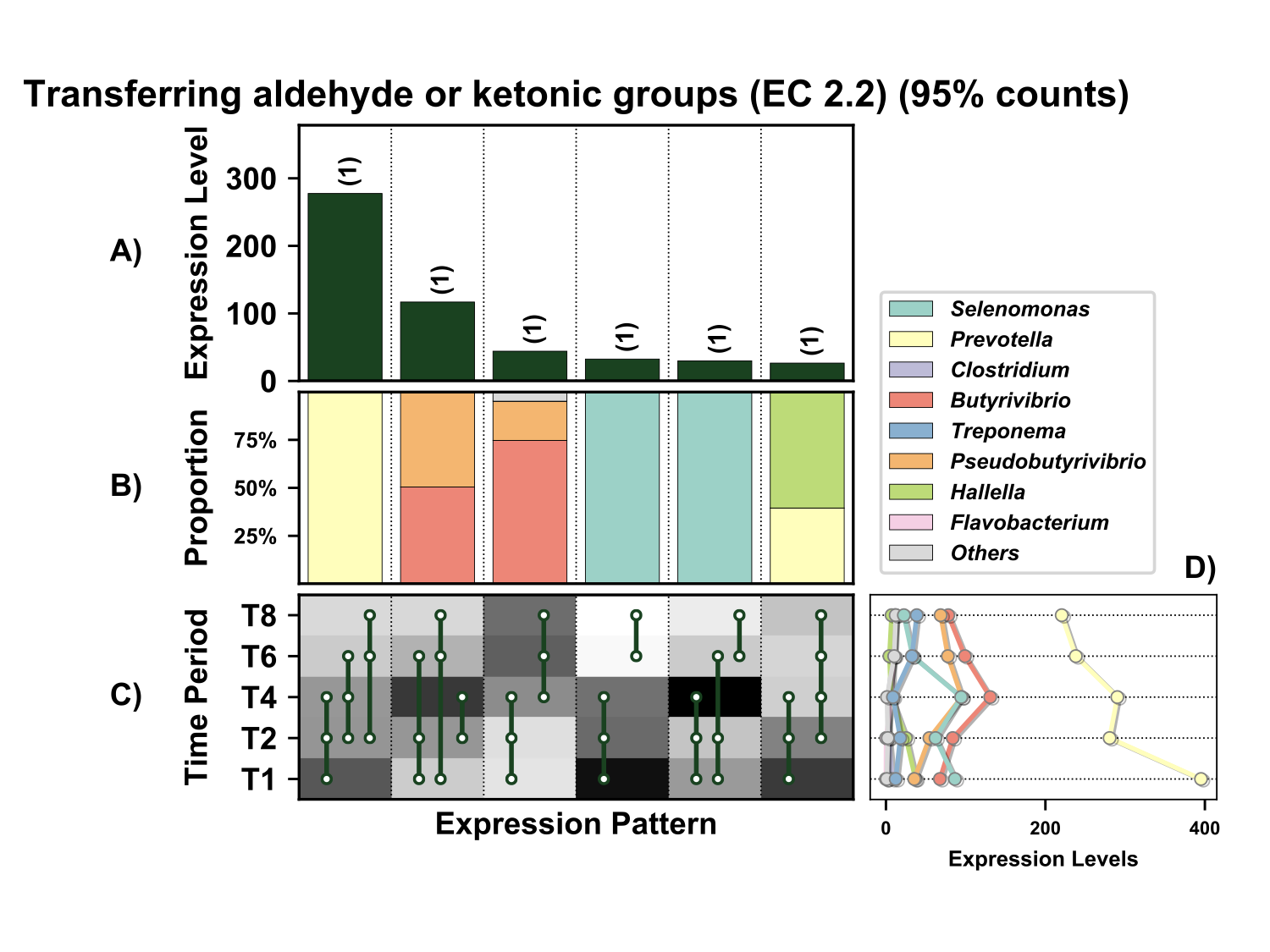

**Supplementary Figure 13.** Temporal expression of the top 95% most highly expressed genes involved in transfer [aldehyde](https://en.wikipedia.org/wiki/Aldehyde) or [ketone](https://en.wikipedia.org/wiki/Ketone) groups (EC 2.2) with a significant interaction with time expressed by prokaryotes attached to fresh perennial ryegrass incubated *in situ* within the rumen. Each column represents a set of genes that showed the same differential expression (DE) pattern (denoted as expression pattern on x axis). A) Summed expression of all genes with the same DE pattern, in brackets the number of genes with the same DE pattern. B) The proportion of taxonomic genera contributing to the expression level for each DE pattern. C) Visual representation of the DE patterns for each set of genes across the timepoints sampled; The heatmap represents the level of expression for each timepoint (Low = White, High = Black); The lines and dots represent the specific DE pattern shared by all genes in this set where the timepoints connected by line and dots were not significantly different from each other. D) The level of expression of genes across each timepoint.

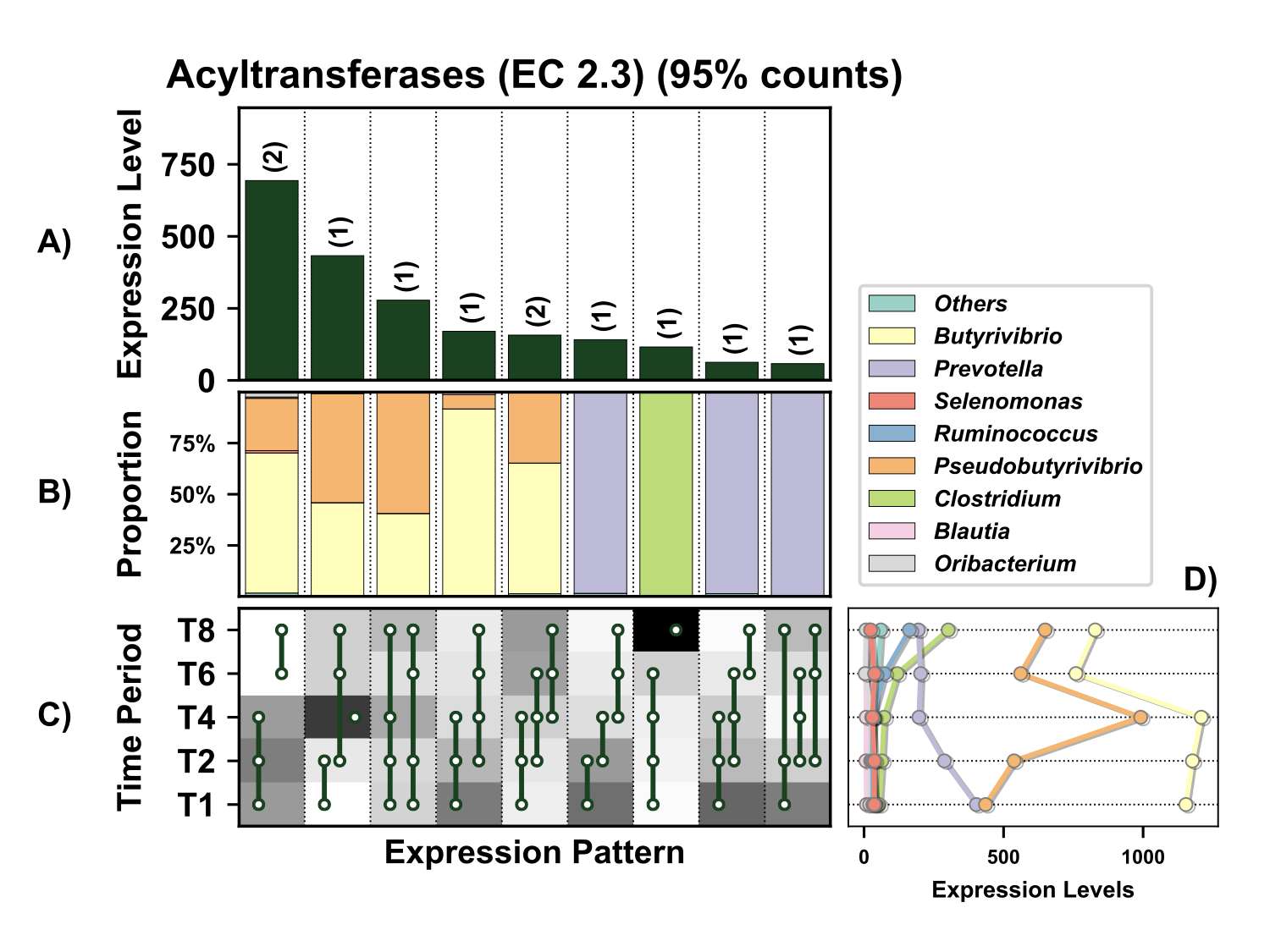

**Supplementary Figure 14.** Temporal expression of the top 95% most highly expressed acyltransferases (EC 2.3) with a significant interaction with time expressed by prokaryotes attached to fresh perennial ryegrass incubated *in situ* within the rumen. Each column represents a set of genes that showed the same differential expression (DE) pattern (denoted as expression pattern on x axis). A) Summed expression of all genes with the same DE pattern, in brackets the number of genes with the same DE pattern. B) The proportion of taxonomic genera contributing to the expression level for each DE pattern. C) Visual representation of the DE patterns for each set of genes across the timepoints sampled; The heatmap represents the level of expression for each timepoint (Low = White, High = Black); The lines and dots represent the specific DE pattern shared by all genes in this set where the timepoints connected by line and dots were not significantly different from each other. D) The level of expression across each timepoint.

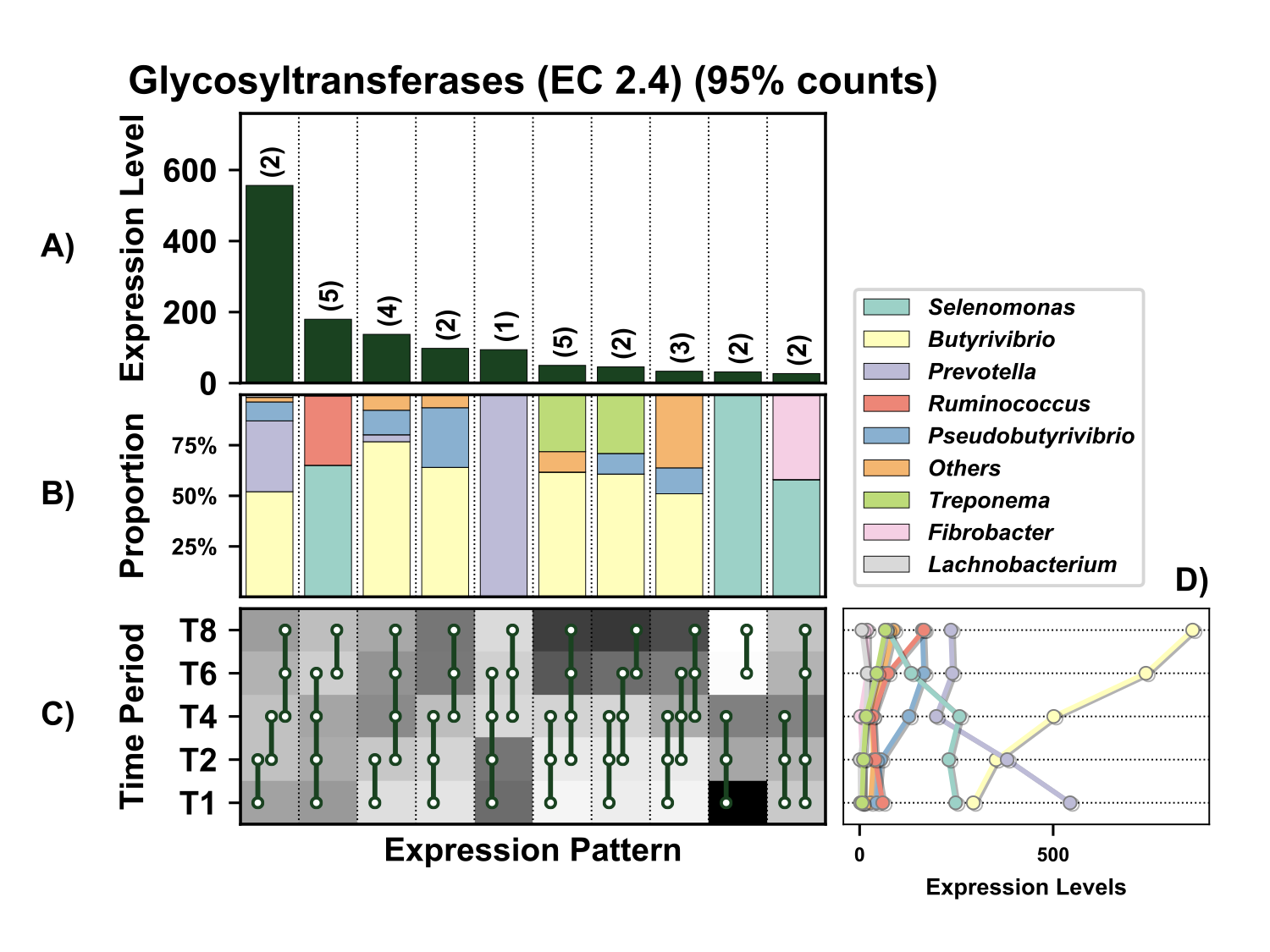

**Supplementary Figure 15.** Temporal expression of the top 95% most highly expressed glycosyltransferases (EC 2.4) with a significant interaction with time expressed by prokaryotes attached to fresh perennial ryegrass incubated *in situ* within the rumen. Each column represents a set of genes that showed the same differential expression (DE) pattern (denoted as expression pattern on x axis). A) Summed expression of all genes with the same DE pattern, in brackets the number of genes with the same DE pattern. B) The proportion of taxonomic genera contributing to the expression level for each DE pattern. C) Visual representation of the DE patterns for each set of genes across the timepoints sampled; The heatmap represents the level of expression for each timepoint (Low = White, High = Black); The lines and dots represent the specific DE pattern shared by all genes in this set where the timepoints connected by line and dots were not significantly different from each other. D) The level of expression of genes across each timepoint.

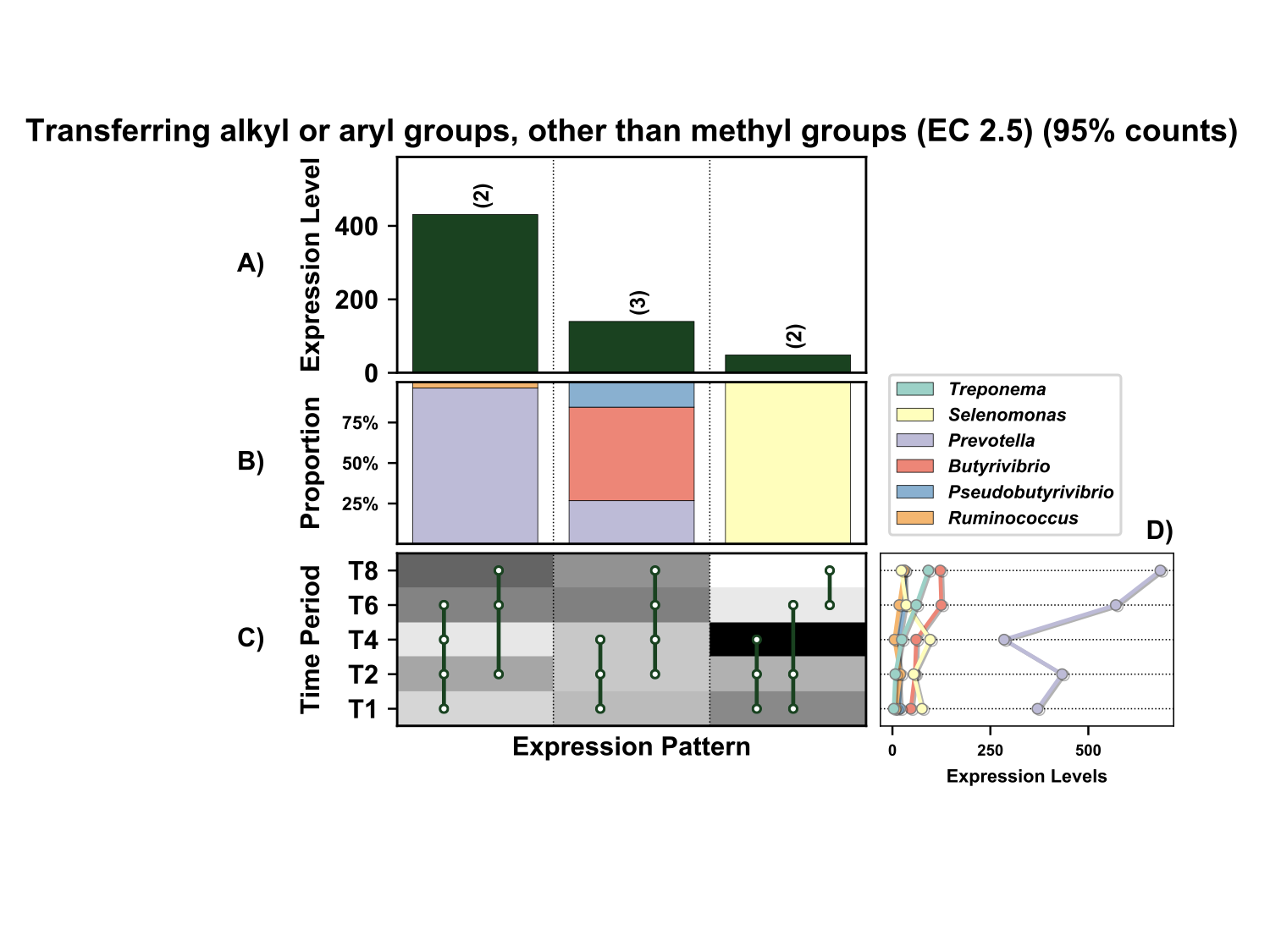
**Supplementary Figure 16.** Temporal expression of the top 95% most highly expressed genes acting on transfer [alkyl](https://en.wikipedia.org/wiki/Alkyl) or [aryl](https://en.wikipedia.org/wiki/Aryl) groups, other than [methyl](https://en.wikipedia.org/wiki/Methyl) groups (EC 2.5) with a significant interaction with time expressed by prokaryotes attached to fresh perennial ryegrass incubated *in situ* within the rumen. Each column represents a set of genes that showed the same differential expression (DE) pattern (denoted as expression pattern on x axis). A) Summed expression of all genes with the same DE pattern, in brackets the number of genes with the same DE pattern. B) The proportion of taxonomic genera contributing to the expression level for each DE pattern. C) Visual representation of the DE patterns for each set of genes acting on transfer [alkyl](https://en.wikipedia.org/wiki/Alkyl) or [aryl](https://en.wikipedia.org/wiki/Aryl) groups, other than [methyl](https://en.wikipedia.org/wiki/Methyl) groups across the timepoints sampled; The heatmap represents the level of expression for each timepoint (Low = White, High = Black); The lines and dots represent the specific DE pattern shared by all genes in this set where the timepoints connected by line and dots were not significantly different from each other. D) The level of expression of genes across each timepoint.

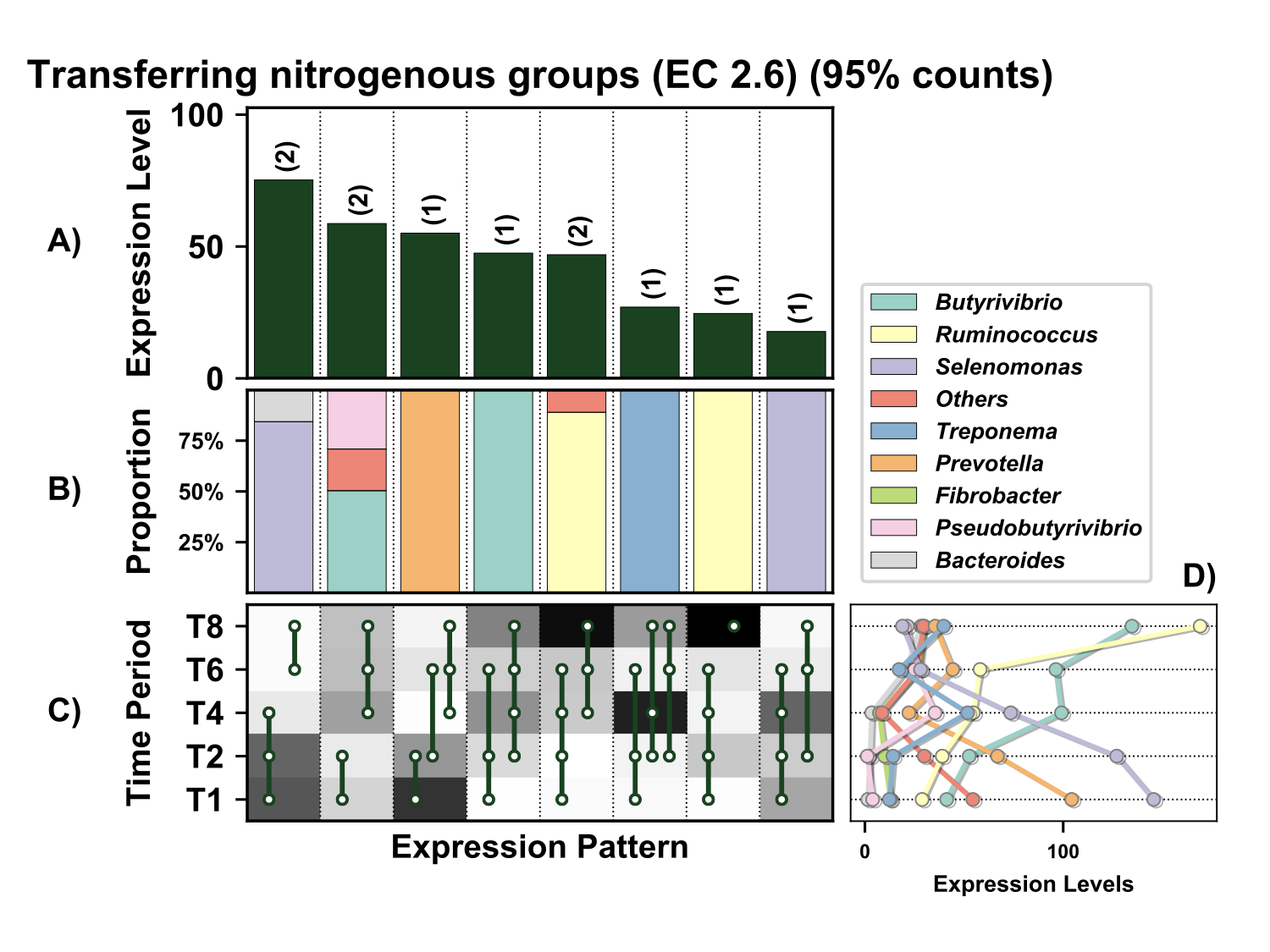

**Supplementary Figure 17.** Temporal expression of the top 95% most highly expressed genes involved in transfer of nitrogenous groups (EC 2.6) with a significant interaction with time expressed by prokaryotes attached to fresh perennial ryegrass incubated *in situ* within the rumen. Each column represents a set of genes that showed the same differential expression (DE) pattern (denoted as expression pattern on x axis). A) Summed expression of all genes with the same DE pattern, in brackets the number of genes with the same DE pattern. B) The proportion of taxonomic genera contributing to the expression level for each DE pattern. C) Visual representation of the DE patterns for each set of genes across the timepoints sampled; The heatmap represents the level of expression for each timepoint (Low = White, High = Black); The lines and dots represent the specific DE pattern shared by all genes in this set where the timepoints connected by line and dots were not significantly different from each other. D) The level of expression of genes across each timepoint.

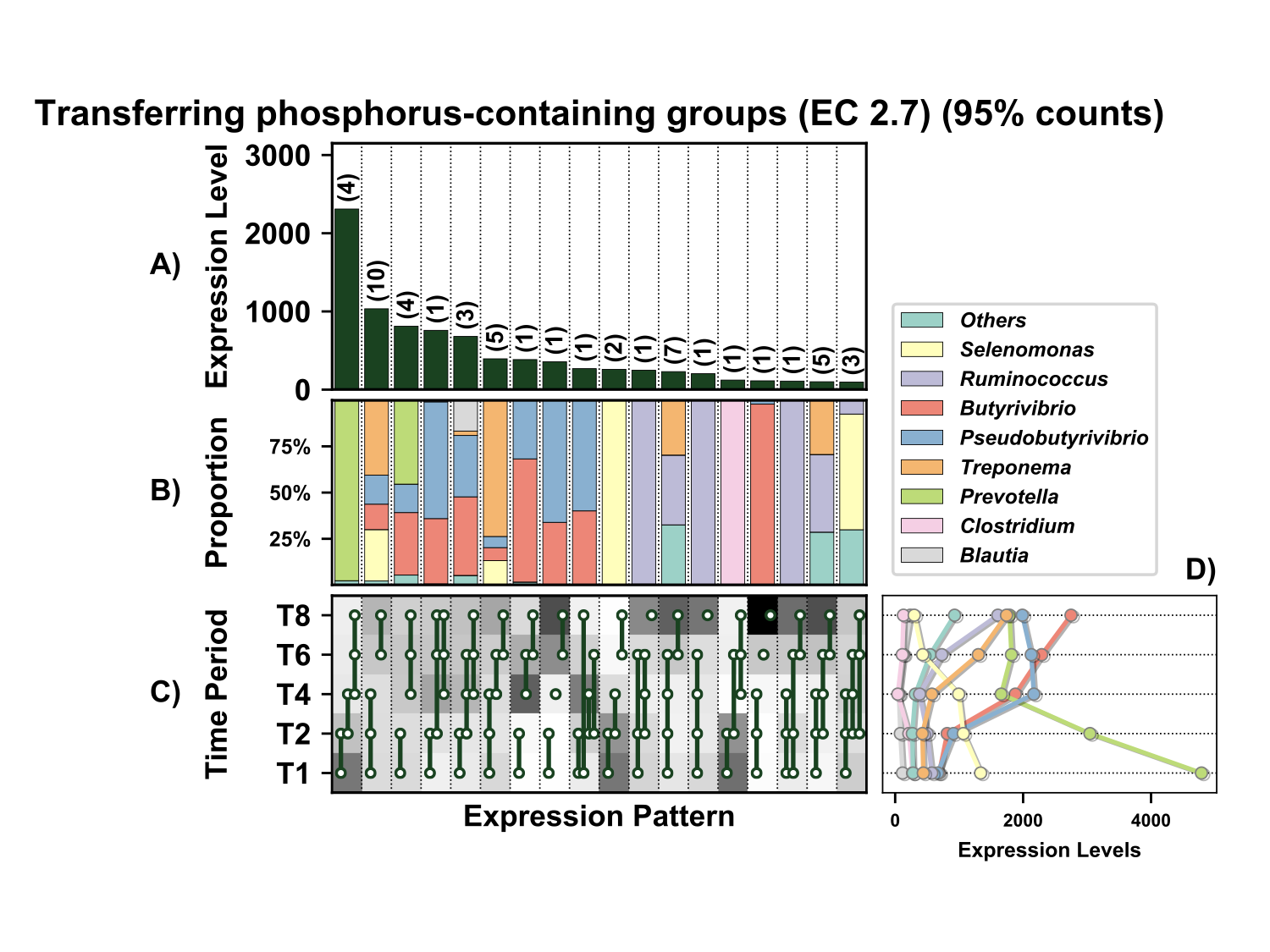

**Supplementary Figure 18.** Temporal expression of the top 95% most highly expressed genes involved in transfer of phosphorus-containing groups (EC 2.7) with a significant interaction with time expressed by prokaryotes attached to fresh perennial ryegrass incubated *in situ* within the rumen. Each column represents a set of genes that showed the same differential expression (DE) pattern (denoted as expression pattern on x axis). A) Summed expression of all genes involved with the same DE pattern, in brackets the number of genes with the same DE pattern. B) The proportion of taxonomic genera contributing to the expression level for each DE pattern. C) Visual representation of the DE patterns for each set of genes across the timepoints sampled; The heatmap represents the level of expression for each timepoint (Low = White, High = Black); The lines and dots represent the specific DE pattern shared by all genes in this set where the timepoints connected by line and dots were not significantly different from each other. D) The level of expression of genes across each timepoint.

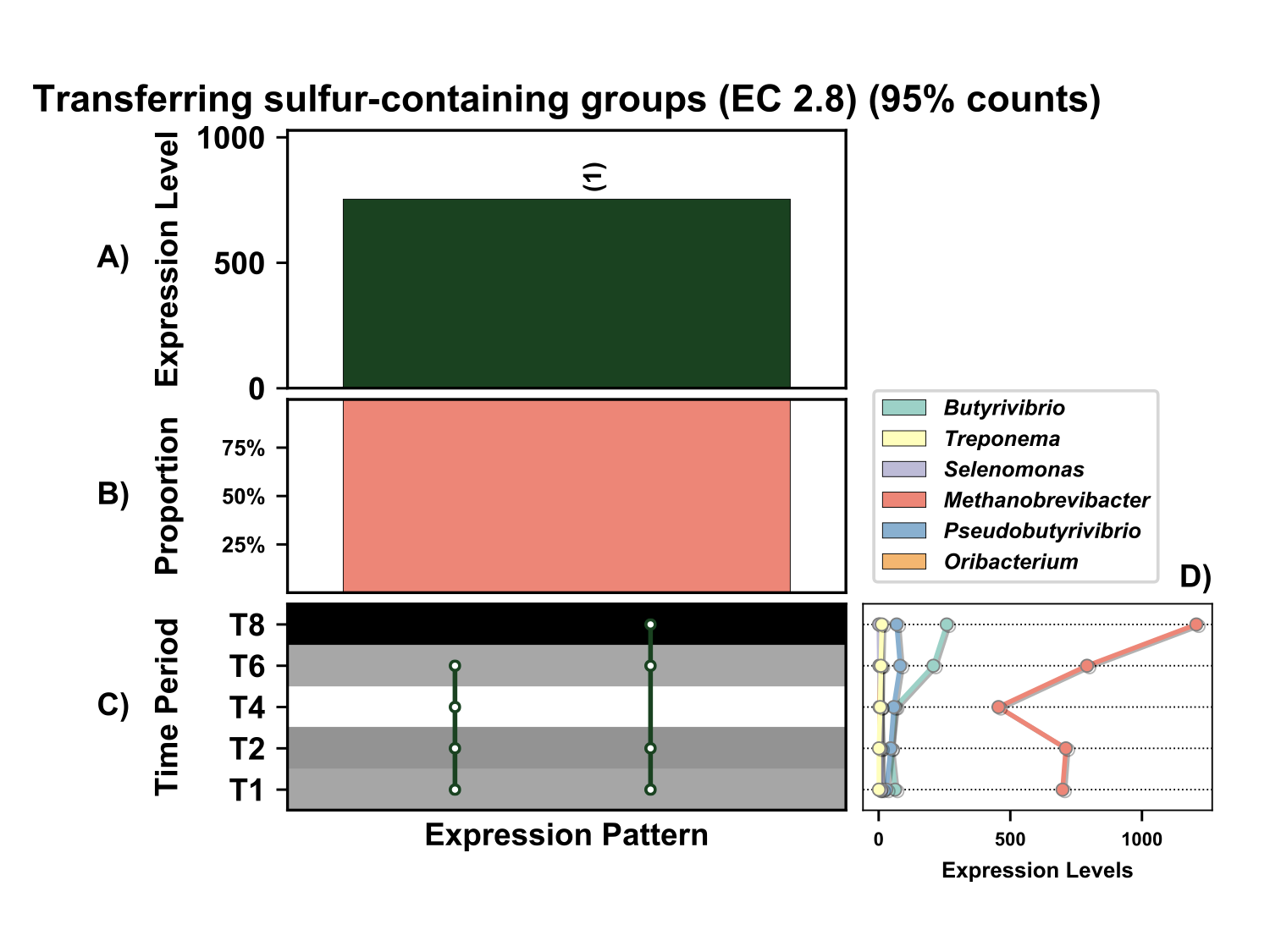

**Supplementary Figure 19.** Temporal expression of the top 95% most highly expressed genes involved in transfer of transfer sulfur-containing groups (EC 2.8) with a significant interaction with time expressed by prokaryotes attached to fresh perennial ryegrass incubated *in situ* within the rumen. Each column represents a set of genes that showed the same differential expression (DE) pattern (denoted as expression pattern on x axis). A) Summed expression of all genes with the same DE pattern, in brackets the number of genes with the same DE pattern. B) The proportion of taxonomic genera contributing to the expression level for each DE pattern. C) Visual representation of the DE patterns for each set of genes across the timepoints sampled; The heatmap represents the level of expression for each timepoint (Low = White, High = Black); The lines and dots represent the specific DE pattern shared by all genes in this set where the timepoints connected by line and dots were not significantly different from each other. D) The level of expression of genes across each timepoint.

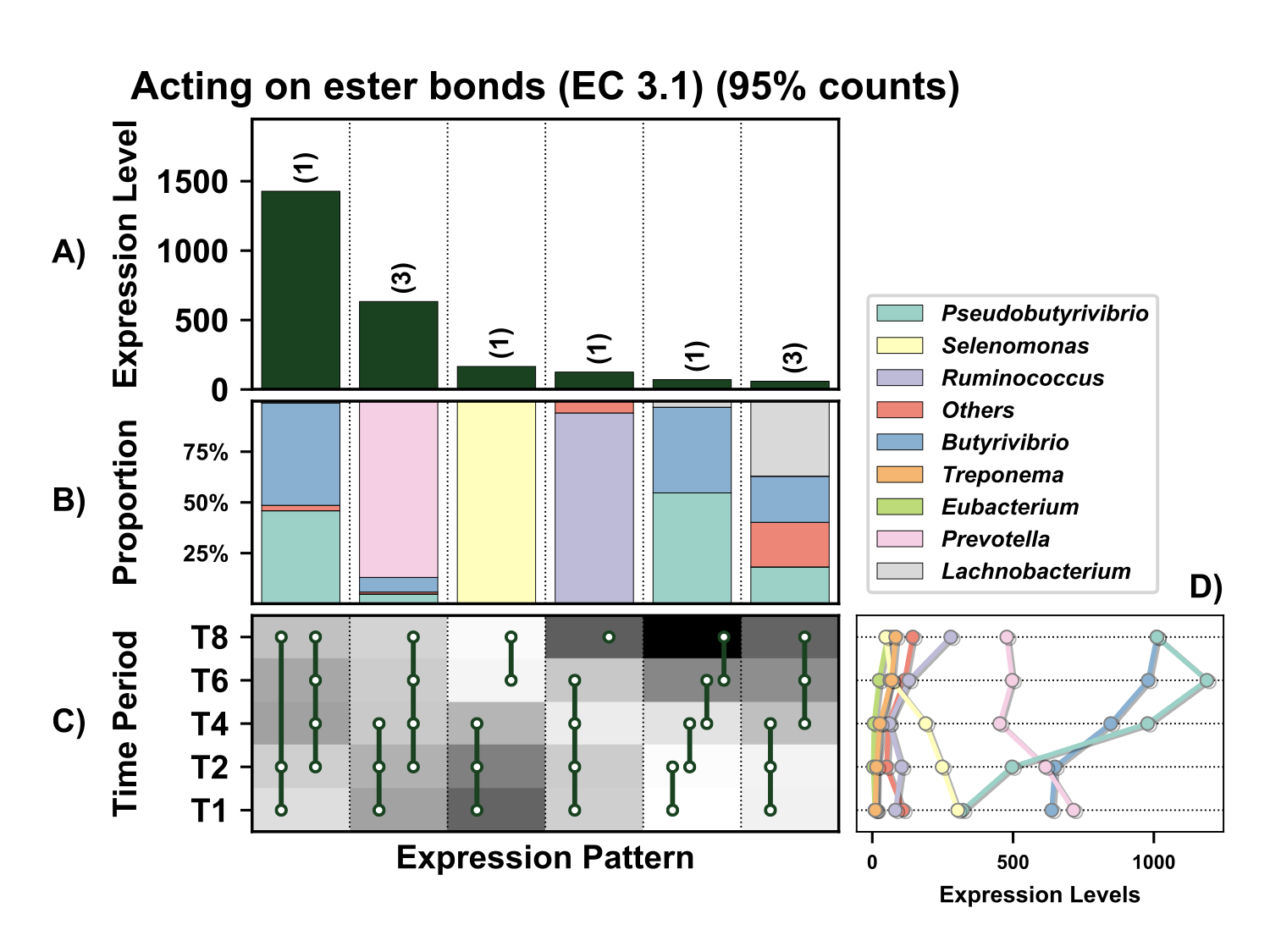

**Supplementary Figure 20.** Temporal expression of the top 95% most highly expressed genes acting on [ester](https://en.wikipedia.org/wiki/Ester) bonds (EC 3.1) with a significant interaction with time expressed by prokaryotes attached to fresh perennial ryegrass incubated *in situ* within the rumen. Each column represents a set of genes that showed the same differential expression (DE) pattern (denoted as expression pattern on x axis). A) Summed expression of all genes with the same DE pattern, in brackets the number of genes with the same DE pattern. B) The proportion of taxonomic genera contributing to the expression level for each DE pattern. C) Visual representation of the DE patterns for each set of genes across the timepoints sampled; The heatmap represents the level of expression for each timepoint (Low = White, High = Black); The lines and dots represent the specific DE pattern shared by all genes in this set where the timepoints connected by line and dots were not significantly different from each other. D) The level of expression of genes across each timepoint.

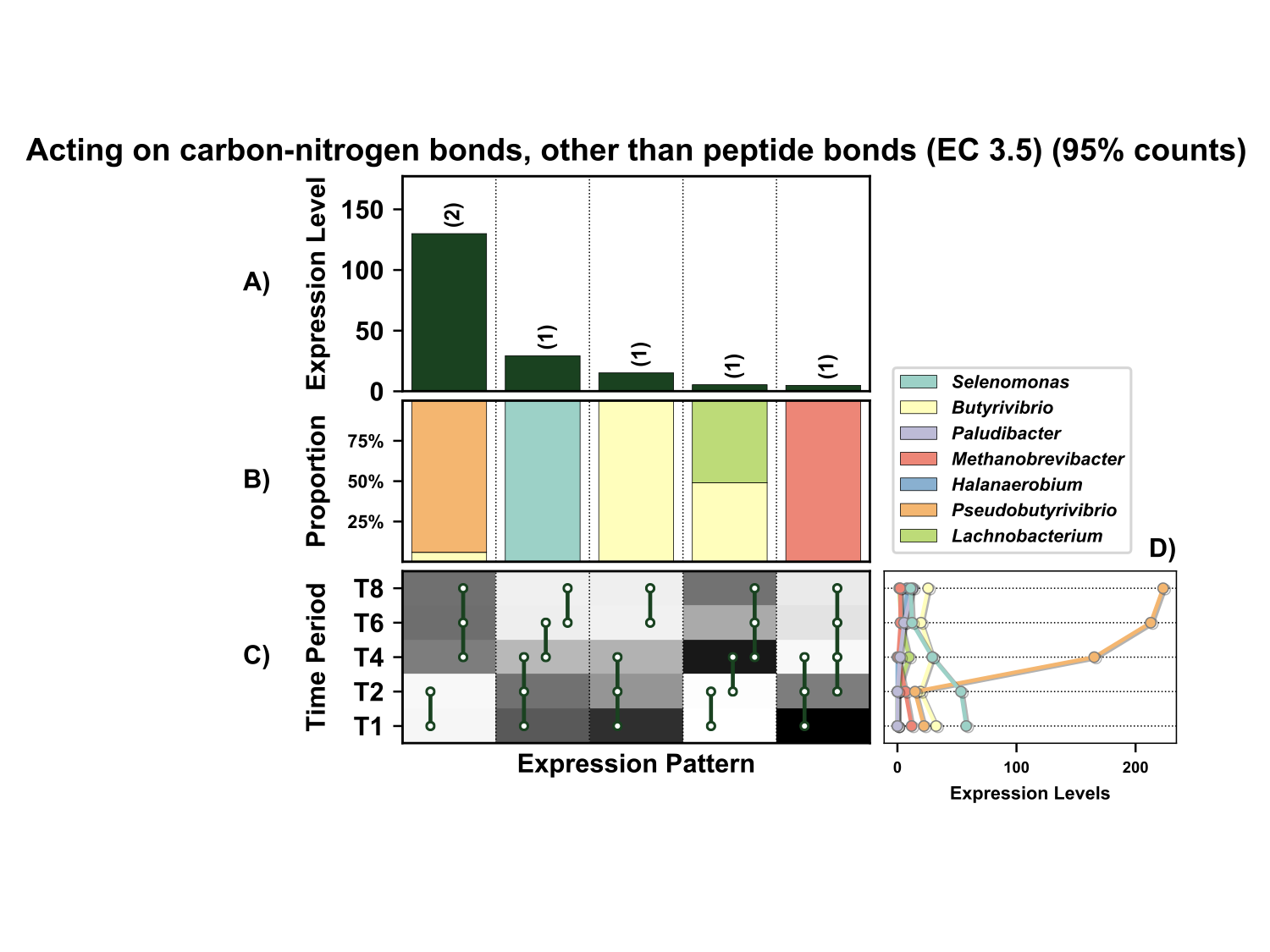

**Supplementary Figure 21.** Temporal expression of the top 95% most highly expressed genes acting on carbon-nitrogen bonds, other than peptide bonds (EC 3.5) with a significant interaction with time expressed by prokaryotes attached to fresh perennial ryegrass incubated *in situ* within the rumen. Each column represents a set of genes that showed the same differential expression (DE) pattern (denoted as expression pattern on x axis). A) Summed expression of all genes with the same DE pattern, in brackets the number of genes with the same DE pattern. B) The proportion of taxonomic genera contributing to the expression level for each DE pattern. C) Visual representation of the DE patterns for each set of genes across the timepoints sampled; The heatmap represents the level of expression for each timepoint (Low = White, High = Black); The lines and dots represent the specific DE pattern shared by all genes in this set where the timepoints connected by line and dots were not significantly different from each other. D) The level of expression of genes across each timepoint.

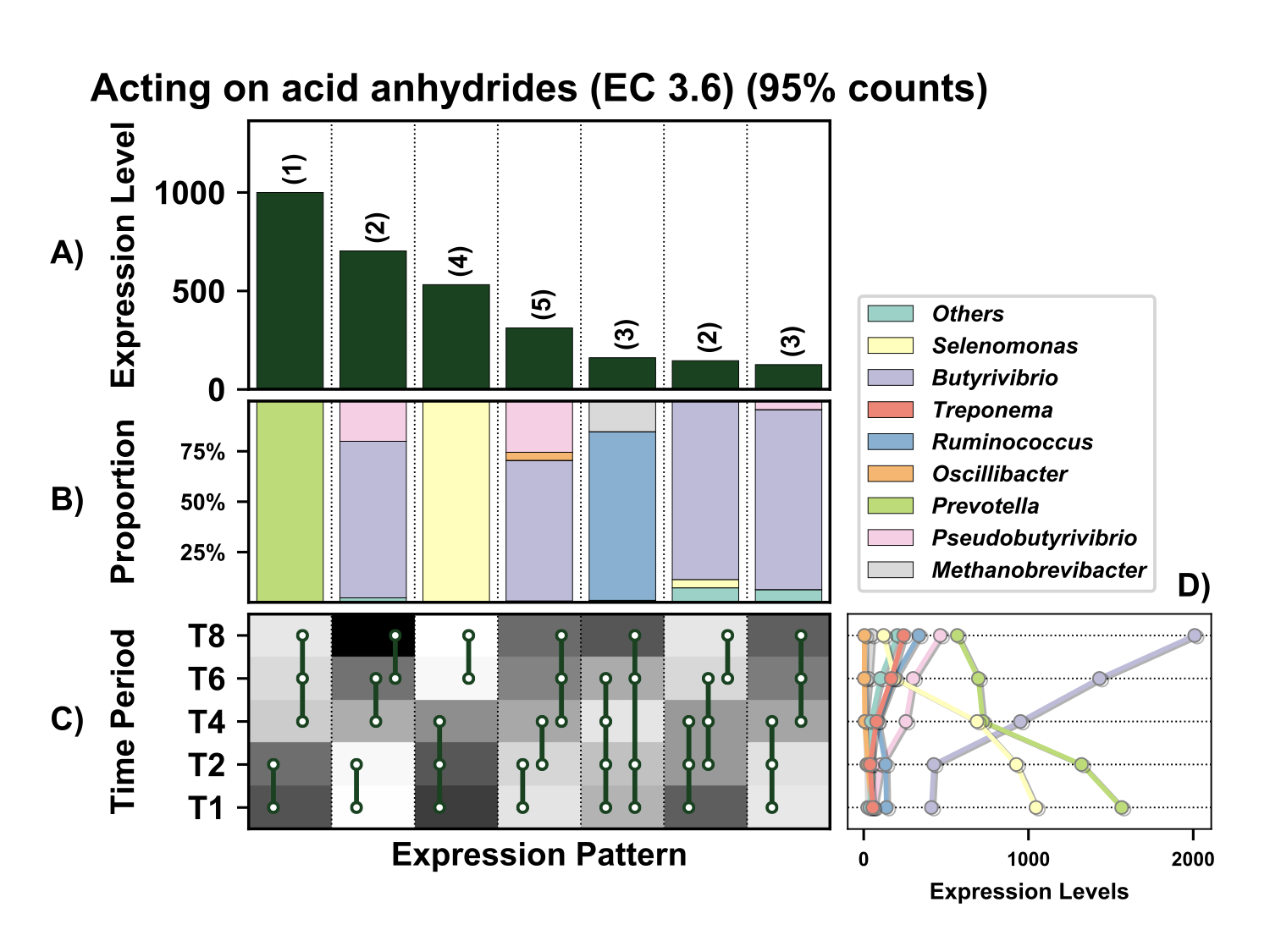

**Supplementary Figure 22.** Temporal expression of the top 95% most highly expressed genes acting on acid anhydrides (EC 3.6) with a significant interaction with time expressed by prokaryotes attached to fresh perennial ryegrass incubated *in situ* within the rumen. Each column represents a set of genes that showed the same differential expression (DE) pattern (denoted as expression pattern on x axis). A) Summed expression of all genes with the same DE pattern, in brackets the number of genes with the same DE pattern. B) The proportion of taxonomic genera contributing to the expression level for each DE pattern. C) Visual representation of the DE patterns for each set of genes across the timepoints sampled; The heatmap represents the level of expression for each timepoint (Low = White, High = Black); The lines and dots represent the specific DE pattern shared by all genes in this set where the timepoints connected by line and dots were not significantly different from each other. D) The level of expression of genes across each timepoint.

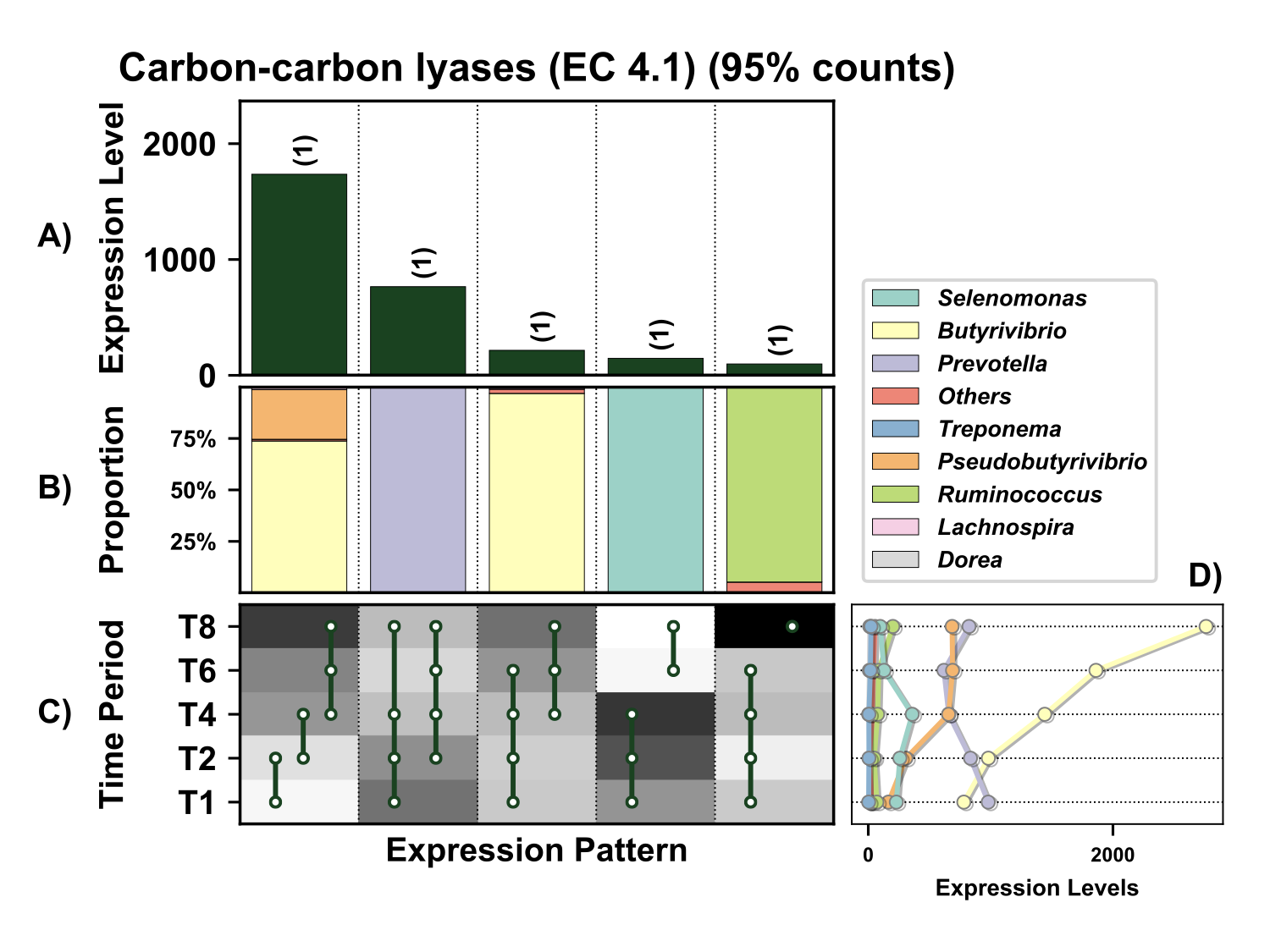

**Supplementary Figure 23.** Temporal expression of the top 95% most highly expressed genes acting on the carbon-carbon [lyases](https://en.wikipedia.org/wiki/Lyase) (EC 4.1) with a significant interaction with time expressed by prokaryotes attached to fresh perennial ryegrass incubated *in situ* within the rumen. Each column represents a set of genes that showed the same differential expression (DE) pattern (denoted as expression pattern on x axis). A) Summed expression of all genes with the same DE pattern, in brackets the number of genes with the same DE pattern. B) The proportion of taxonomic genera contributing to the expression level for each DE pattern. C) Visual representation of the DE patterns for each set of genes across the timepoints sampled; The heatmap represents the level of expression for each timepoint (Low = White, High = Black); The lines and dots represent the specific DE pattern shared by all genes in this set where the timepoints connected by line and dots were not significantly different from each other. D) The level of expression of genes across each timepoint.

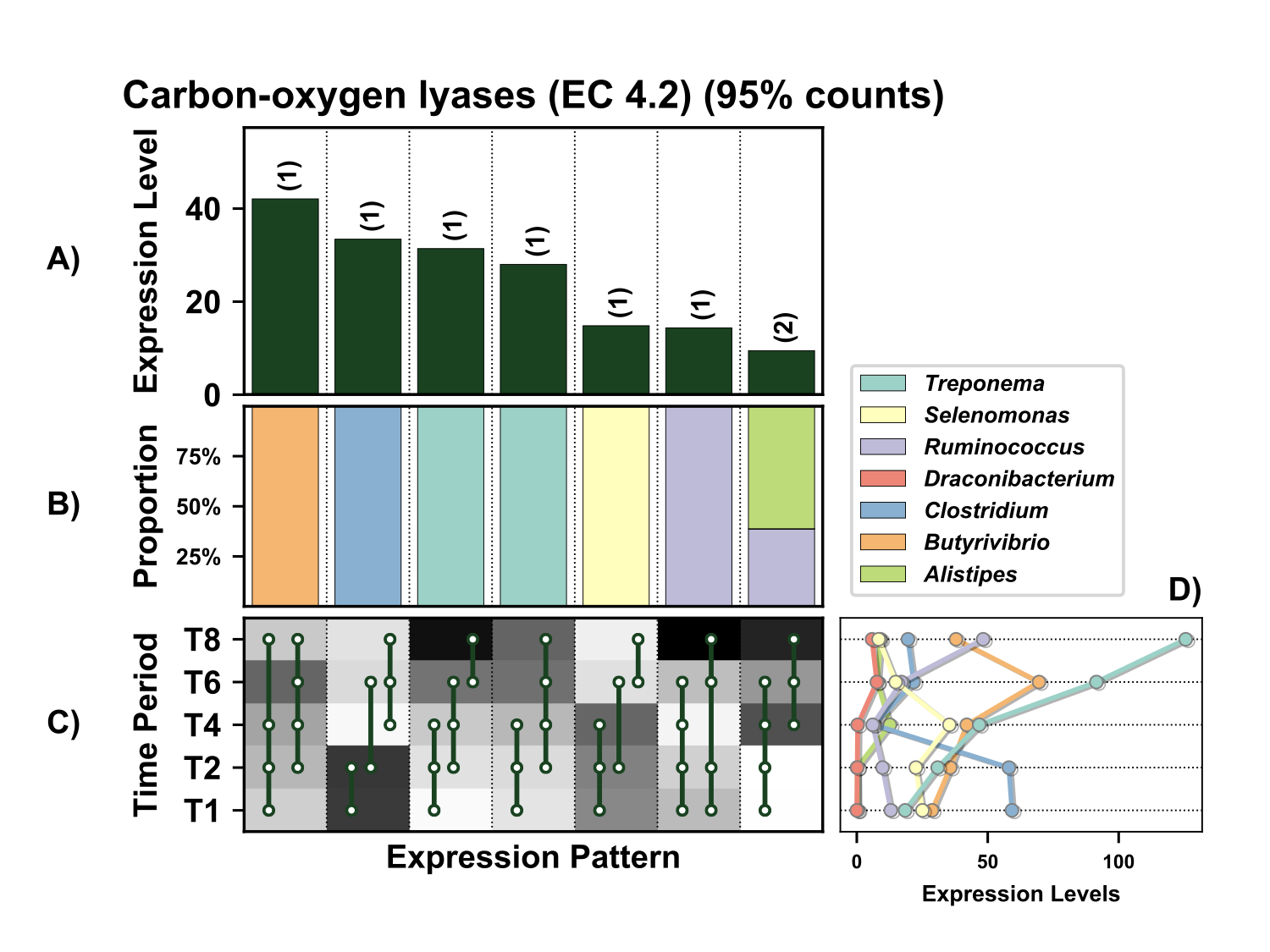

**Supplementary Figure 24.** Temporal expression of the top 95% most highly expressed carbon-oxygen [lyases](https://en.wikipedia.org/wiki/Lyase) (EC 4.2) with a significant interaction with time expressed by prokaryotes attached to fresh perennial ryegrass incubated *in situ* within the rumen. Each column represents a set of genes that showed the same differential expression (DE) pattern (denoted as expression pattern on x axis). A) Summed expression of all genes with the same DE pattern, in brackets the number of genes with the same DE pattern. B) The proportion of taxonomic genera contributing to the expression level for each DE pattern. C) Visual representation of the DE patterns for each set of genes across the timepoints sampled; The heatmap represents the level of expression for each timepoint (Low = White, High = Black); The lines and dots represent the specific DE pattern shared by all genes in this set where the timepoints connected by line and dots were not significantly different from each other. D) The level of expression of genes across each timepoint.

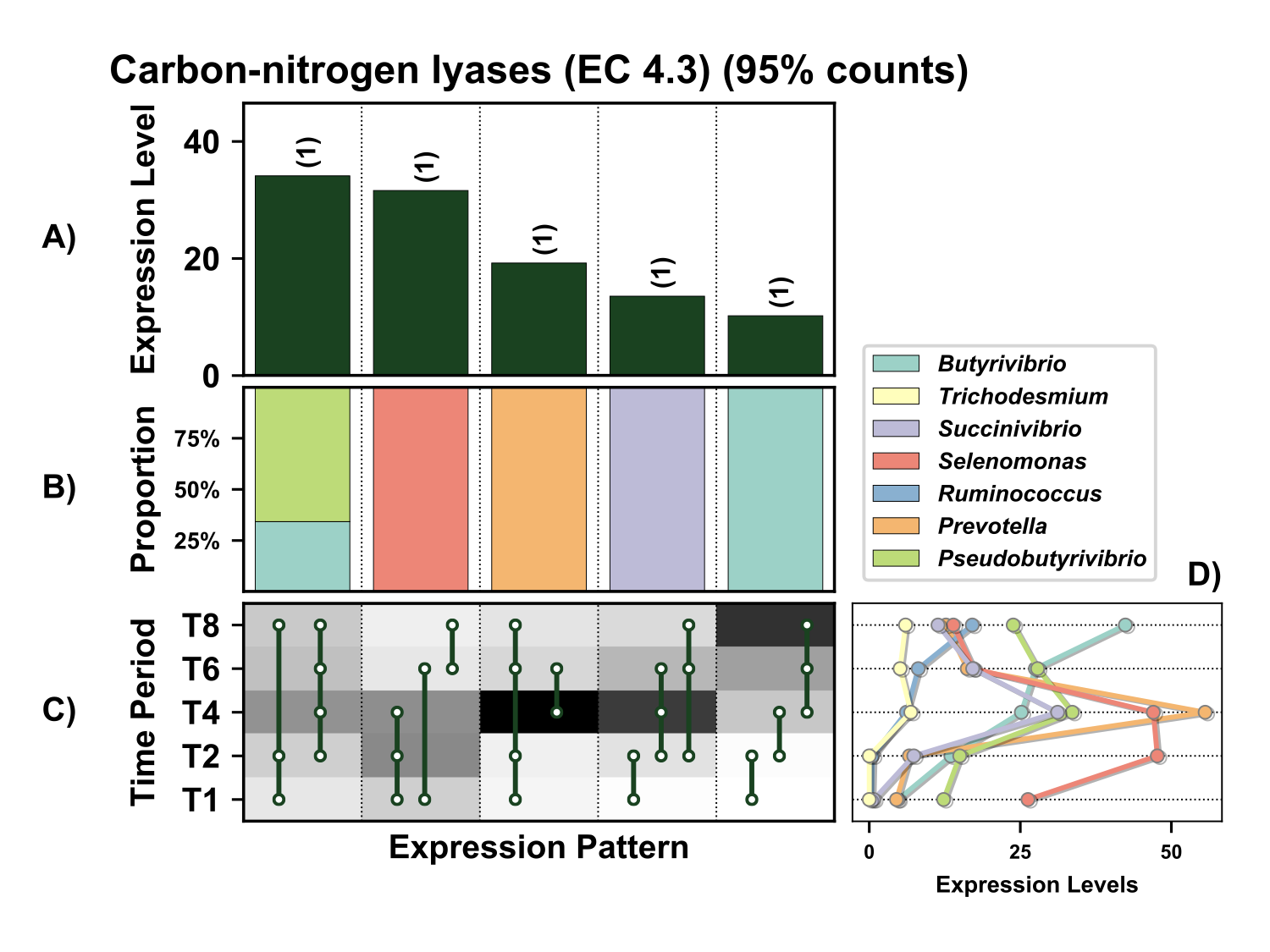

**Supplementary Figure 25.** Temporal expression of the top 95% most highly expressed carbon-nitrogen lyases (EC 4.3) with a significant interaction with time expressed by prokaryotes attached to fresh perennial ryegrass incubated *in situ* within the rumen. Each column represents a set of genes that showed the same differential expression (DE) pattern (denoted as expression pattern on x axis). A) Summed expression of all genes with the same DE pattern, in brackets the number of genes with the same DE pattern. B) The proportion of taxonomic genera contributing to the expression level for each DE pattern. C) Visual representation of the DE patterns for each set of genes across the timepoints sampled; The heatmap represents the level of expression for each timepoint (Low = White, High = Black); The lines and dots represent the specific DE pattern shared by all genes in this set where the timepoints connected by line and dots were not significantly different from each other. D) The level of expression of genes across each timepoint.

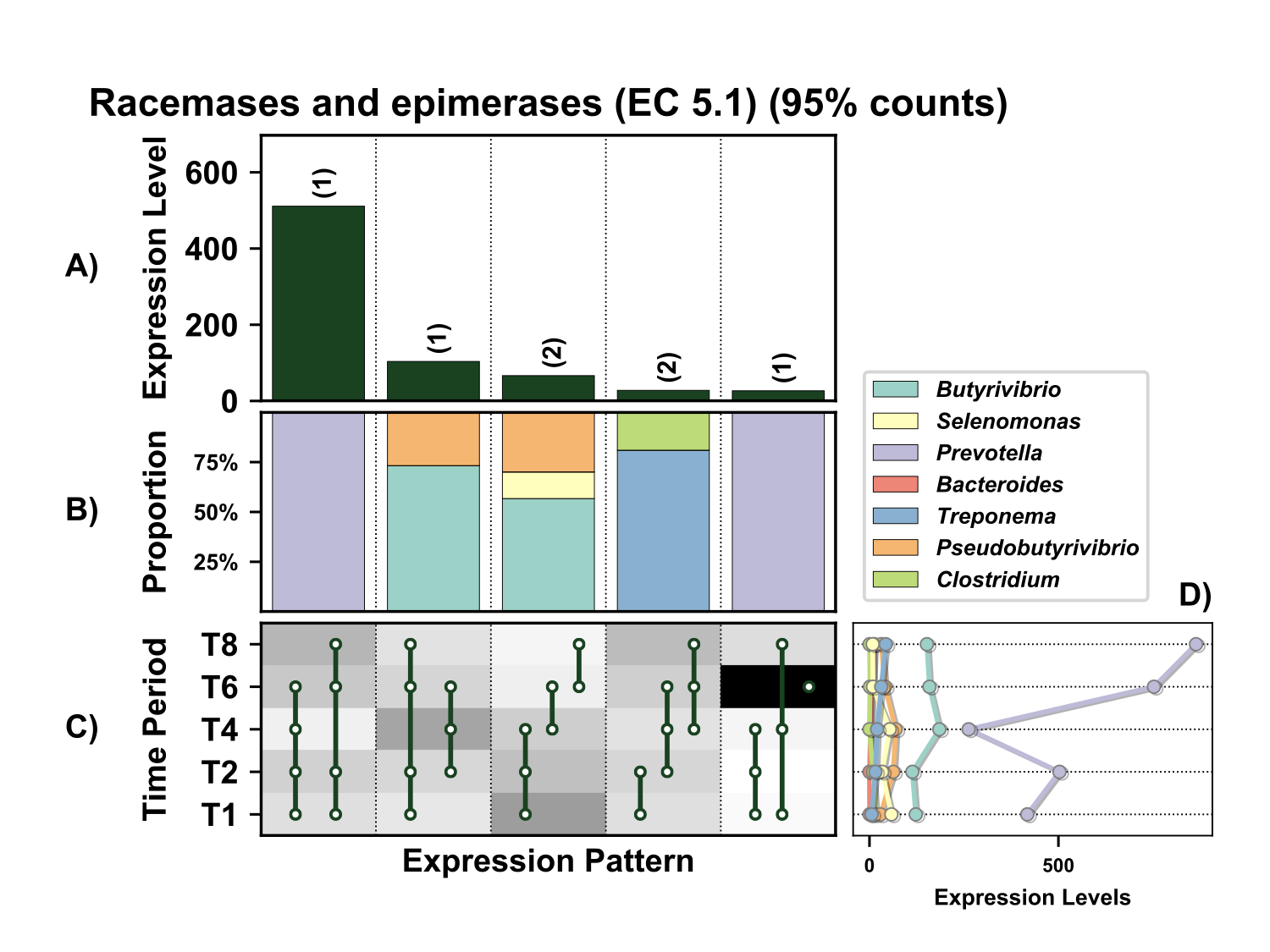

**Supplementary Figure 26.** Temporal expression of the top 95% most highly expressed racemases and epimerases (EC 5.1) with a significant interaction with time expressed by prokaryotes attached to fresh perennial ryegrass incubated *in situ* within the rumen. Each column represents a set of genes that showed the same differential expression (DE) pattern (denoted as expression pattern on x axis). A) Summed expression of all genes with the same DE pattern, in brackets the number of genes with the same DE pattern. B) The proportion of taxonomic genera contributing to the expression level for each DE pattern. C) Visual representation of the DE patterns for each set of genes across the timepoints sampled; The heatmap represents the level of expression for each timepoint (Low = White, High = Black); The lines and dots represent the specific DE pattern shared by all genes in this set where the timepoints connected by line and dots were not significantly different from each other. D) The level of expression of genes across each timepoint.

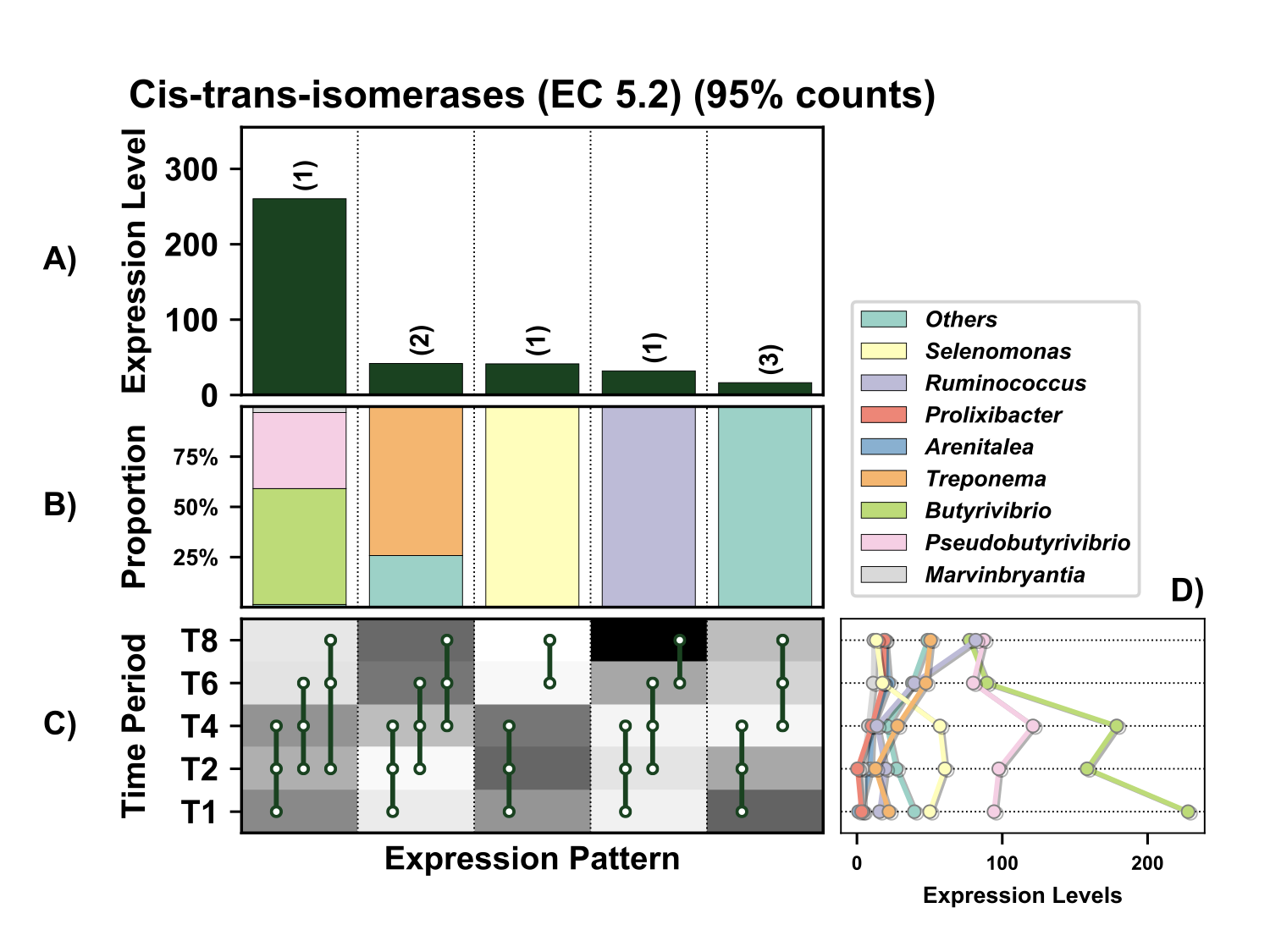

**Figure 27.** Temporal expression of the top 95% most highly expressed cis-trans-isomerase genes (EC 5.2) with a significant interaction with time expressed by prokaryotes attached to fresh perennial ryegrass incubated *in situ* within the rumen. Each column represents a set of genes that showed the same differential expression (DE) pattern (denoted as expression pattern on x axis). A) Summed expression of all genes with the same DE pattern, in brackets the number of genes with the same DE pattern. B) The proportion of taxonomic genera contributing to the expression level for each DE pattern. C) Visual representation of the DE patterns for each set of genes across the timepoints sampled; The heatmap represents the level of expression for each timepoint (Low = White, High = Black); The lines and dots represent the specific DE pattern shared by all genes in this set where the timepoints connected by line and dots were not significantly different from each other. D) The level of expression of genes across each timepoint.

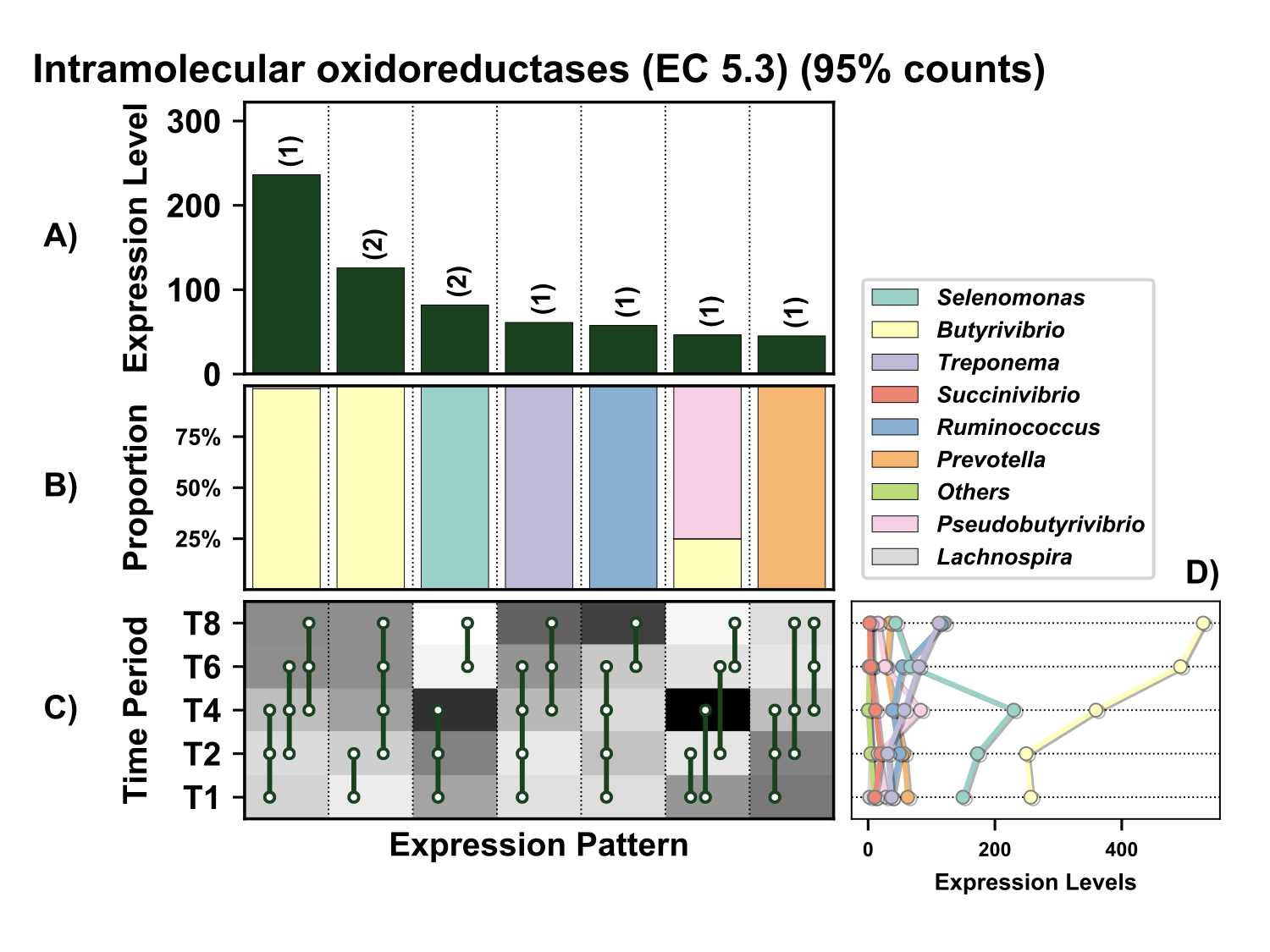

**Supplementary Figure 28.** Temporal expression of the top 95% most highly expressed intramolecular oxidoreductases (EC 5.3) with a significant interaction with time expressed by prokaryotes attached to fresh perennial ryegrass incubated *in situ* within the rumen. Each column represents a set of genes that showed the same differential expression (DE) pattern (denoted as expression pattern on x axis). A) Summed expression of all genes with the same DE pattern, in brackets the number of genes with the same DE pattern. B) The proportion of taxonomic genera contributing to the expression level for each DE pattern. C) Visual representation of the DE patterns for each set of genes across the timepoints sampled; The heatmap represents the level of expression for each timepoint (Low = White, High = Black); The lines and dots represent the specific DE pattern shared by all genes in this set where the timepoints connected by line and dots were not significantly different from each other. D) The level of expression of genes across each timepoint.

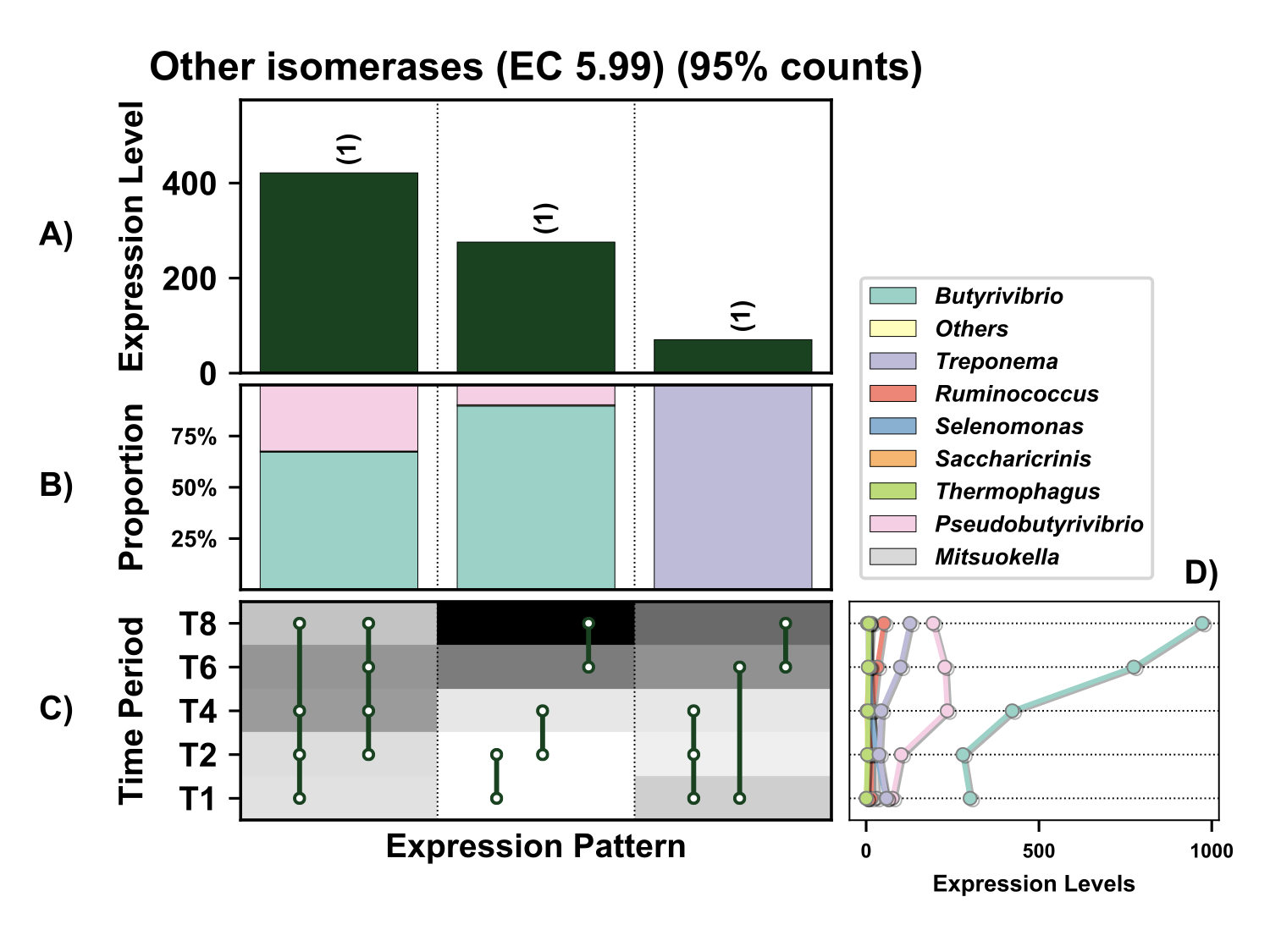

**Supplementary Figure 29.** Temporal expression of the top 95% most highly expressed other isomerases (EC 5.99) with a significant interaction with time expressed by prokaryotes attached to fresh perennial ryegrass incubated *in situ* within the rumen. Each column represents a set of genes that showed the same differential expression (DE) pattern (denoted as expression pattern on x axis). A) Summed expression of all genes with the same DE pattern, in brackets the number of genes with the same DE pattern. B) The proportion of taxonomic genera contributing to the expression level for each DE pattern. C) Visual representation of the DE patterns for each set of genes across the timepoints sampled; The heatmap represents the level of expression for each timepoint (Low = White, High = Black); The lines and dots represent the specific DE pattern shared by all genes in this set where the timepoints connected by line and dots were not significantly different from each other. D) The level of expression of genes across each timepoint.

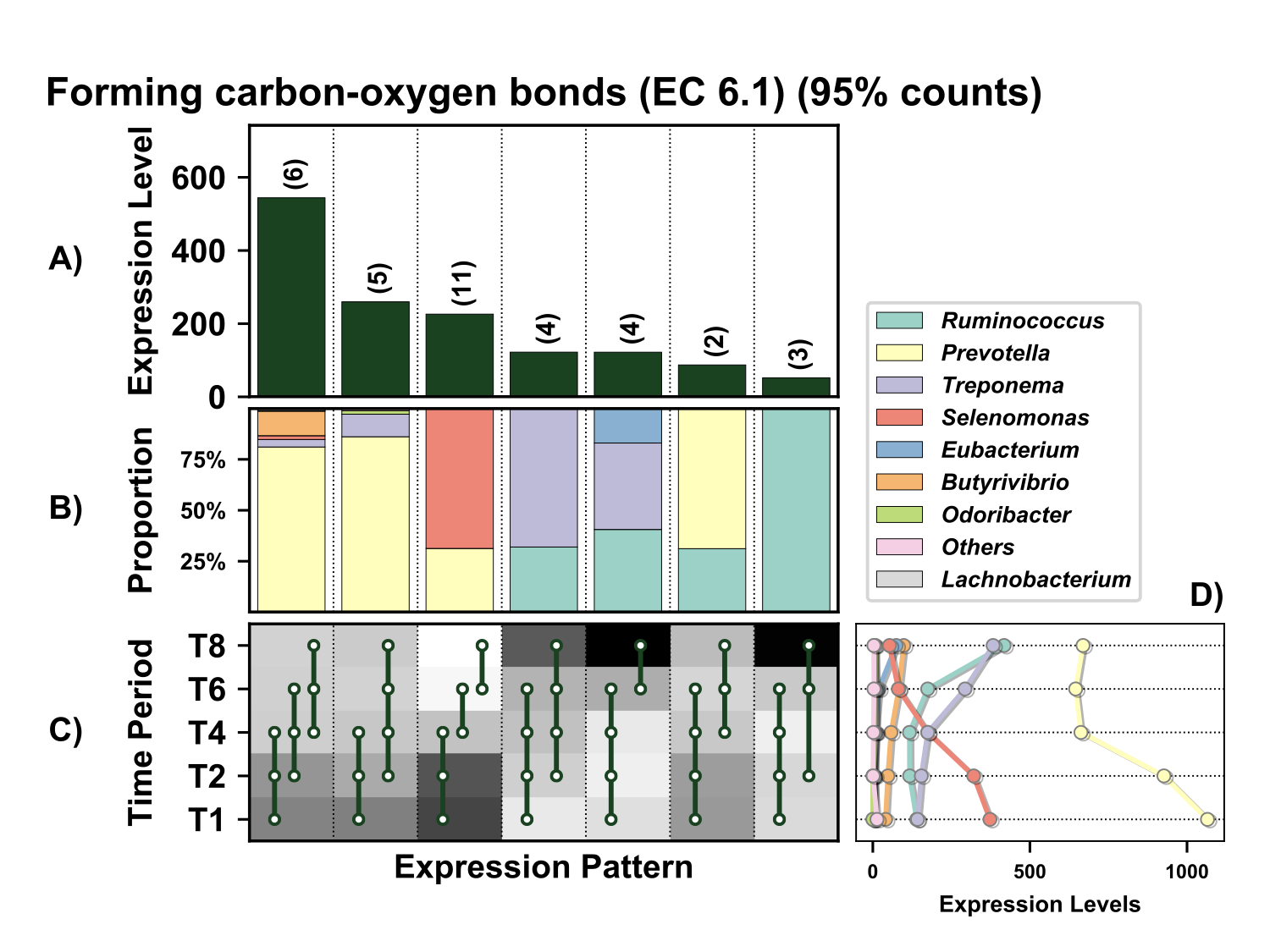

**Supplementary Figure 30.** Temporal expression of genes involved in formation of carbon-oxygen bonds (EC 6.1) within the top 95% of expressed by prokaryotes attached to fresh perennial ryegrass incubated *in situ* within the rumen. Each column represents a set of genes that showed the same differential expression (DE) pattern (denoted as expression pattern on x axis). A) Summed expression of all genes with the same DE pattern, in brackets the number of genes with the same DE pattern. B) The proportion of taxonomic genera contributing to the expression level for each DE pattern. C) Visual representation of the DE patterns for each set of genes across the timepoints sampled; The heatmap represents the level of expression for each timepoint (Low = White, High = Black); The lines and dots represent the specific DE pattern shared by all genes in this set where the timepoints connected by line and dots were not significantly different from each other. D) The level of expression of genes across each timepoint.

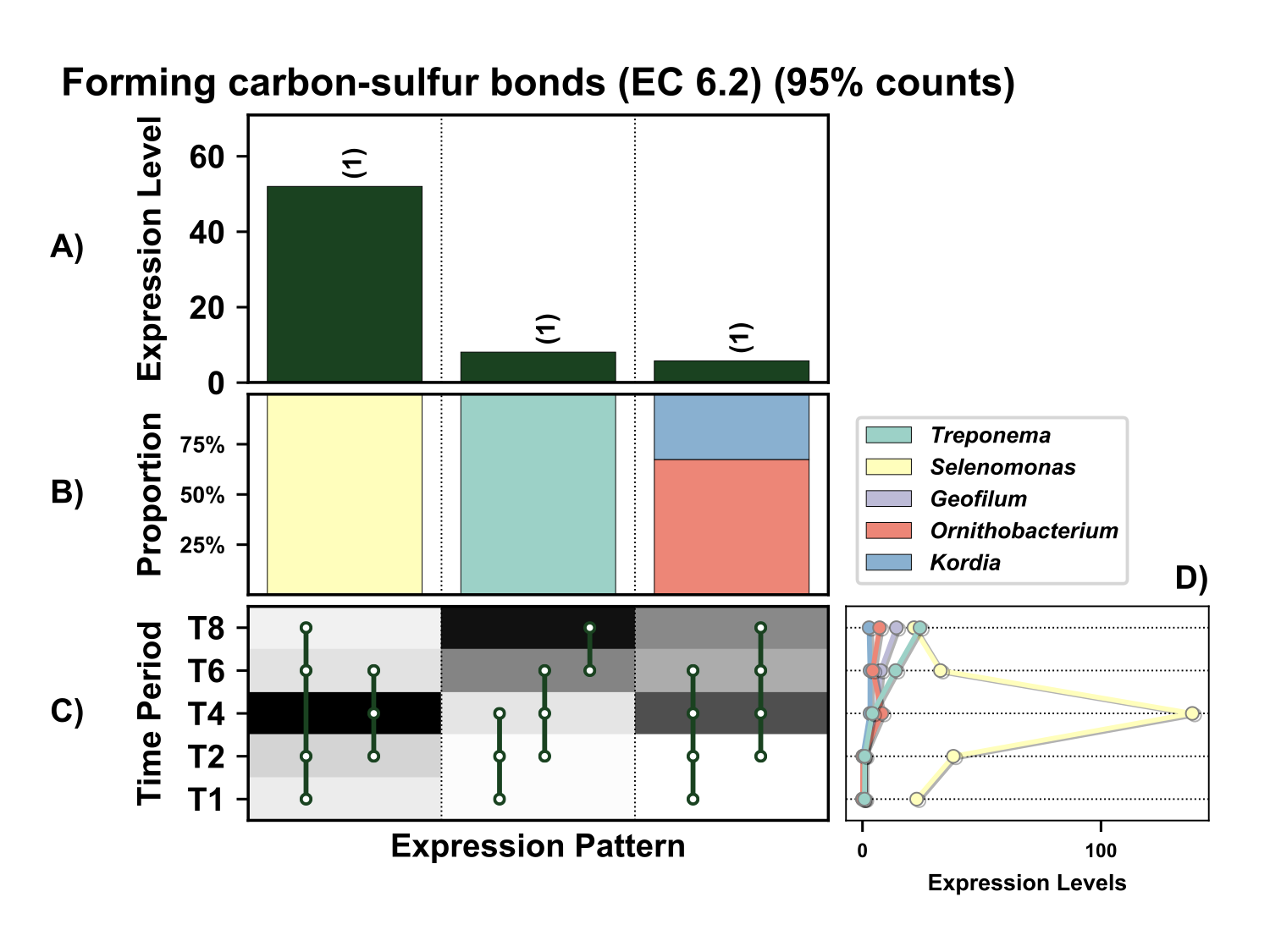

**Supplementary Figure 31.** Temporal expression of the top 95% most highly expressed genes involved in the formation of carbon-sulfur bonds (EC 6.2) with a significant interaction with time expressed by prokaryotes attached to fresh perennial ryegrass incubated *in situ* within the rumen. Each column represents a set of genes that showed the same differential expression (DE) pattern (denoted as expression pattern on x axis). A) Summed expression of all genes with the same DE pattern, in brackets the number of genes with the same DE pattern. B) The proportion of taxonomic genera contributing to the expression level for each DE pattern. C) Visual representation of the DE patterns for each set of genes across the timepoints sampled; The heatmap represents the level of expression for each timepoint (Low = White, High = Black); The lines and dots represent the specific DE pattern shared by all genes in this set where the timepoints connected by line and dots were not significantly different from each other. D) The level of expression of genes across each timepoint.

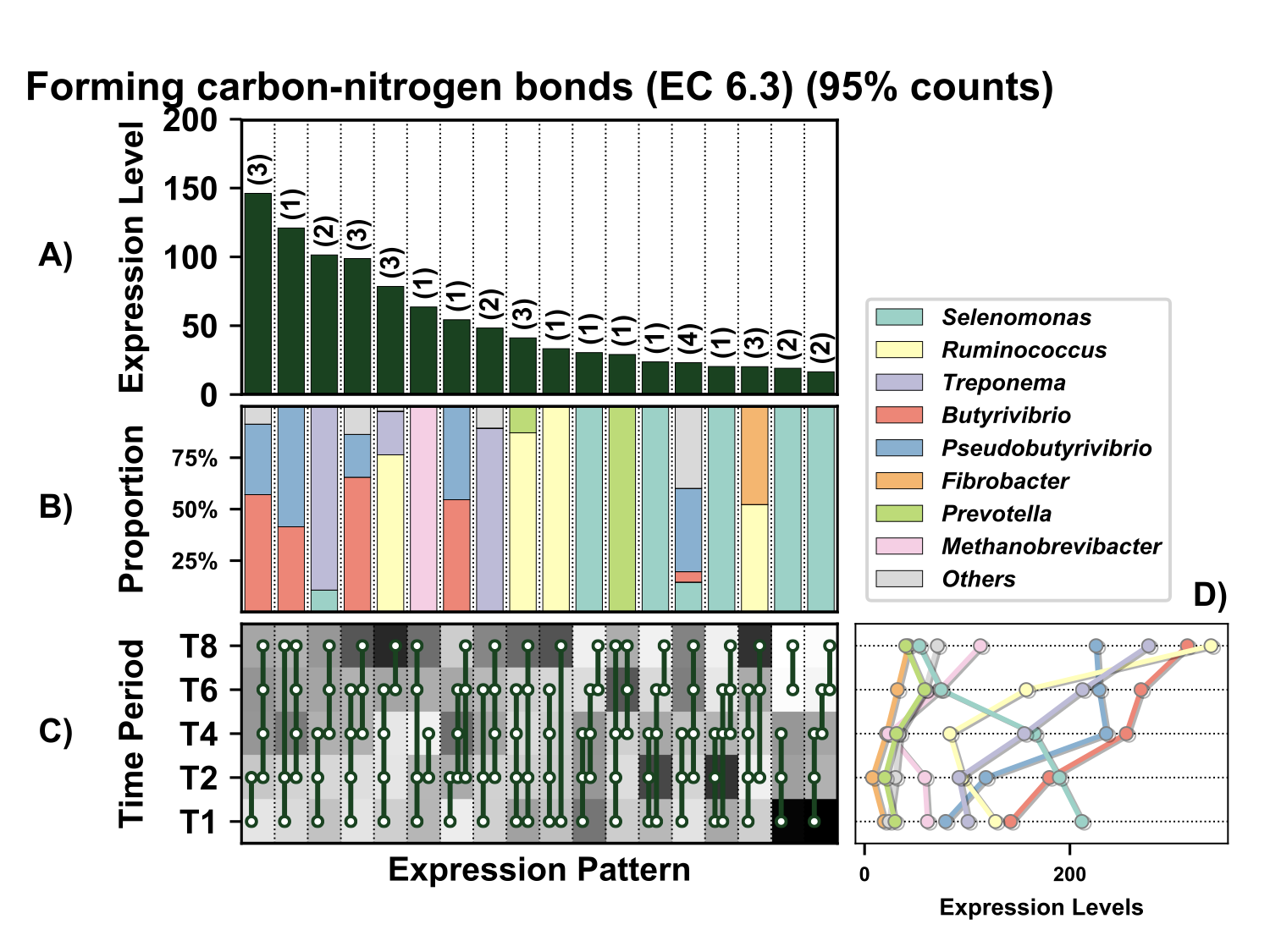

**Supplementary Figure 32.** Temporal expression of the top 95% most highly expressed genes involved in formation of carbon-nitrogen bonds (EC 6.3) with a significant interaction with time expressed by prokaryotes attached to fresh perennial ryegrass incubated *in situ* within the rumen. Each column represents a set of genes that showed the same differential expression (DE) pattern (denoted as expression pattern on x axis). A) Summed expression of all genes with the same DE pattern, in brackets the number of genes with the same DE pattern. B) The proportion of taxonomic genera contributing to the expression level for each DE pattern. C) Visual representation of the DE patterns for each set of genes across the timepoints sampled; The heatmap represents the level of expression for each timepoint (Low = White, High = Black); The lines and dots represent the specific DE pattern shared by all genes in this set where the timepoints connected by line and dots were not significantly different from each other. D) The level of expression of genes across each timepoint.

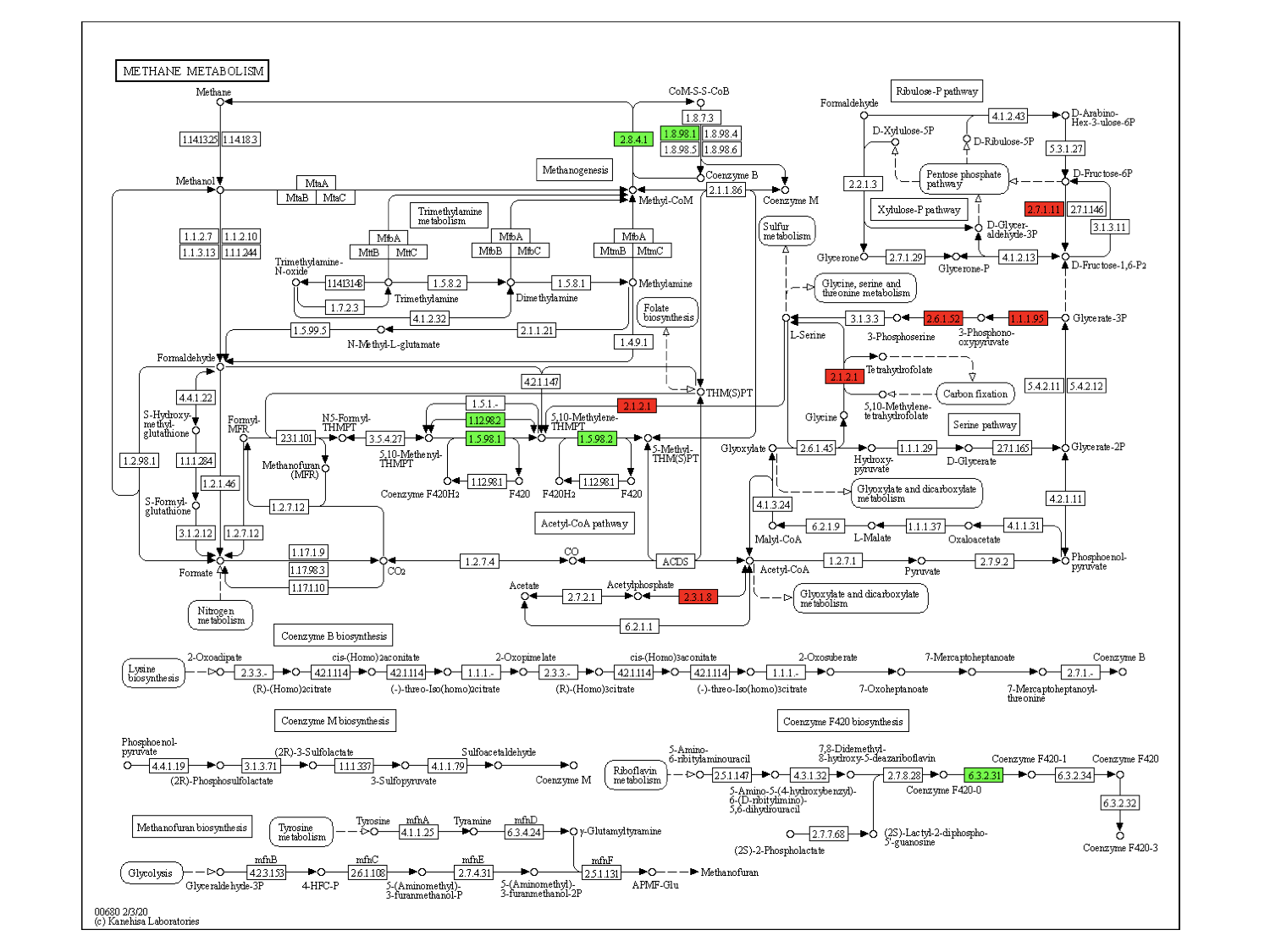

**Supplementary Figure 33.** Enzyme classifications involved in the KEGG methane metabolism pathway that contribute to data presented in Figure 3. The genera *Methanobrevibacter* (green) and *Butyrivibrio* (red) were dominated in terms of methane metabolism.

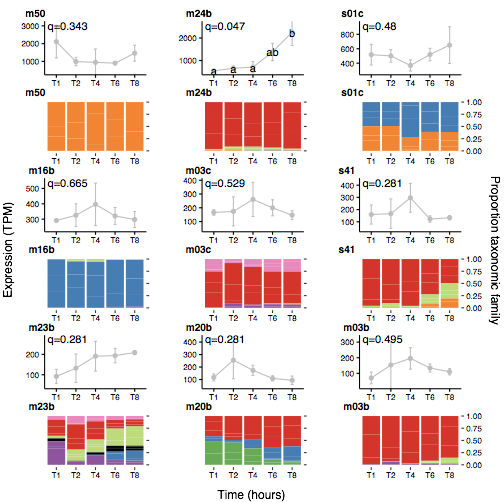

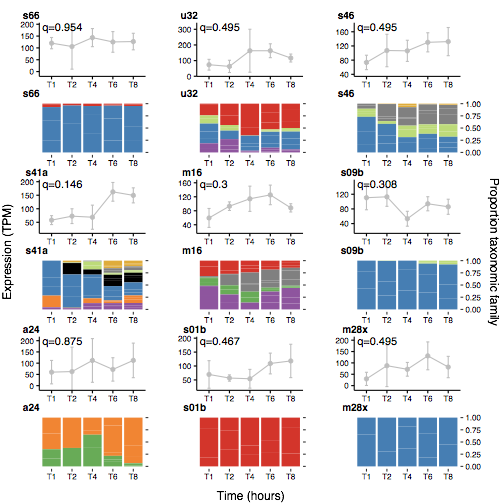

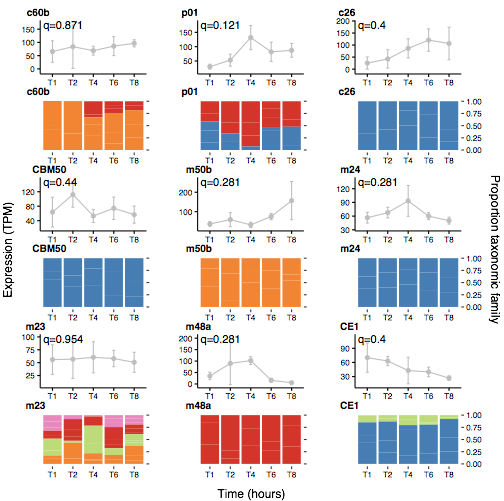

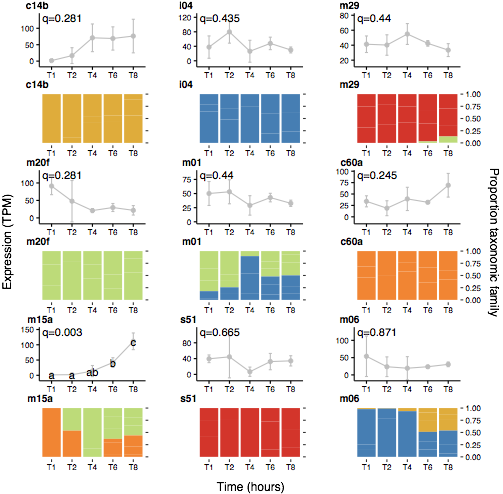

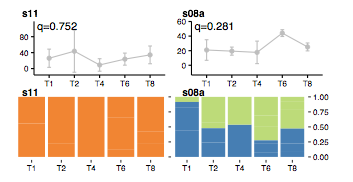

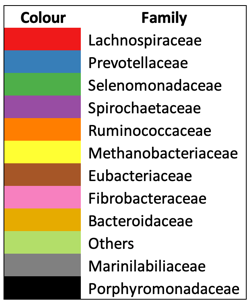

**Time(hours)**

**Supplementary Figure 34.** Temporal expression of all peptidases within the top 95% of expressed genes by perennial ryegrass attached bacteria within the rumen (Top figure) and their taxonomic origins (Figure below expression data).

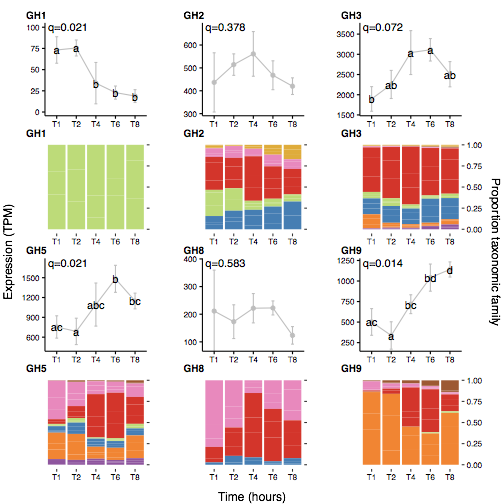

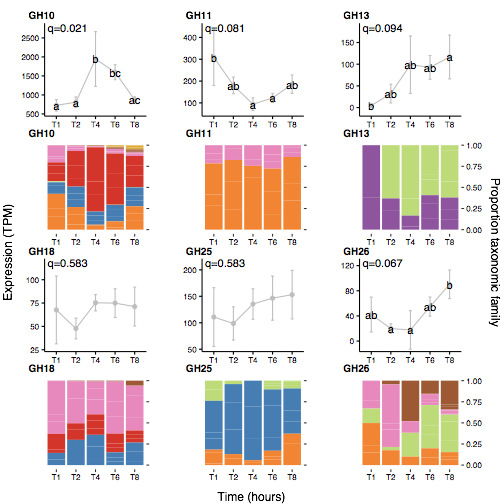

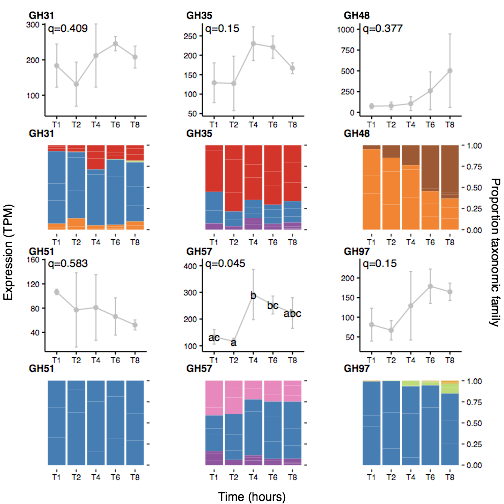

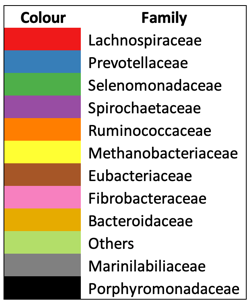

**Supplementary Figure 35.** Temporal expression of all carbohydrate active enzymes (CAZYmes also known as glycosyl hydrolases (GHs)) within the top 95% of expressed genes by perennial ryegrass attached bacteria within the rumen (Top figure) and their taxonomic origins (Figure below expression data).

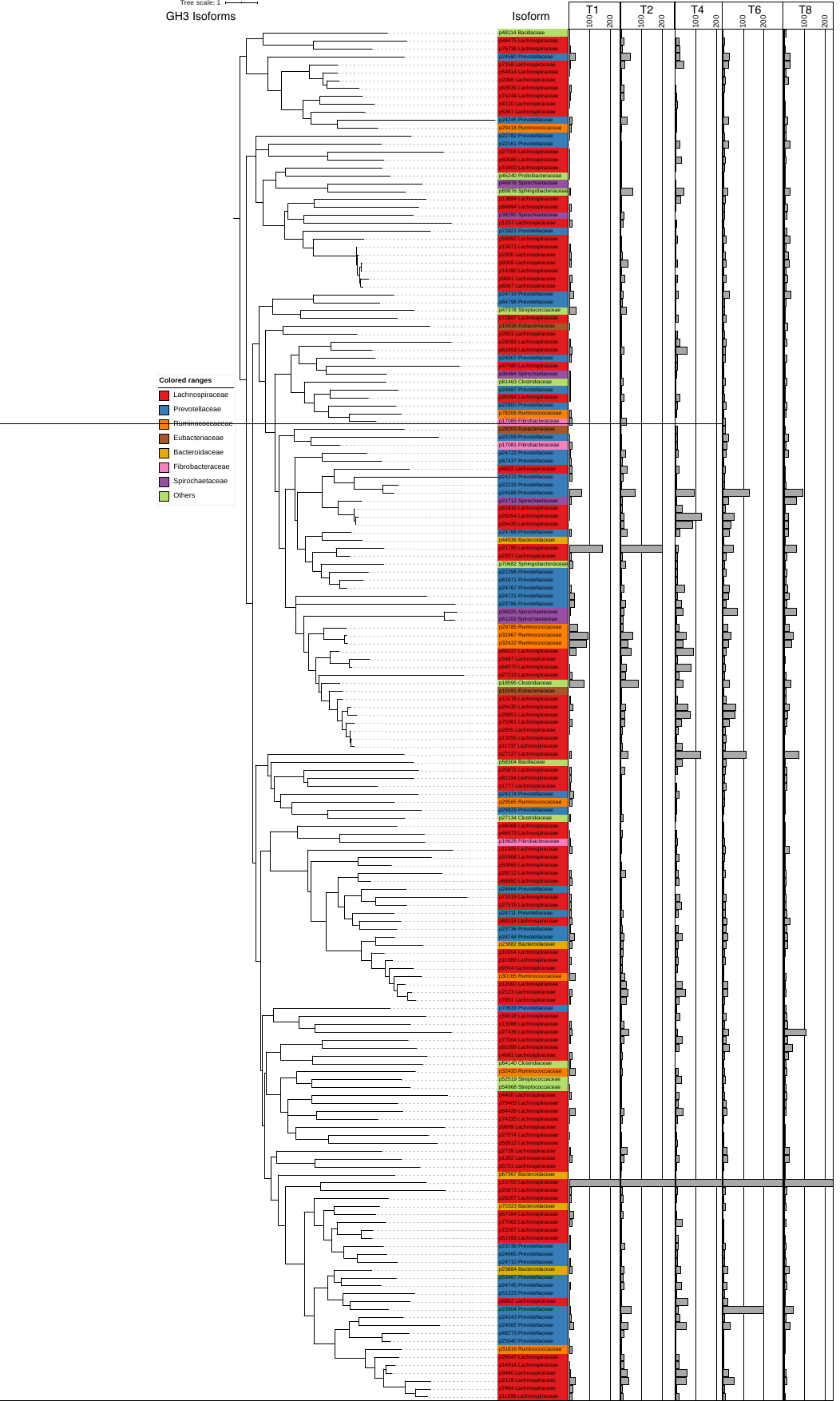

**Supplementary Figure 36.** Phylogenetic relationships and temporal expression of glycosyl hydrolase family 3 isoforms and their associated taxonomic origins.

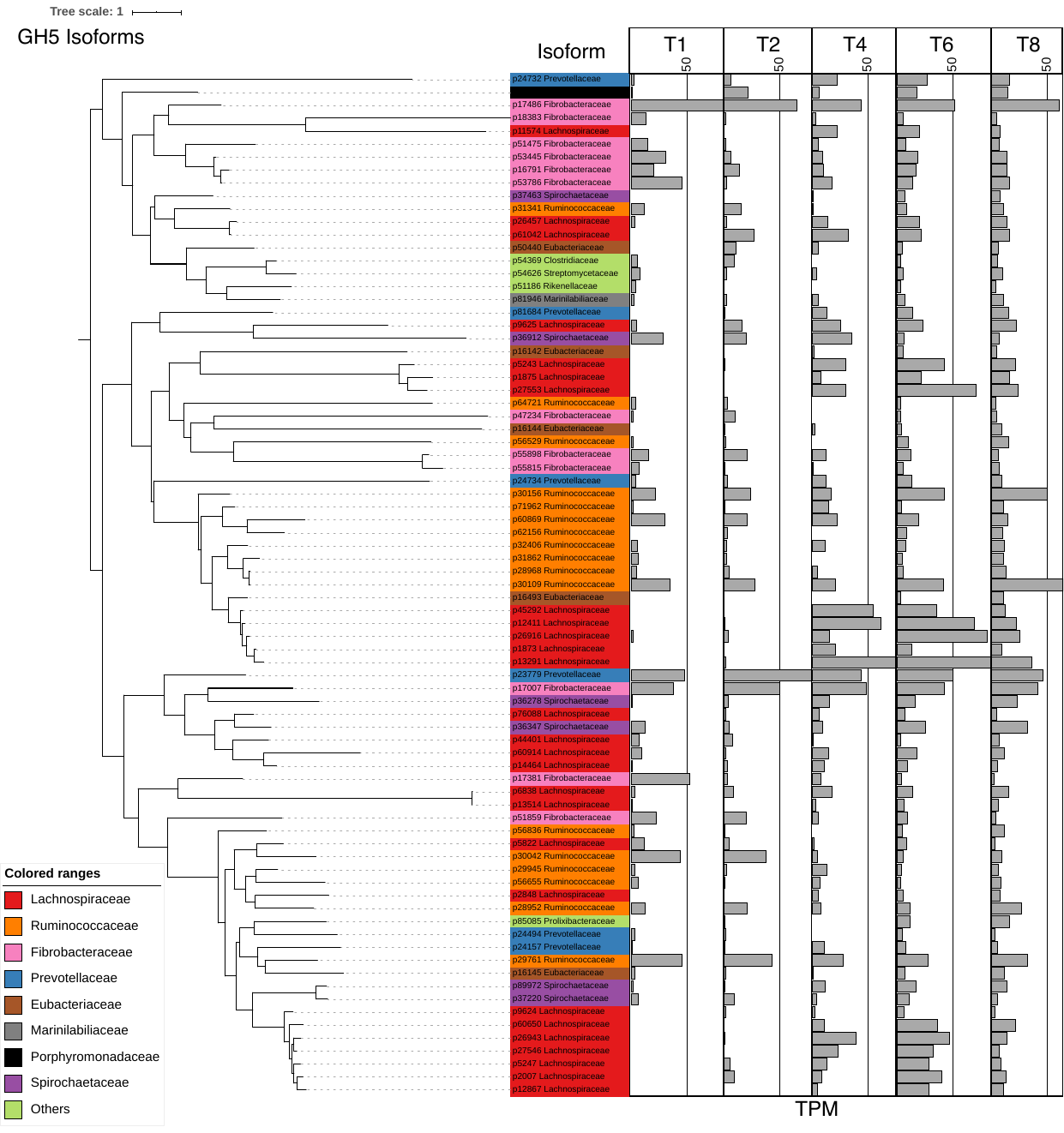

**Supplementary Figure 37.** Phylogenetic relationships and temporal expression of glycosyl hydrolase family 5 isoforms and their taxonomic origins.

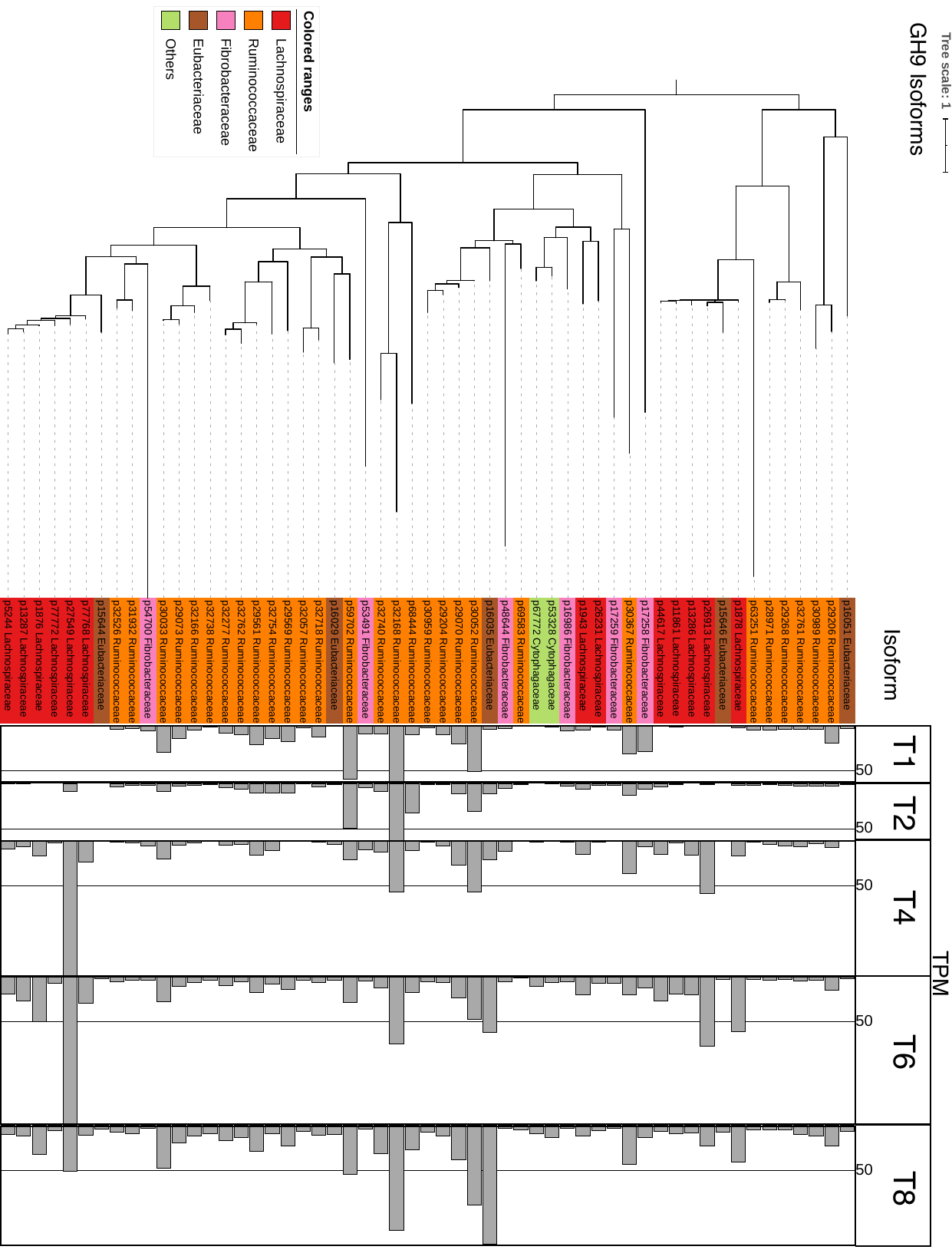

**Supplementary Figure 38.** Phylogenetic relationships and temporal expression of glycosyl hydrolase family 9 isoforms and their taxonomic origins.

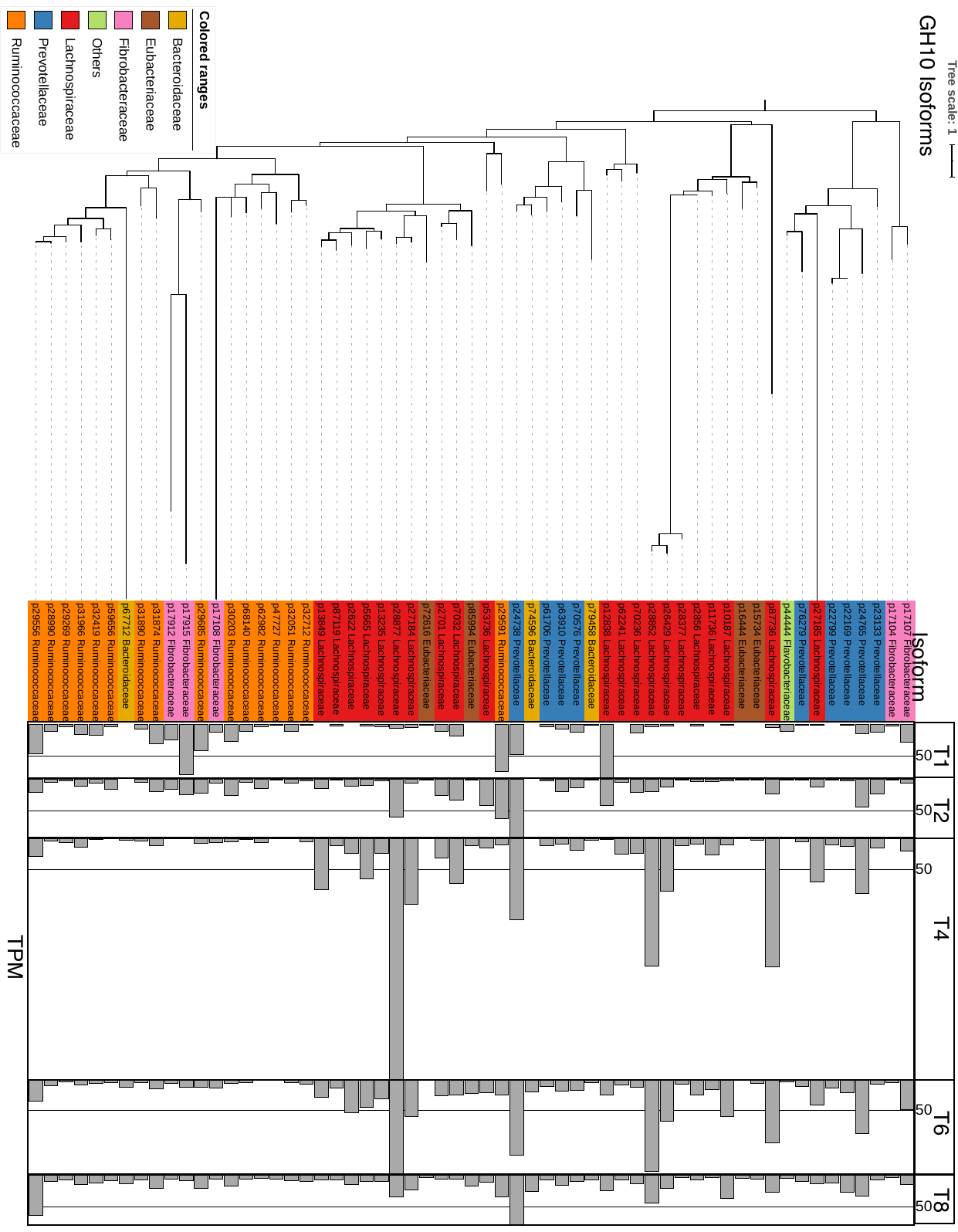

**Supplementary Figure 39.** Phylogenetic relationships and temporal expression of glycosyl hydrolase family 10 isoforms and their taxonomic origins.

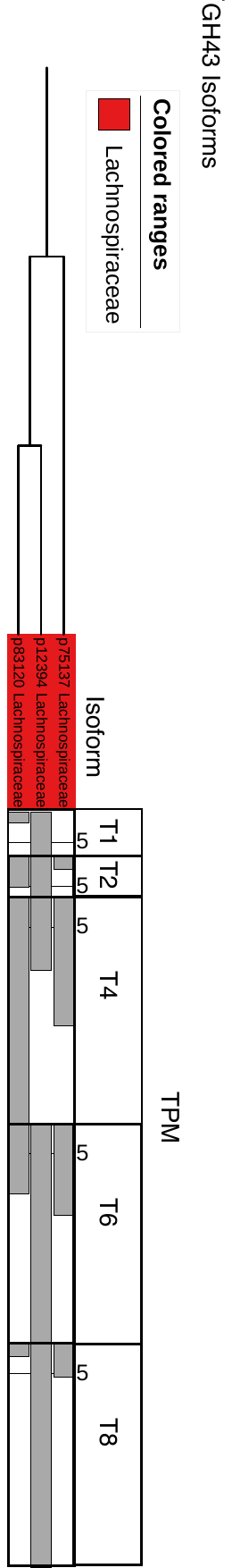

**Supplementary Figure 40.** Phylogenetic relationships and temporal expression of glycosyl hydrolase family 43 isoforms and their taxonomic origins.
